# Large structural variations in the haplotype-resolved African cassava genome

**DOI:** 10.1101/2021.06.25.450005

**Authors:** Ben N. Mansfeld, Adam Boyher, Jeffrey C. Berry, Mark Wilson, Shujun Ou, Seth Polydore, Todd P. Michael, Noah Fahlgren, Rebecca S. Bart

**Affiliations:** Donald Danforth Plant Science Center, St. Louis MO, USA 63132; Department of Ecology, Evolution, and Organismal Biology, Iowa State University, Ames, IA 50011; The Molecular and Cellular Biology Laboratory, The Salk Institute for Biological Studies, La Jolla, CA 92037, USA

**Keywords:** Cassava, Genome assembly, High heterozygosity, Haplotype phasing, Structural variants

## Abstract

Cassava (*Manihot esculenta* Crantz, 2n=36) is a global food security crop. Cassava has a highly heterozygous genome, high genetic load, and genotype-dependent asynchronous flowering. It is typically propagated by stem cuttings and any genetic variation between haplotypes, including large structural variations, is preserved by such clonal propagation. Traditional genome assembly approaches generate a collapsed haplotype representation of the genome. In highly heterozygous plants, this results in artifacts and an oversimplification of heterozygous regions. We used a combination of Pacific Biosciences (PacBio), Illumina, and Hi-C to resolve each haplotype of the genome of a farmer-preferred cassava line, TME7 (Oko-iyawo). PacBio reads were assembled using the FALCON suite. Phase switch errors were corrected using FALCON-Phase and Hi-C read data. The ultra-long-range information from Hi-C sequencing was also used for scaffolding. Comparison of the two phases revealed more than 5,000 large haplotype-specific structural variants affecting over 8 Mb, including insertions and deletions spanning thousands of base pairs. The potential of these variants to affect allele specific expression was further explored. RNA-seq data from 11 different tissue types were mapped against the scaffolded haploid assembly and gene expression data are incorporated into our existing easy-to-use web-based interface to facilitate use by the broader plant science community. These two assemblies provide an excellent means to study the effects of heterozygosity, haplotype-specific structural variation, gene hemizygosity, and allele specific gene expression contributing to important agricultural traits and further our understanding of the genetics and domestication of cassava.

**Significance statement:** The cassava varieties grown by subsistence farmers in Africa largely differ from the inbred reference genome due to their highly heterozygous nature. We used multiple sequencing technologies to assemble and resolve both haplotypes in TME7, a farmer-preferred cassava line, enabling us to study the considerable haplotypic structural variation in this line.

## Introduction

Cassava *(Manihot esculenta* Crantz 2n=2x=36) is a globally important crop and is particularly critical for subsistence farmers in the developing world (Ceballos *et al*., 2004). As an outcrossing plant, cassava is considerably heterozygous with a high genetic load and, thus suffers from inbreeding depression (Rojas *et al*., 2009). This has hindered genetic improvement via breeding in cassava, and many agriculturally favorable lines are commonly clonally propagated, which maintains any heterozygosity in the germplasm (Aye, 2011; Ramu *et al*., 2017). Moreover, the heterozygous nature of the cassava genome and limitations in sequencing technologies have limited the ability to accurately sequence and assemble the genome (Chin *et al*., 2016). Due to this, a partially inbred cassava accession, AM560-2, was selected as the cassava reference genome (Prochnik *et al*., 2012). AM560-2 is the product of three generations of selfing of the Colombian cassava line MCol1505, and is 94% homozygous (Bredeson *et al*., 2016). The reference genome has been an asset to the cassava community for more than 10 years, but due to the homozygous nature of the genome it does not accurately represent lines grown in farmer’s fields.

The development of long-read and long-range sequencing technologies and recent advancement in assembly algorithms have strong implications for genome assembly of heterozygous plant and animal species. Such haplotype-resolved genome assemblies can be crucial to our comprehension of genetics in crops with strong inbreeding depression where generation of inbred lines is very difficult and not representative of the agriculturally grown plants. However even with these advances, assembling fully haplotype-phased genomes is difficult, especially when rates of heterozygosity are high (Michael and VanBuren, 2020). New genome assembly strategies now exist for separate assembly of homologous and homeologous chromosome copies, allowing for accurate phasing of haplotypes and polyploid genomes (Chin *et al*., 2016; Koren *et al*., 2018; Kronenberg *et al*., 2018). One such strategy uses sequence data from parental lines to discern the haplotype-specificity of offspring sequence reads prior to their assembly (Koren *et al*., 2018). However, this strategy requires access to the parental genotypes, which are unknown in many clonally propagated farmer-preferred cassava lines. Another novel approach utilizes single cell sequencing of gamete cells to gain insight into phasing information and haplotype assembly (Campoy *et al*., 2020). This “Gamete binning” approach was showcased in the heterozygous tree crop apricot (*Prunus armeniaca*), and while potentially a viable option for field grown cassava lines, it requires extraction of pollen nuclei and other technical skills that are potentially limiting factors to its immediate adoption (Campoy *et al*., 2020). An alternate computational approach, implemented in the FALCON-Phase algorithm, uses mapping information from long-range chromatin conformation capture (Hi-C) sequencing to correctly phase haplotype assembled sequences (Kronenberg *et al*., 2018). This *de novo* approach can be used to correct assembly phase switch errors, and accurately represent the chromosome from telomere to telomere (Kronenberg *et al*., 2018).

Recent attempts at assembling heterozygous farmer-preferred cassava lines have produced contiguous large assemblies (Kuon *et al*., 2019). These assemblies however are limited due to the lack of haplotypic separation; the primary assemblies include both haplotypes and thus contain many duplicated sequences (Kuon *et al*., 2019; Lyons *et al*., 2021). This has implications on the assembly size and scaffolding which can be severely impacted by these duplications (Guan *et al*., 2020). Sequence duplication can also cause problems for downstream analyses such as read mapping and gene annotation. Assessing the deduplication, completeness, and quality of heterozygous genomes thus plays a critical role in each assembly step, to ensure truly resolved haplotypic sequences (Rhie *et al*., 2020).

Here, we assemble a phased diploid assembly of the Nigerian cassava landrace (<u>T</u>ropical-*<u>M</u>anihot*-*esculenta*) TME7, also known as “Oko-iyawo”, a farmer-preferred line resistant to the cassava mosaic disease virus (Rabbi *et al*., 2014). By assembling and phasing the moderately sized (∼700 Mb) diploid cassava genome we have a unique opportunity to study haplotype-specific structural polymorphisms maintained for generations by clonal propagation. Elucidation of haplotype-specific structural variations in cassava will have direct implications for our understanding of these types of variations in other clonally propagated, heterozygous crops with larger genomes, including many tree fruit crops and other horticulturally important species. The two haplotype assemblies will also provide an excellent means to study the haplotype-specific structural variation, synteny, and allele-specific gene expression that contribute to important agricultural traits, furthering our understanding of the genetics and domestication of cassava. As breeding is difficult in a crop such as cassava, a better understanding of the haplotype-specific genetics will allow for more accurate, appropriate, and targeted gene editing to improve lines for agricultural purposes.

## Results and discussion

### Genome size and heterozygosity

Due to the significant differences between TME7 (a clonally propagated, heterozygous, farmer-preferred line grown in Africa) and AM560-2 (an inbred South American line) we opted to re-estimate the genome size of TME7 prior to assembly. Both flow cytometry and a k-mer based approach [GenomeScope (Vurture *et al*., 2017)], estimated the genome size to be within the range of 670-711 Mb (Figure 1). We settled on ∼700 Mb as a target haploid size for this assembly. This estimate is moderately lower than that estimated for the reference genome line AM560-2 (∼750Mb, Bredeson *et al*., 2016). Based on the k-mer analysis, the repeat content was estimated at roughly 61% of the estimated genome size and the two very distinct k-mer frequency peaks suggested a high level of heterozygosity (Figure 1B, Supplementary Figure 1). The GenomeScope model further estimated the heterozygosity of this cassava line to be ∼1.4%, or roughly one polymorphism every ∼70 bp (Figure 1B). This is slightly lower than other outcrossing clonally propagated crops such as pear (1.6%, Vurture *et al*., 2017), grape (1.6-1.7%, (Patel *et al*., 2018; Guan *et al*., 2020), as well as the closely related rubber tree (1.6%, Shi *et al*., 2019). Nonetheless, this level of estimated heterozygosity suggested that haplotype-resolved assembly approaches would be appropriate for assembly of the cassava genome.

**Figure 1.**
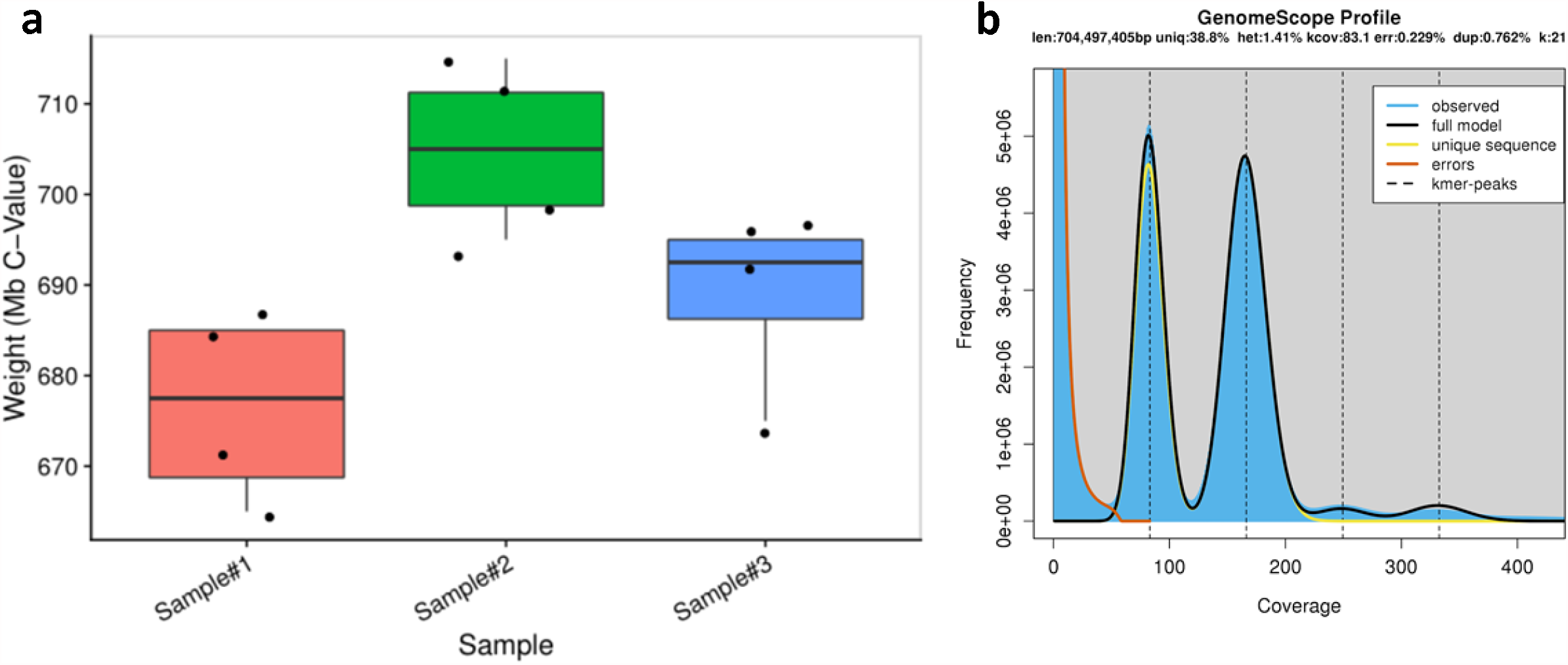
Estimates of TME7 genome parameters using flow cytometry and short reads. **(A)** Three biological samples of TME7, each with four technical replicates, were analyzed using flow cytometry. A mean genome size was estimated at 690 Mb. **(B)** Estimation of genome size, heterozygosity, and repetitiveness using GenomeScope Profile. K-mer size was set to 21, and k-mer coverage cutoff was set at 1e6 to include repeat regions in genome size estimates. The haploid genome size was estimated to be 704 Mb consisting of 61% repetitive sequence and a heterozygosity of 1.41%.

### Maximizing the diploid assembly

With that goal in mind, we sequenced the TME7 cassava genome using PacBio single-molecule long-read sequencing (SMRT) sequencing cells yielding roughly 90x coverage. We generated 64.2 Gb of data in 8,018,064 raw PacBio subreads (Supplementary Figure 2) that had an N50 of 11,099 bp; 4,970,318 of the reads were longer than 5,000 bp, which was used as a seed read size. We generated a PacBio-only assembly with FALCON and FALCON-Unzip (Chin *et al*., 2016). FALCON-Unzip assembled a total of 874 Mb in primary contigs, as well as an additional 157 Mb in haplotigs. FALCON-Unzip is limited in its ability to identify sequences with greater than 4-5% variation as haplotypic sequences, and these are often retained as primary contigs (Chin *et al*., 2016, also eg. Padgitt-Cobb *et al*., 2019). The total sequence assembled was ∼1 Gb, and while not yet well partitioned into haplotypes, included about 300 Mb in potentially haplotypic sequences. This represented the potential for an approximately 50% “unzipped” genome assembly. Assembly statistics for each stage of assembly and phasing are reported in Table 1.

**Table 1:**
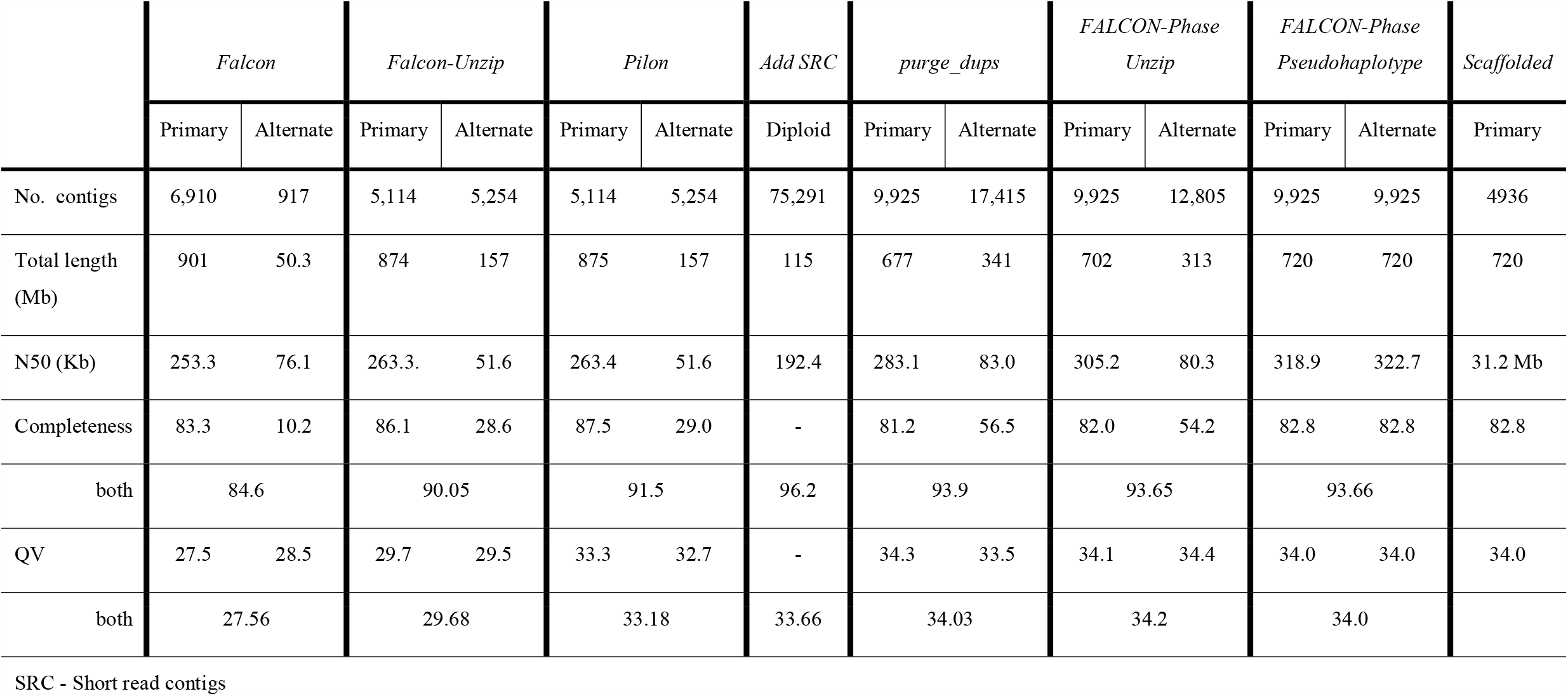
Assembly contiguity, completeness, and quality assessment

|  | *Falcon* |  | *Falcon-Unzip* |  | *Pilon* |  | *Add SRC* | *purge_dups* |  | *FALCON-Phase Unzip* |  | *FALCON-Phase Pseudohaplotype* |  | *Scaffolded* |
| --- | --- | --- | --- | --- | --- | --- | --- | --- | --- | --- | --- | --- | --- | --- |
|  | Primary | Alternate | Primary | Alternate | Primary | Alternate | Diploid | Primary | Alternate | Primary | Alternate | Primary | Alternate | Primary |
| No. contigs | 6,910 | 917 | 5,114 | 5,254 | 5,114 | 5,254 | 75,291 | 9,925 | 17,415 | 9,925 | 12,805 | 9,925 | 9,925 | 4936 |
| Total length (Mb) | 901 | 50.3 | 874 | 157 | 875 | 157 | 115 | 677 | 341 | 702 | 313 | 720 | 720 | 720 |
| N50 (Kb) | 253.3 | 76.1 | 263.3 | 51.6 | 263.4 | 51.6 | 192.4 | 283.1 | 83.0 | 305.2 | 80.3 | 318.9 | 322.7 | 31.2 Mb |
| Completeness | 83.3 | 10.2 | 86.1 | 28.6 | 87.5 | 29.0 | - | 81.2 | 56.5 | 82.0 | 54.2 | 82.8 | 82.8 | 82.8 |
| both | 84.6 |  | 90.05 |  | 91.5 |  | 96.2 | 93.9 |  | 93.65 |  | 93.66 |  |  |
| QV | 27.5 | 28.5 | 29.7 | 29.5 | 33.3 | 32.7 | - | 34.3 | 33.5 | 34.1 | 34.4 | 34.0 | 34.0 | 34.0 |
| both | 27.56 |  | 29.68 |  | 33.18 |  | 33.66 | 34.03 |  | 34.2 |  | 34.0 |  |  |

SRC - Short read contigs

**Figure 2.**
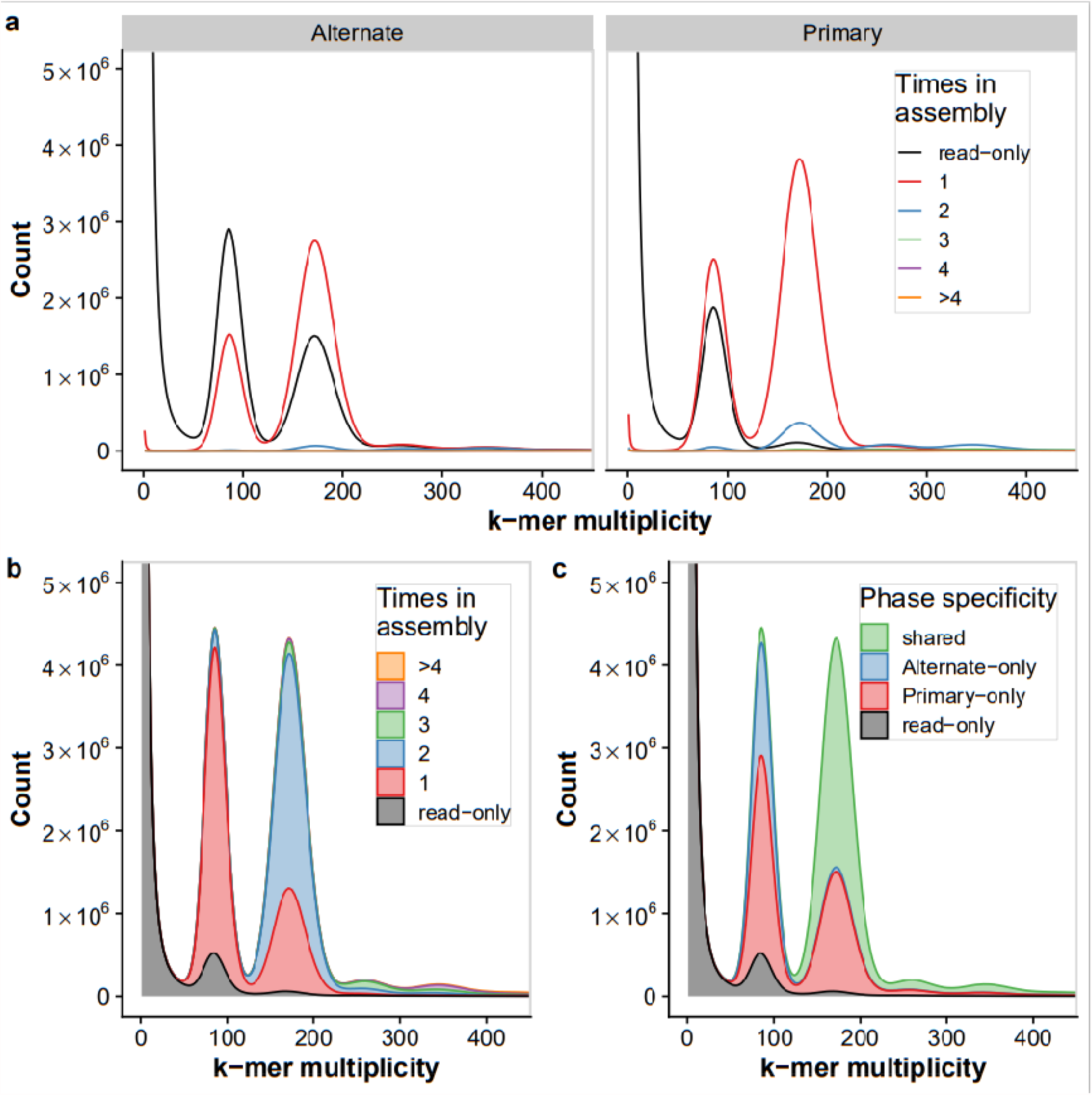
K-mer copy number and assembly analyses for the final phased TME7 assemblies. **(A)** K-mer count spectra for the alternate (haplotigs) and primary assemblies after phasing. **(B)** Diploid (primary + haplotigs) k-mer count spectra. In both **(A)** and **(B)**, short read k-mer distribution plots are colored by the number of times a k-mer is present in the assembly. K-mers denoted in grey are missing from the assembly and represent probable short read sequencing errors (k-mer multiplicity < 50) or missing assembled sequence (≥ 50). **(C)** Assembly spectra of the diploid assembly suggest that most homozygous k-mers (∼200x peak) are shared between the assemblies, while most of the heterozygous (∼100x peak) k-mers are phase specific.

To estimate the success of haplotypic separation and assembly quality we performed k-mer based analyses using Merqury (Rhie *et al*., 2020). Using raw, highly accurate short read sequencing representing data from both haploid sequences, k-mers which exist in 1- or 2-copies arise from heterozygous and homozygous regions, respectively. The k-mer distributions are then represented by the number of times each k-mer appears in the assembly allowing for the comparison of observed and expected coverage, estimation of reference-free completeness, and overall phasing success.

We first observed that even after polishing INDELs with pilon (Walker *et al*., 2014), a peak of heterozygous (1-copy) k-mers are missing from either the primary or alternate assemblies (Supplementary Figure 3). As our goal was to assemble a full heterozygous diploid phased assembly, we sought to maximize the amount of haplotypic sequence assembled. To this end we supplemented the long-read assembly with short read contigs containing additional heterozygous sequence. We identified k-mers that contained the short-reads pertaining to the missing heterozygous sequence and assembled them using SPAdes. (Bankevich *et al*., 2012). Some of these extra short read contigs (SRC) contained duplicates of already assembled sequences, but importantly many included the missing heterozygous sequence. Adding these SRCs to the full assembly brought the total assembled sequence to 1.15 Gb, or nearly the anticipated diploid size of ∼1.4 Gb. The number of missing k-mers was brought down from 23.7 M to 9.9 M using this approach and the “Completeness” score was brought up to 96.2% when including the SRCs (Table 1). Based on the missing k-mers, after adding the SRC an estimated 9.8 Mb of missing heterozygous sequence remained un-assembled.

**Figure 3.**
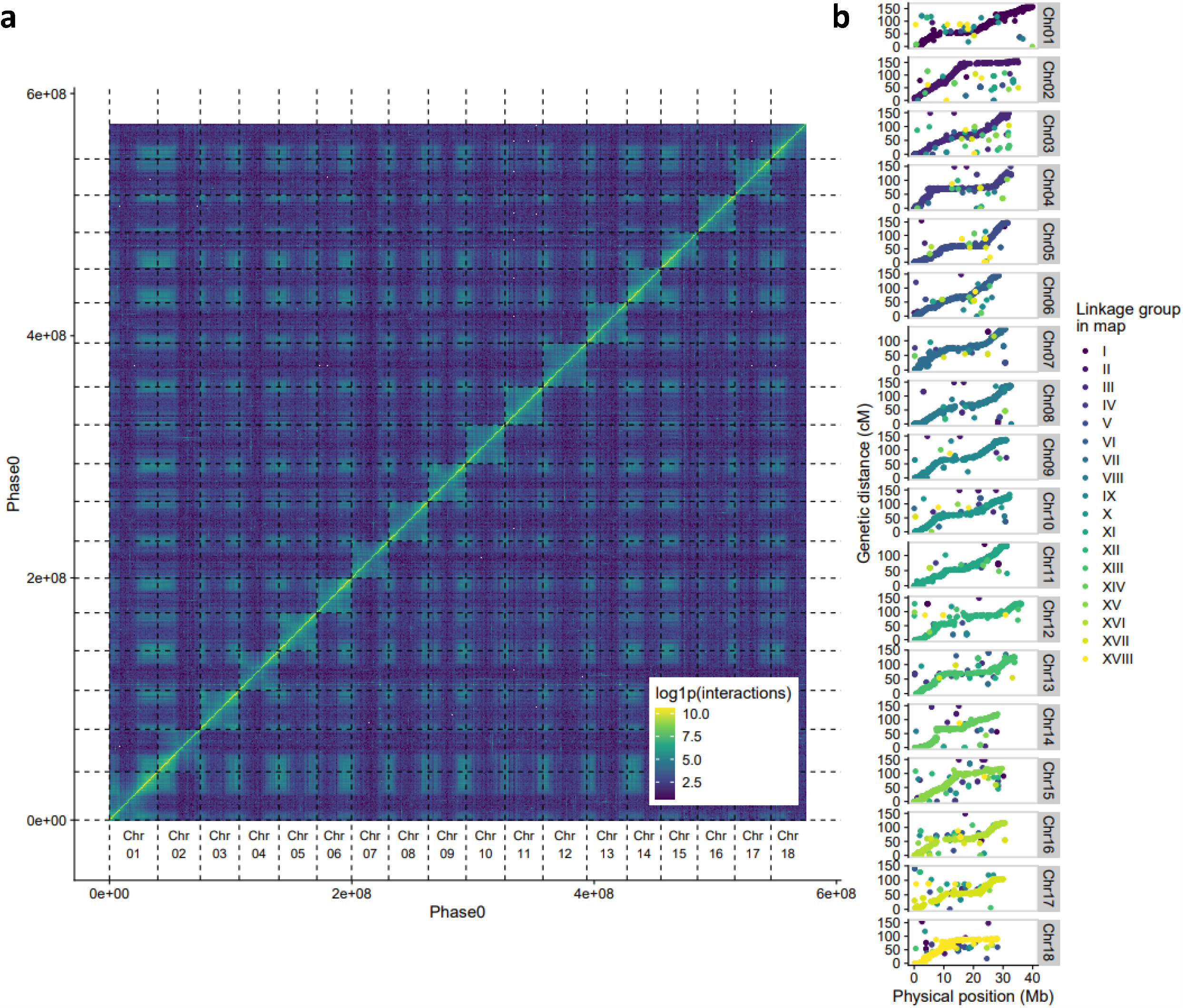
Validation of Hi-C scaffolding order and orientation by contact map and linkage map alignment. **(A)** Post-scaffolding Hi-C contact heatmap of the 18 largest scaffolds in the Phase0 assembly of TME7 showing the density of Hi-C interactions between regions of the genome. Color represents the intensity of interactions between regions, reported in log(1 + x). **(B)** Strong collinearity between the 22K marker Cassava Linkage Map and the TME7 Phase0 assembly. Markers are colored by their originating linkage group in the map.

### Haplotypic purging and deduplication

To complement the graph-based assembly approach used in FALCON-Unzip (Chin *et al*., 2016), other, orthogonal tools have since been developed to extract haplotypic sequences from primary assemblies (eg. Huang *et al*., 2017; Roach *et al*., 2018; Guan *et al*., 2020). These typically use read mapping coverage and sequence homology to identify potential haplotigs and “purge” them from the primary assembly (Roach *et al*., 2018). After maximizing our diploid assembly size to include as much haplotypic sequence as possible, our goal was to purge the primary assembly of haplotypic contigs, overlaps and sequence duplication, including those from our SRCs (Figure 2, Supplementary Figure 3). To this end we used purge_dups (Guan *et al*., 2020) which improves on the previous state-of-the-art, purge_haplotigs (Roach et al., 2018), by identifying and purging haplotypic overlaps. The final set of primary contigs included approximately 677 Mb assembled in 9925 contigs with an N50 of 283.1 kb. The resulting alternate assembly contained over 341 Mb assembled in haplotigs, representing a ∼50% “unzipped” genome.

K-mer spectra plots showed that the amount of sequence duplication was drastically reduced after purging, and that most of the heterozygous (1-copy) k-mers were now successfully separated into the two assemblies (Figure 2, Supplementary Figure 3). This was further confirmed by alignment of markers from the cassava 20k linkage map (ICGMC, 2015) (Figure 3B) and deduplication of BUSCO genes (Figure 4). After purging, the haplotig N50 size, which corresponds to the haplotype phase block, was 83 kb (90.5 kb if excluding SRC derived haplotigs). This is substantively smaller than the 7 Mb block described in the Arabidopsis F_1_ assembly by Chen and colleagues (2016), but is more similar to that observed in Carménère grape (89.5 Kb, Minio *et al*., 2019). Furthermore, it is consistent with relatively short dispersed regions of heterozygosity, and with the high rate of linkage disequilibrium decay described in cassava, an obligate out crosser (Ramu *et al*., 2017).

**Figure 4.**
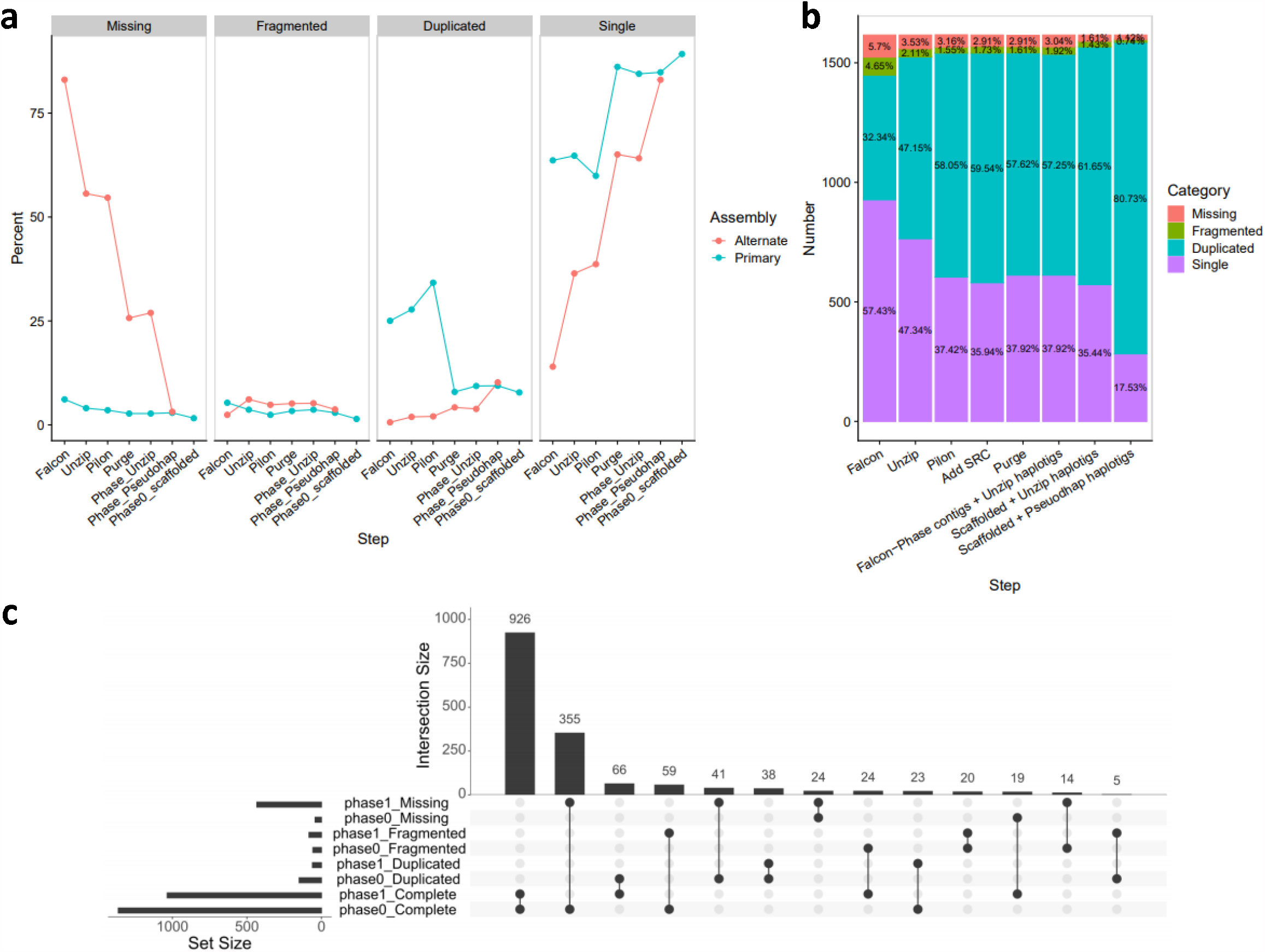
Summary of BUSCO analyses and phase specific BUSCOs. **(A)** The BUSCO scores for each step are reported for the **(A)** alternate or primary assemblies or **(B)** full (concatenated) assemblies. In **(B)**, after polishing with Pilon, short read contigs (SRC) were assembled and added to the full assembly, prior to haplotypic purging. **(C)** Overlap in BUSCO categories of the final FALCON-Phase Unzip-emit primary (Phase0) and alternate (Phase1) assemblies shows that most BUSCOs are phased and exist in both assemblies.

### Haplotype phasing and scaffolding with Hi-C sequencing

To get a more accurate representation of the TME7 pseudo-haplotypes, we phased the primary and haplotig assemblies using Hi-C data and FALCON-Phase (Kronenberg *et al*., 2018). We noticed however that during the placement and mincing stages of the algorithm, FALCON-Phase was discarding over 40 Mb of sequence from both primary and haplotig assemblies. We compared the haplotig truncation lengths with the contig vs. haplotig alignment lengths and identified that the FALCON-Phase *coords2hp*.*py* script truncated both contigs and haplotigs at the ends of alignments. We hypothesized that if large structural variations exist between haplotypes, this could affect how FALCON-Phase aligns and places haplotigs vs. their primary contigs. Due to these large structural variations between the haplotypes, haplotig sequences were truncated to exclude the non-aligning sequences. Merqury analysis showed that removal of these sequences reduced the number of heterozygous k-mers in the assembly (Supplementary Figure 4). We thus modified the *coords2hp*.*py* script in FALCON-Phase to force it to include the entire length of each haplotig, rather than only the length of the sequences that aligned.

The result was one complete set of 9,925 contigs comprising ∼720 Mb for each phase which included almost all the original heterozygosity assembled. This suggests we were able to successfully assemble nearly the entirety of the TME7 genome (720 Mb haploid assembly vs. ∼700 Mb estimated genome size) with a contig N50 of approximately 320 kb for both assemblies. When emitted in “unzip” format, the total primary and haplotig contig length was 702 (N50 = 305 kb) and 311 Mb (N50 = 80 kb), respectively. We assessed the success of the phasing step using Merqury (Figure 2). A modest increase in homozygous sequence duplication was observed after phasing, probably due to incorporation of homozygous contig boundaries into the primary assembly (Figure 2A). This minor sequence duplication in the “pseudohaplotype” assembly was also observed with the unmodified version of the *coords2hp*.*py* script suggesting it may be an inherent issue with the FALCON-Phase algorithm (Supplementary Figure 4). While this additional minor duplication is a limitation with this phase correction approach, the benefits of accurate phasing outweigh this issue.

After phasing, the Hi-C data was further used to scaffold the assembly into 18 chromosome length scaffolds. Contigs designated as part of Phase0 were scaffolded using the Proximo algorithm (Phase Genomics) and manual scaffolding curation with Juicebox (Rao *et al*., 2014; Durand *et al*., 2016). This process resulted in placing ∼80% of all sequence in a set of 18 chromosome-scale scaffolds containing 580 Mb of sequence (Figure 3A). We validated the scaffolding order and orientation by aligning 22,403 SNP markers from the cassava composite map (ICGMC, 2015) to both phases. After filtering for > 95% identity and >150 bp length, more than 19,000 markers aligned uniquely to both phases. We plotted the concordance between the new *de novo* assembly and the linkage map and observed high collinearity between the two (Figure 3B). Except for a few cases, there was high agreement between the physical and linkage maps (average Spearman’s correlation of 0.96). Approximately 1,900 marker sequence tags had duplicate mapping sites on the same scaffold in both phases and were distributed along the chromosome scaffolds (Supplementary Figure 5). While this may suggest potential sequence duplication or retained heterozygosity, an alternate explanation for some of these duplications are genotype specific duplications in TME7 that differ from the inbred reference genome. This represents a significant improvement over the previous attempts at assembly of heterozygous African cassava lines (Kuon *et al*., 2019), where close to 30% of markers had multiple map hits, indicating a not well deduplicated assembly.

**Figure 5.**
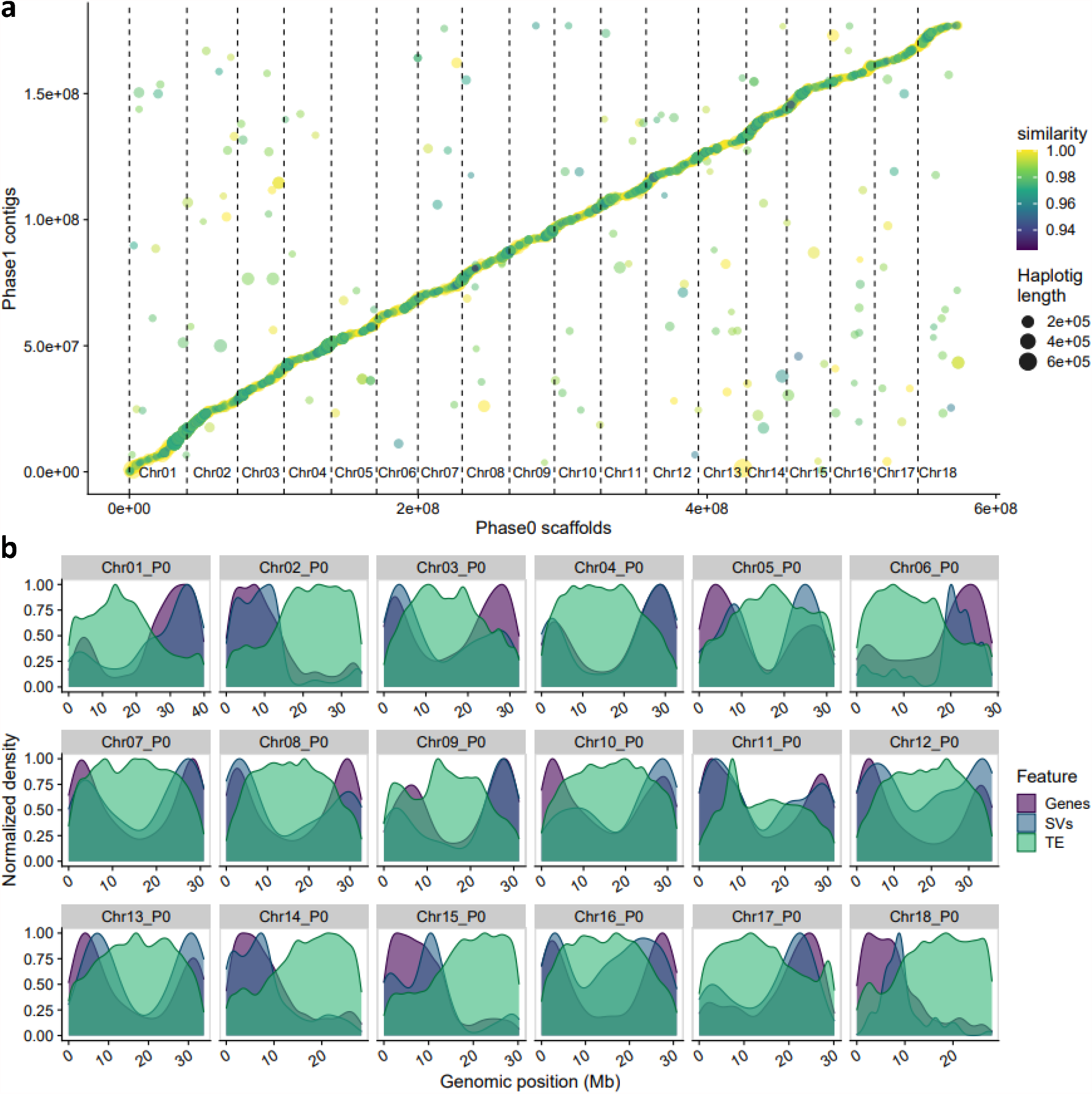
Comparison of the TME7 haplotype phased assemblies. **(A)** Dotplot of the best sequence alignments of the two haplotype assemblies. Color represents the alignment percent identity between the alternate assembly (Phase1) contig (haplotig) and the primary assembly (Phase0). **(B)** Chromosomal distribution of annotated genes, transposable elements (TE) and large haplotypic structural variants (SVs) between the two phased assemblies. Structural variants were identified by sequence alignment of the two phases. P0 = Phase0 assembly.

### Assessing the quality of the final assembly

When compared to the raw diploid short read data, the final assemblies showed ∼94% completeness and a phred scaled quality score (QV) of >33 (or greater than 99.9995% accurate) (Table 1). More short-read polishing could be performed to increase accuracy; however, this might come at a cost of falsely correcting heterozygosity. While some heterozygous sequence is still missing from the assembly, the majority of 1-copy k-mers are uniquely assigned to one of the haploid assemblies and not shared between them (Figure 2C). These results show that we have accurately produced one full haplotype assembly of TME7 and a second alternate assembly that contains most of the haplotypic variation in this genotype.

We used BUSCO (Simão *et al*., 2015) analysis to verify that we successfully resolved the TME7 haplotypes (Figure 4). The primary (phase0-scaffolded) assembly had a complete BUSCO score of 96.9%, marginally outperforming the AM560-2 v6.1 assembly (complete: 95.1%; duplicated: 5.1%) (Bredeson *et al*., 2016). The majority of complete single BUSCOs (969) are assembled in both phases, yet another 374 are missing from the alternate assembly (Figure 4C). This could be because these BUSCOs are homozygous and thus assembled in the collapsed regions of assembly, and/or due to the missing heterozygosity. Importantly, our deduplicated, TME7 Phase0 assembly only contains 7.9% duplicated BUSCOs, which is comparable to that of AM560-2 and represents a significant improvement compared to ∼15% and ∼19% of the non-haplotype-purged assemblies described in Kuon et al (2019). Interestingly, we identified haplotype-specific complete BUSCOs (Figure 4C), and together the full diploid assembly (Phase0 scaffolds + Phase1 pseudohaplotype contigs) has a complete BUSCO score of greater than that of each phase separately (complete: 98.2%; 80.7% duplicated). This indicates that some BUSCOs may exist in a hemizygous state in the TME7 genome, and that complementation between the phases preserves the existence of these potentially crucial single copy genes.

### Transposable elements and gene annotation

#### Transposable element and repeat annotation

Assembling the repetitive portion of the cassava genome is challenging as it is predicted to contain about 60% repetitive sequence (Figure 1B, Supplementary Figure 1). We used the LTR Assembly Index (LAI) to assess the quality and contiguity of the repetitive sequence assembly (Ou *et al*., 2018). Overall, both haploid assemblies display reference-quality contiguity in the repetitive portions of the genome, with LAI values of 10.53 and 11.17 for the phase0 and phase1 assembly, respectively (Supplementary Figure 6A). Further, we found that the contiguity of the repetitive space in the assembly was much improved compared to the unplaced scaffolds (SupplementaryFigure 6B). We annotated both structurally intact and fragmented transposable elements (TEs) in the full diploid assembly using EDTA (Ou *et al*., 2019). As expected, 59% of the TME7 genome are repeats and transposable elements, which are dominated by LTR retrotransposons that contribute about 50.5% of the genome (Table 2, Supplementary Figure 7). Terminal inverted repeat (TIR) and Helitron DNA transposons contributed 2.43% to the total genome size. There were only marginal differences in TE content between the phases.

**Table 2.** Summary of transposable elements in the TME7 genome assembly.

| Category | Phase0 | Phase1 | Average |
| --- | --- | --- | --- |
| LTR/Copia | 6.24% | 6.25% | 6.25% |
| LTR/Gypsy | 35.36% | 36.22% | 35.79% |
| LTR/unknown | 8.49% | 8.46% | 8.48% |
| TIR/CACTA | 0.64% | 0.63% | 0.64% |
| TIR/Mutator | 0.90% | 0.88% | 0.89% |
| TIR/PIF_Harbinger | 0.13% | 0.16% | 0.15% |
| TIR/Tc1_Mariner | 0.01% | 0.01% | 0.01% |
| TIR/hAT | 0.77% | 0.67% | 0.72% |
| LINE/unknown | 0.44% | 0.44% | 0.44% |
| DNA/Helitron | 0.02% | 0.03% | 0.03% |
| repeat/unknown | 6.05% | 5.45% | 5.75% |
| Total LTR | 50.09% | 50.93% | 50.51% |
| Total DNA TE | 2.47% | 2.38% | 2.43% |
| Total TE | 59.06% | 59.19% | 59.13% |

**Figure 6.**
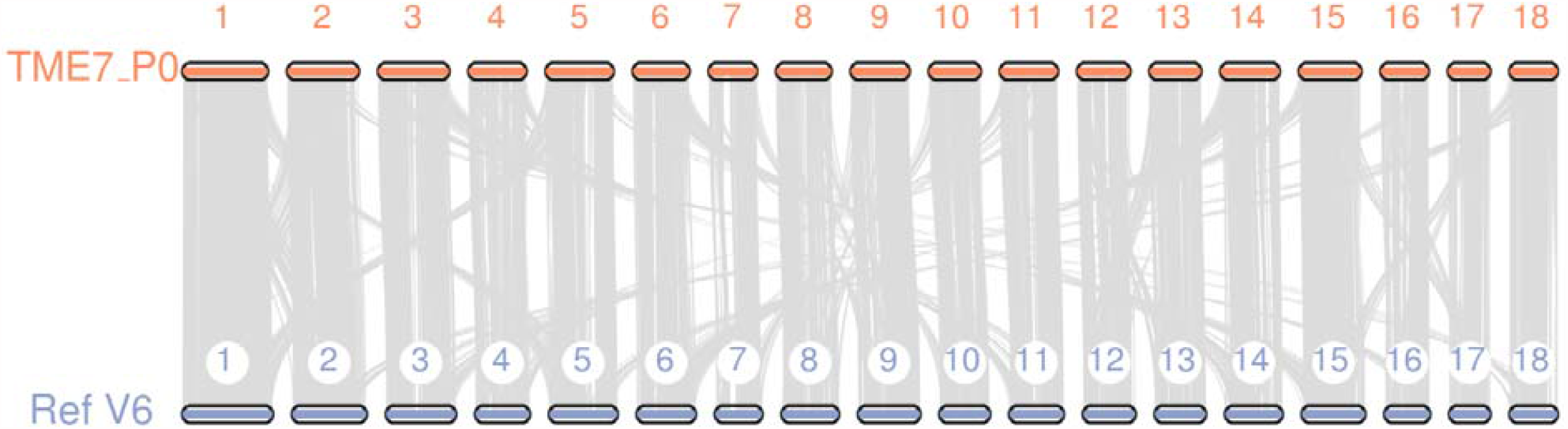
Macro-synteny between of the TME7 and the AM560-2 Ref6.1 genome. Gene synteny comparison between the scaffolded TME7-Phase0 assembly and the AM560-2 reference genome shows largely co-linear genomes with multiple inter-chromosomal duplications attributable to the paleotetraploidy described in cassava in Bredeson et al., (2016).

**Figure 7.**
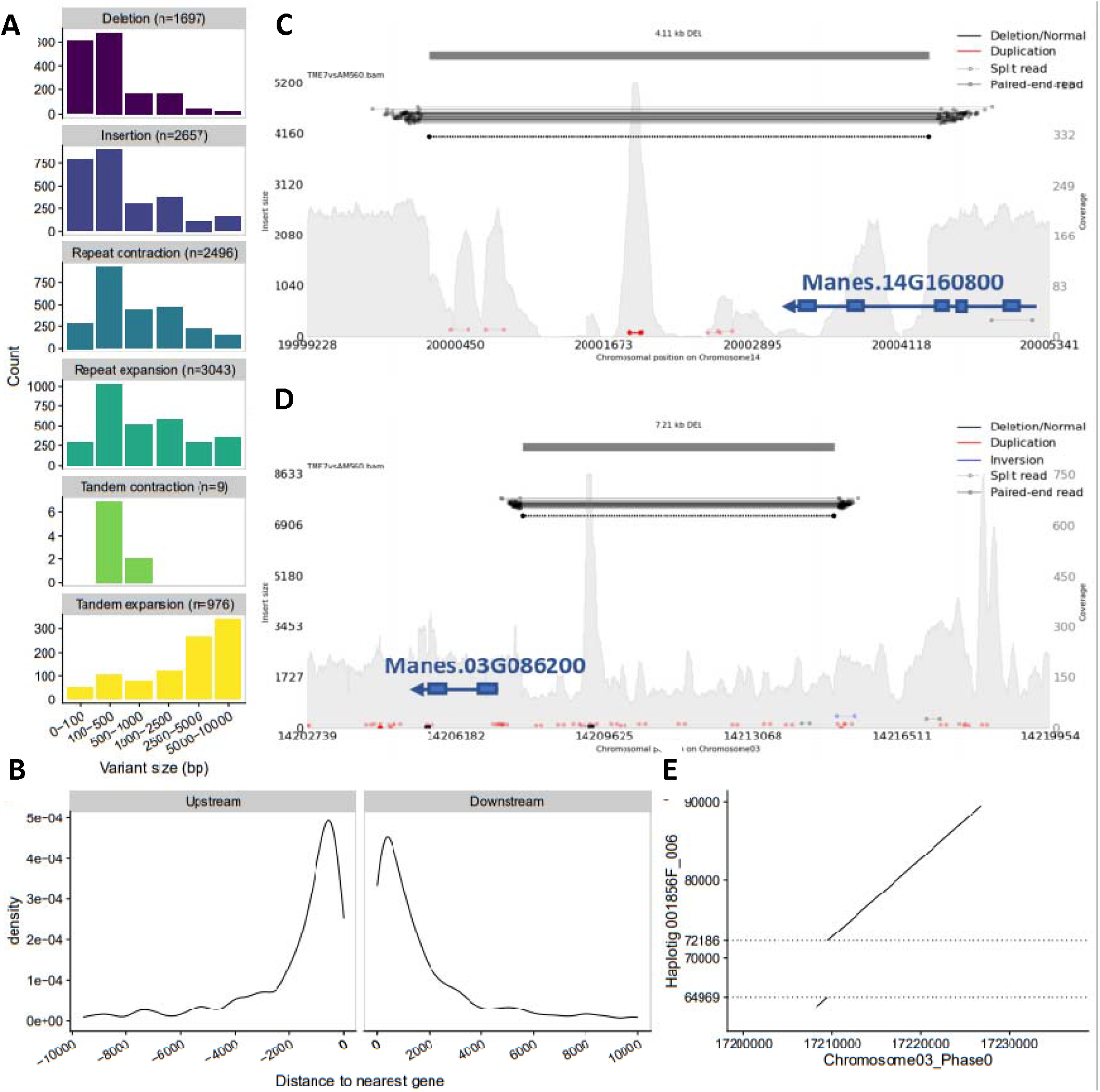
Large structural variants identified in TME7 vs the AM560-2 reference genome. **(A)** The size distribution histograms of structural variants identified by comparison of the phase0 assembly to the AM560-2 reference genome. **(B)** Density of distances (<10 kb away) of large deletions (50-10,000 bp) in TME7 from genes annotated in the AM560-2 reference. **(C and D)** Structural variants interrogated by paired-end reads. Reads with abnormally large insert sizes (color-coded horizontal bars, left y-axis) corroborate deletions identified by alignment of the assemblies. The depth of coverage (grey filled background, right y-axis) aid in determining the zygosity of the deletions. Gene models from the AM560-2 v6.1 annotation are in blue. **(C)** TME7 Phase0 assembly contains a homozygous 4.11 kb deletion compared to chromosome 14 of the AM560-2 Reference genome which overlaps the 3’-end of *Manes*.*14G160800*. **(D)** A 7.21 kb heterozygous deletion is verified on Chromosome 3, potentially overlapping with upstream regulatory region of *Manes*.*03G086200*. Other smaller sequence duplications are also observable (marked in red in **C** and **D**). The 7.21 kb heterozygous deletion in TME7 is correctly phased and assembled as an insertion in haplotig 001856F_006. **(E)** The deletion between 64.9 kb and 72.1 kb on the haplotig, is delineated between the two dashed horizontal lines.

#### Gene annotation and synteny with AM560-2

Gene annotation was performed using the MAKER, AUGUSTUS, and SNAP pipelines including transcript evidence from RNA-seq from 11 tissue types (Wilson *et al*., 2017). We annotated 33,653 and 35,684 genes in phase0 and phase1 assemblies, respectively (Figure 5B). Over 70% of annotated genes had an Annotation Edit Distance (AED) of less than 0.25 suggesting most genes were supported by high evidence levels (Supplementary Figure 8). Comparison of our annotations to that of the AM560-2 ref6 showed that gene synteny between the two cassava genomes was largely conserved, however several macro-level rearrangements are identifiable (Figure 6). Furthermore, this comparison revealed a largely 2:2 pattern of syntenic depth between the annotations (Supplementary Figure 9), consistent with the whole genome duplication described in cassava (Bredeson *et al*., 2016). About 36% of cassava genes exist in one syntenic block reciprocally in either genome, suggesting that these genes may have lost their extra copy since the paleo-duplication. Based on our analysis, it thus appears that the percent of genes which have retained their duplicate status is closer to 60%, rather than ∼36% as previously reported (Bredeson et al. 2016). The prior analysis used homologous genes identified in *Jatropha curcas* as the reference; this likely limited the total numbers of homologs in the analysis, leading to the underestimate of retained duplicated genes. Only 2% of AM560-2 genes were not shared in syntenic blocks in TME7 suggesting they may be unannotated, lost, or translocated out of their block.

**Figure 8.**
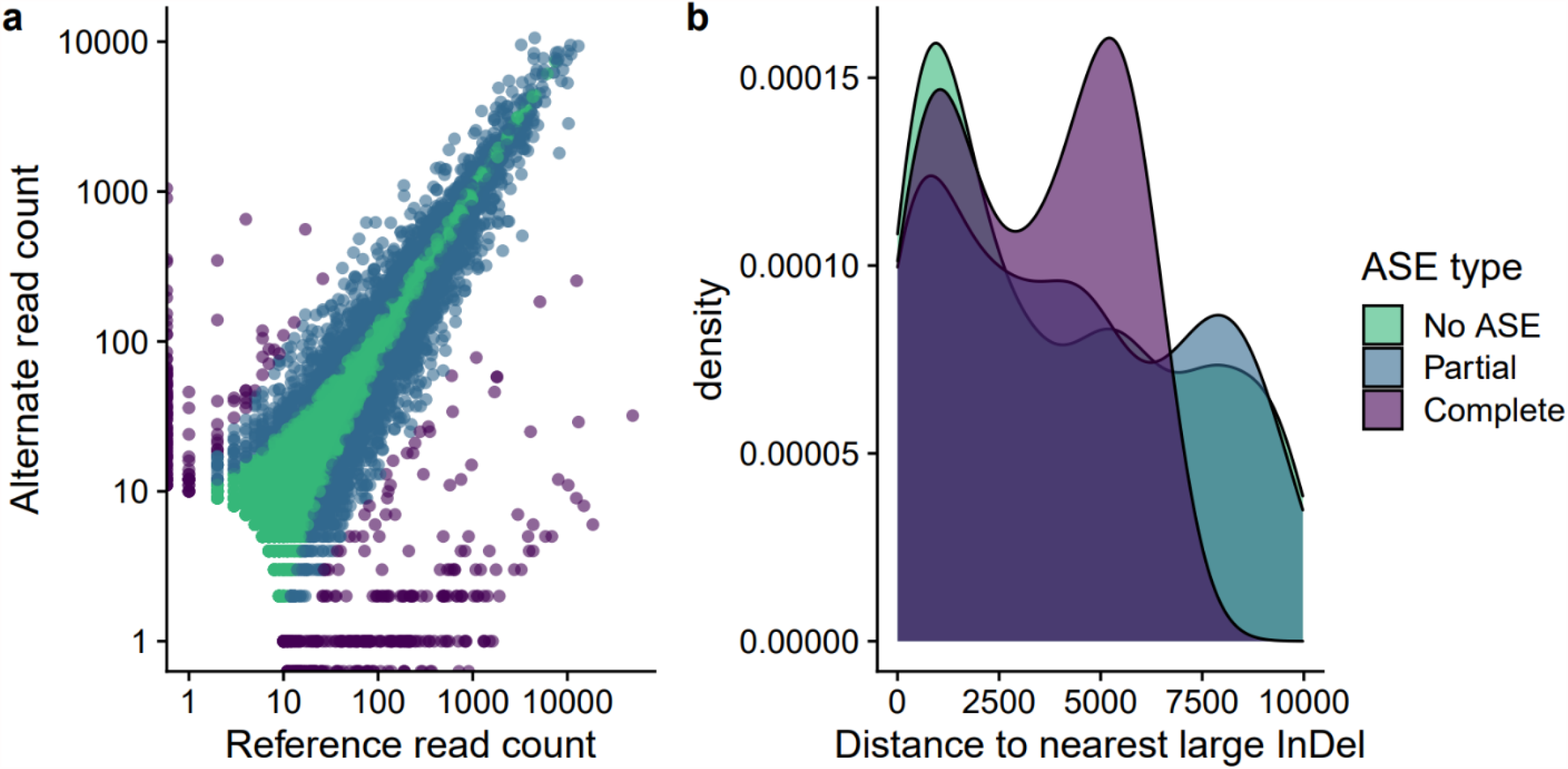
Potential effects of large haplotypic structural variants on allele specific expression. **(A)** Allele specific expression (ASE) patterns in cassava leaf RNA sequencing data. Each point represents an expressed gene and its respective read counts for either the reference or alternate alleles. If greater than 90% of read counts supported one allele of a gene over the other, the gene is characterized as having “Complete ASE” (Purple). Genes showing significant ASE but less than 90% allelic enrichment are categorized as “partial ASE” (Blue). If no significant ASE (FDR > 0.05) was observed genes are denoted in green. **(B)** The distribution of distances to the nearest upstream large insertion or deletion (InDel) for each category of gene.

### Haplotype-specific sequence and structural variation

#### Comparison to the inbred AM560-2 reference

The differences in origin, genome size, and levels of heterozygosity between TME7 and the reference line AM560-2, prompted us to further compare the assemblies. Comparison of the TME7 phase0 assembly to the AM560-2 ref v6.1 assembly revealed 2,257,216 SNPs and 1,666,639 bases affected by small INDELs (<50 bp) that differed (Supplementary Figure 10). We further identified over 10,000 large structural variants (50-10,000 bp) affecting more than 15.99 Mb of sequence (Figure 7A, Supplementary File 1). There is increasing evidence pointing to the importance of large genomic structural variants, and their contribution to phenotypic traits (Alonge *et al*., 2020; Zhou *et al*., 2019). We thus examined the potential effects of the large INDELs (>50 bp) on gene function by measuring the distance to the nearest genes (Figure 7B). Out of 4,354 large INDELs, 1,217 were predicted to be within gene models and another 882 within 2,000 bp upstream of genes, potentially affecting cis-regulatory elements.

To visually validate, and assess the heterozygosity state of several of the largest deletions (>4 kb in length), we aligned short-reads from TME7 to the AM560-2 genome. Both homozygous and heterozygous deletions were identified, and an example of each is in Figure 7C and Figure 7D, respectively. A homozygous deletion identified on Chromosome14, where paired-end reads map to either side of the 4.11 kb deletion and a sharp decline in read coverage is observed, overlaps with the 3’-end of RNA CLEAVAGE STIMULATION FACTOR (Manes.14G160800) (Figure 7C). A heterozygous deletion on Chromosome03, that has read coverage approximately half that of the surrounding area, overlaps the potential promoter region of Manes.03G086200, annotated to encode Ribosomal protein L6 (Figure 7D). This further supports the importance of assembling both haplotypes and suggests that many large haplotypic structural variants might be present with potential impact on gene expression or function.

#### Large haplotypic structural variation in TME7

Recently shown in grape (Zhou *et al*., 2019) and tomato (Alonge *et al*., 2020), large genomic structural variations may have substantive effects on important agricultural traits. For example, the white berries of Chardonnay grape could be a result of a large inversion and deletion, causing hemizygosity at the *MybA* locus (Zhou *et al*., 2019). To further examine the within-genome, haplotypic variation in TME7 we aligned the alternate assembly to the primary assembly. FALCON-Phase has two options for emitting phased assemblies. In “unzip” style, short haplotigs containing alternate sequences are emitted alongside the phased primary contigs (as in FALCON-Unzip). In contrast, in “psuedohap” mode, pseudo-haplotype contigs are generated by collapsing alternate sequence from the phased haplotigs with homozygous sequence from primary assembly. Thus, the pseudo-haplotype alternate assembly might contain artificially homozygous sequences that were missing from the original alternate assembly, originating from lack of assembly or true hemizygosity in the alternate assembly. We therefore used the “unzip”-emit-style haplotigs for comparison to the primary assembly and calculated the mean haplotype divergence to be 2.09% +/-0.18%. We further identified 1,116,832 SNPs and 300,883 small INDELs (<50 bp) in non-repetitive regions, collectively representing more than 2.14 Mb of heterozygous sequence between the two assemblies (Figure 5A). This confirms the high rate of heterozygosity predicted using k-mer based approaches and suggests a well extracted set of haplotigs.

To directly compare the two independently assembled TME7 haplotypes, we aligned the phase1 contigs to the scaffolded phase0 assembly and identified large structural variations (SV). Overall, we identified more than 5,000 variants 50-10,000 bp in size including large insertions, deletions, tandem duplications, and contractions as well as repeat expansions and contractions (Table 3, Figure 5B, Supplementary File 2). The total sequence space that was affected by these structural variants was greater than 8 Mb. Thus, this within-genotype, haplotypic structural variation amounts to greater than half of the between-genotype differences that TME7 has with the AM560-2 reference line. The Assemblytics pipeline can also identify variants greater than 10 kb, however the accuracy with which these are distinguished from translocations or assembly errors is limited (Nattestad and Schatz, 2016). Though we primarily focused on a more conservative approach to identify large SVs, potentially larger haplotypic SVs were identified using Assemblytics. Including SVs up to 50 kb in size in the analysis, yielded close to 16 Mb of sequence affected by SV (Supplementary Figure 11). While these larger SVs should be considered with caution, we note that this is comparable to structural heterozygosity reported in other species such as wine-grape (Minio *et al*., 2019).

**Table 3.** Summary of haplotype-specific structural variants

|  | 50-500 bp<br>Count | 50-500 bp<br>Total bp | 500-10000 bp<br>Count | 500-10000 bp<br>Total bp | Total<br>Count | Total bp |
| --- | --- | --- | --- | --- | --- | --- |
| **Insertions** | 699 | 110,936 | 348 | 791,434 | 1,047 | 902,370 |
| **Deletions** | 676 | 99,453 | 226 | 663,975 | 902 | 763,428 |
| **Repeat expansion** | 649 | 139,116 | 938 | 2,722,146 | 1,587 | 2,861,262 |
| **Repeat contraction** | 668 | 144,659 | 1,136 | 3,486,466 | 1,804 | 3,631,125 |
| **Tandem expansion** | 27 | 5,575 | 31 | 125,712 | 58 | 131,287 |
| **Tandem contraction** | 7 | 819 | 3 | 5,070 | 10 | 5,889 |
|  |  |  |  | **Total:** | 5,408 | 8,295,361 |

### Effects of haplotypic structural variation on allele specific expression

The identified haplotypic SVs are primarily distributed in the chromosome arms and thus are often in close proximity to genes (Figure 5B). For example, the 7,217 bp heterozygous deletion, upstream of *Manes*.*03G086200* (Figure 7C) is correctly phased in our assemblies, as it was detected as an insertion in the phase1 contigs by alignment of the phase1 contigs vs the phase0 scaffolds (Figure 7E). We posited that large haplotype-specific INDELs upstream of genes, such as this one, would impact their allele specific expression (ASE). We thus examined ASE patterns in cassava leaf RNA-seq data (Wilson *et al*., 2017) and observed that of the 14,346 genes expressed in this set, 4,459 showed significant ASE (FDR < 0.05, Supplementary File 3). Such a large number of genes with ASE is congruent with the high heterozygosity of TME7 and may have important biological implications as it has been observed in other heterozygous/hybrid crops (Shao *et al*., 2019; Zhang *et al*., 2020). In hybrid rice for example, patterns of ASE of over 3,000 genes may contribute to the genetic basis of heterosis (Shao *et al*., 2019).

While there could be multiple reasons for ASE of genes (Wood *et al*., 2015; Castel *et al*., 2015), large haplotypic INDELs in cis-regulatory regions, such as the one in Figure 7D, could cause expression of one allele to be severely repressed. We thus defined two categories of ASE genes: If greater than 90% of read counts supported one allele of a gene over the other, we categorized the gene as having “complete ASE.” Conversely, we defined genes as having “partial ASE” if significant ASE was observed, yet allele ratios were not as enriched in either direction. We observed that greater than 12% of genes with ASE show patterns of “complete ASE” (Figure 8A).

We then compared the distribution of distances to the nearest large INDEL between ASE and non-ASE genes. “Complete ASE” genes had significantly different distance distributions from both “partial ASE” and “no ASE” categories (K-S test, *p* < 0.05). Genes with “partial ASE” did not have different distance distributions compared to those with no ASE. For all genes with an INDEL within 10 kb upstream of the transcriptional start site, we further observed that the 26 genes identified in this set with “complete ASE” had different distance distributions, with an enrichment of INDELs around 5,000 bp upstream with a median distance of 3,174 bp to the nearest INDEL, compared to 4,012 and 3,442 bp for “partial-” and no ASE, respectively (Figure 8B). While the genes themselves are not in a hemizygous state, the hemizygosity in their cis-regulatory regions might have important impacts on their allelic expression and potentially on downstream phenotypes. Though this is a narrow dataset of untreated leaf samples, examining the relationship between ASE and SVs in other datasets under additional treatments and/or conditions may further yield important cases where gene expression is affected by large haplotypic SVs (Knowles *et al*., 2017).

Together, the single-nucleotide and large structural variants identified by comparing the two phased TME7 assemblies open a window into the complexity of the heterozygous cassava genome. Work in grapevine and their wild relatives suggests that SVs are primarily deleterious and that they are under strong purifying selection (Zhou *et al*., 2019). Examining the conservation and diversity of large variants within a wide range of farmer-preferred cassava lines would shed light on the effect of SVs on cassava genome evolution in this clonally propagated crop. Further, potentially deleterious alleles such as these large haplotypic SVs, as well as SNPs previously characterized (Ramu *et al*., 2017), warrant further research as these may contribute to limits in inbreeding of cassava.

### Tissue specific gene expression Cassava Atlas

We previously published gene expression patterns for 11 different cassava tissue types based on the AM560-2 reference genome (Wilson et al., 2017). With our newly assembled phased genome, we updated this existing resource. All 11 RNA-seq datasets were mapped to the Phase0 scaffolded and annotated TME7 assembly, and differentially expressed genes were identified as previously described. These results can be further explored at: shiny.danforthcenter.org/cassava_atlas.

### Summary

While recently released assemblies of farmer-preferred cassava lines contain information from both haplotypes in the assembly, the limitation of these assemblies is in the lack of haplotypic purging and sequence deduplication (Kuon *et al*., 2019). Thus, these assemblies do not fully represent either of the haplotypes. Our assembly was successfully deduplicated of most haplotypic sequences, as evidenced by k-mer, BUSCO, and linkage map-based analyses. We further successfully used Hi-C sequencing data to phase and create pseudo-haplotype assemblies. The phased assembly described herein, is thus currently the most accurate assembly of a cassava genotype representative of those grown by millions of subsistence farmers around the world. The differences in genome size compared to the published reference (∼700Mb vs the estimated ∼750Mb for AM560-2), alongside the large SVs identified between the genotypes, showcases how diversity in cassava goes beyond small nucleotide level variation between accessions. We further show that not only does TME7 have large structural variation compared to AM560-2, but that within the genome there are thousands of haplotypic structural variants, potentially perpetuated through clonal variation. Many of these SVs are in close proximity to annotated genes and allelic specific expression of these genes was observed. Further research will help inform how these variants interact and affect gene hemizygosity, copy number, and expression as well as the impact agronomically important traits. We believe this assembly will be an invaluable resource to the cassava research and breeding community, and will further aid in developing tools to ensure food security to those who rely on cassava.

## Data Availability

Both haplotype genome assemblies are stored under NCBI accession number #####. Short and long reads in assembly have been uploaded under the SRA accession #####. Custom scripts used for assembly and analysis are available in Supplementary Files 5 and 6.

## Funding

This work was funded by the Donald Danforth Plant Science Center and the Bill and Melinda Gates Foundation (OPP1093529, OPP1194889 and INV-002958).

## Acknowledgements

We thank Nigel Taylor, Getu Deguma, Narayanan Narayanan, Wilhelm Gruissem, and Yi-Wen Lim for contributing the TME7 tissue samples and Illumina reads and for helpful discussions throughout this research. We thank Alex Harkess for advice on the assembly strategies. We thank Kerrigan Gilbert for discussion and for the critical reading of the manuscript.

## Methods

### Plant material and nucleic acid extraction

Cassava line TME7 (Oko-iyawo) were obtained from Peter Kulakow at IITA in Ibadan, Nigeria. Plantlets were maintained in tissue culture by Nigel Taylor’s lab at the Donald Danforth Plant Science Center. Fresh young leaves were collected for extraction of high molecular weight DNA using a CTAB extraction method (Clarke, 2009).

### Library preparation and sequencing

#### Illumina

Data from Illumina short sequencing DNA libraries of TME7 were provided by Wilhelm Gruissem’s lab at ETH Zurich. After adapter trimming by the sequencing facility, reads were *de novo* de-duped using Nubeam-dedup (Dai and Guan, 2020) prior to further use.

#### PacBio

Initial PacBio sequencing was contributed by Todd Michael in 2016 and did not include size selection prior to sequencing. The PacBio libraries were sequenced on a PacBio RSII system with P6C4 chemistry. A second set of PacBio libraries were constructed using the manufacturer’s protocol and were size selected for 20 kb fragments on the BluePippen system (Sage Science) followed by subsequent purification using AMPure XP beads (Beckman Coulter). Sequencing was performed by the University of Delaware DNA Sequencing & Genotyping Center.

#### Chromatin Conformation Capture sequencing (Hi-C)

Fresh, young cassava leaf material was sent to Dovetail Genomics (Scotts Valley, CA) for DNA extraction, digestion with DpnII, library preparation, and sequencing.

### Genome size and heterozygosity estimation

Flow cytometry protocol was performed at the Benaroya Research Institute at Virginia Mason in Seattle, Washington following their standard methods.

Genome size and heterozygosity were also estimated by means of k-mer counting. We used Jellyfish (Marçais and Kingsford, 2011) to count k-mers of size 21 and plot their depth distributions from the ∼100x Paired-End adapter-trimmed and deduped Illumina sequencing reads of TME7. The maximum k-mer depth was set to 1e6, which allows inclusion of repetitive regions of the genome. We then used the GenomeScope v1 web application (Vurture *et al*., 2017) to model the genome size and heterozygosity for each one of these histograms, and used the model fit to select the best k-mer size for analysis.

### De novo genome assembly and scaffolding

#### Maximizing the diploid assembly

We first assembled the PacBio reads *de novo* using the FALCON and FALCON-Unzip (Chin *et al*., 2016) suite of tools (v1.5.2) which included one round of consensus polishing with quiver. The config files for all FALCON tools are supplied as Supplementary File 4. We further polished only INDELS with 1 round of Pilon (Walker *et al*., 2014). We identified missing heterozygous sequences using Merqury count spectra plots (Rhie *et al*., 2020). The k-mers unique to the short-reads and missing from the assembly were then extracted using Meryl tool set (Rhie *et al*., 2020; Miller *et al*., 2008) and finally extracted the reads containing those k-mers using the function meryl lookup. The short-reads were first down sampled and normalized to ∼100x coverage using BBnorm from the BBTools suite (https://sourceforge.net/projects/bbmap/) then assembled using SPAdes (Bankevich *et al*., 2012) and the resulting contigs were filtered for a minimum coverage depth of 10x and length of 500 bp.

#### Assembly deduplication

The complete set of assembled sequences was concatenated and processed through the purge_dups (Guan *et al*., 2020) pipeline. Alignment coverage histograms inform assembly purging software, such as purge_dups or purge_halpotigs (Roach *et al*., 2018), as to what sequences are potential haplotigs or duplication. While these software packages were developed for use with long reads, we found that short-reads allow for higher resolution when plotting coverage histograms, which in turn results in more accurate sequence purging. Thus we aligned ∼100x deduped PE short-reads to the entire diploid assembly for purging. First, duplicates, caused by retained haplotigs, haplotypic overlaps, and junk contigs, were purged from the primary assembly using manual depth cutoff settings of `5, 76, 126, 151, 252, 453`. A second round of purging on the “haplotig” output of purge_dups was useful to remove duplicates and artifact contigs created by purge_dups during purging of overlaps, again using automatic depth cutoffs (`5, 70, 136, 137, 219, 534`). We then renamed all contigs and haplotigs in the `FALCON-U`nzip naming convention for further processing using scripts in R and python (Supplementary File 5). Briefly, haplotigs which had associated primary contigs in the `dups.bed` file were renamed to match their respective primary contigs. Those that did not have matches (i.e. contigs with low coverage in round 1 of purging etc.) were aligned to the primary assembly using nucmer (Delcher *et al*., 2018) and BLAST. The primary contig with the longest set of alignments was selected as the associated primary contig.

#### Haplotype phasing

The resulting pseudo-haplotype primary contigs and haplotigs alongside the Hi-C data were passed to FALCON-Phase for phase switch correction, creating one complete set of contigs for each phase (Kronenberg *et al*., 2018). However, due to the large number of structural variants between the TME7 haplotypes, we modified the *coords2hp*.*py* script in FALCON-Phase to always include the entire length of the haplotig in placement (Supplementary File 5). This reduced the length of haplotig sequence discarded by FALCON-Phase during phasing. We output the results in both “pseudohap” and “unzip” formats.

#### Scaffolding

The Proximo Hi-C genome scaffolding platform from Phase Genomics’(Seattle, WA) was used to create chromosome-scale scaffolds from the FALCON-Phase phase0 assembly, following the same single-phase scaffolding procedure described in Bickhart *et al*. (2017). As in the LACHESIS method (Burton *et al*., 2013), this process computes a contact frequency matrix from the aligned Hi-C read pairs, normalized by the number of Sau3AI restriction sites (GATC) on each contig, and constructs scaffolds in such a way as to optimize expected contact frequency and other statistical patterns in Hi-C data. Juicebox (Rao *et al*., 2014; Durand *et al*., 2016) was then used to correct scaffolding errors. The Hi-C contact map was created by separately aligning the Hi-C read pairs to the scaffolded genome then generating a Hi-C contact matrix using the command line version of HiCExplorer (Wolff *et al*., 2020). A 10 kb matrix was first created, then bins were merged to get a 500 kb resolution for ease of plotting. Bin interaction data was then exported to table separated format (tsv) then imported to R for plotting.

### Assembly quality assessment

#### Linkage map alignment

To further confirm the order and contiguity of the assembly we aligned the 22k marker composite linkage map (ICGMC, 2015) from cassava base (cassavabase.org). In this map, each SNP marker is aligned to the cassava v4.1 draft genome assembly and a scaffold and physical position is reported alongside the genetic position. Using the *marker_seqs*.*py* python script (Supplementary File 5) we extracted sequence from 100 nt on both sides of each SNP in the v4.1 assembly. If the SNP marker was closer than 100 nt from the end of a scaffold, then the sequence with the maximum length possible around that SNP was extracted. These ∼200 nt sequence tags were then aligned via BLAST to each phase of the current assembly. The numbers of uniquely mapping markers with alignment length >150 nt and >95% identity were used to assess levels of sequence duplication.

#### K-mer based evaluation

Merqury (Rhie *et al*., 2020) and the built-in Meryl implementation were used to enumerate the k-mer distribution in the Illumina PE reads and compare it to the diploid and haploid assemblies. Using the provided script in Merqury, a k-mer of 21 was selected to best represent a genome size of ∼700 Mb. Copy number spectra and assembly spectra were plotted using the hist files provided and ggplot2. When k-mer distributions were used to estimate genome sequence length (i.e. to measure missing sequence space), the sum of counts of k-mers under the respective distribution was divided by the mean k-mer multiplicity of the distribution: (*sum*(*kmer count* * *kmer multiplicity*))/*mean*(*kmer multiplicity*)

### Haplotype-specific annotation

#### Transposable element annotation and repeat masking

Transposable elements (TEs) of each assembly were independently annotated using EDTA v1.9.7 (Ou *et al*., 2019) with parameters `‘--sensitive 1 --anno 1 -t 18`’ and ‘`--cds`’ providing the coding sequences of the *M. esculenta* v6.1 assembly. Library sequences from the *de novo* TE library generated by EDTA were filtered and those present more than three full-length copies in the respective haploid assembly were retained. The remaining sequences from the two TE libraries were combined using the ‘*make_panTElib*.*pl*’ script in the EDTA package, generating a high-quality TE library. The final TE library was then used to annotate the two haploid genomes using RepeatMasker v4.1.1 (www.repeatmasker.org) with parameters ‘`-q -no_is -norna -nolow -div 40 -cutoff 225`’ that allow for up to 40% of sequence divergence. This step helped to annotate fragmented TEs. To consistently annotate intact TEs in the two haploid genomes, the final TE library and the final homology-based TE annotation were provided to EDTA with parameters `‘--evaluate 1 --anno 1 -t 18 --step final`’. In depth commands for TE annotation and LAI calculation are supplied in Supplementary File 5.

#### Gene annotation

Transcriptome data of 11 tissue types (Wilson *et al*., 2017) was used to generate transcript evidence for annotation. Reads were trimmed with Trimmomatic (Bolger *et al*., 2014) and aligned to the soft masked diploid reference (Phase0 scaffolds + Phase1 pseudohaplotype contigs concatenated) using Hisat2 v2.1.0 (Kim *et al*., 2019). Stringtie v1.3.5 was used to assemble transcripts from each alignment file and all files were merged with ‘stringtie merge’ (Pertea *et al*., 2015). A fasta containing CDS for all transcripts was produced using gffread tool from the cufflinks (Trapnell *et al*., 2010) package. These transcripts, together with AM560-2 v6.1 CDS sequences and protein sequence from Araport11 (Cheng *et al*., 2017), were used for a first round of MAKER v2.31.8 (Cantarel *et al*., 2008) gene annotation. Gene prediction was further performed by training SNAP (library 2013-02-16) (Korf, 2004) and AUGUSTUS v3.3 (Stanke and Morgenstern, 2005) as suggested in (Bowman *et al*., 2017) and the output of the first round of MAKER annotation. After gene prediction the genes in the gff file were renamed and the file was split to produce one gff for each phase.

#### Gene synteny analysis

Comparison of gene synteny between the TME7 phase0 assembly and the AM560-2 ref6 assembly was performed with the Python MCScanX pipeline v1.1.12 (Tang *et al*., 2008; Wang *et al*., 2012). Briefly, annotation gff files were converted to bed format keeping one isoform per gene using `jcvi.formats.gff --primary_only.` A pairwise synteny search was performed and the high quality synteny block (anchors) were used in syntenic depth comparisons and plotting of karyotypes and dot plots.

#### Assessment of genic and repetitive sequence space

The completeness and duplication of the genic regions in the assembly was performed by using BUSCO v4.1.2 (Simão *et al*., 2015) benchmark software (http://busco.ezlab.org/) and the “eudicotyledons_odb10” ortholog dataset with default settings.

To evaluate the contiguity of the repetitive sequence assembly, the LTR Assembly Index (LAI) was evaluated using LAI beta3.2 (Ou *et al*., 2018) with input files generated by EDTA. The initial LAI estimation was done using the ‘`-q`’ parameter, then average LTR identity and total LTR content were obtained and further provided to the standardization of LAI, with parameters ‘`-iden 95.63 - totLTR 53`’. Regional LAI was calculated in 3 Mb windows with 300 kb overlapping steps.

### Structural variation and polymorphisms

Structural variants between TME7 and the AM560-2 reference genome were identified by aligning the phase0 contigs vs the reference genome. The authors of the Assemblytics (Nattestad and Schatz, 2016) software recommend analysis using contigs and not scaffolds, to minimize bias from different gap sizes in the assembly. Thus, initially the reference assembly was split at gaps of greater than ten Ns using the python script *split_scaffolds*.*py* (Supplementary File 5). After alignment with nucmer (Delcher *et al*., 2018) with settings: `--maxmatch -l 100 -c 500` the delta file was gziped and uploaded to the Assemblytics web interface (www.assemblytics.com) for analysis. The results were exported as a bed file and imported into R for plotting. Dot plots of the alignments were produced using scripts modified from https://jmonlong.github.io/Hippocamplus/2017/09/19/mummerplots-with-ggplot2/ (Supplementary File 6).

The locations of the five largest deletions identified were then examined for evidence of structural variation using short read mapping. Deduplicated Illumina reads from TME7 were aligned to the AM560- 2 v6.1 reference using bwa mem (Li and Durbin, 2009). The sorted bam file was then loaded into samplot (Belyeu *et al*., 2021) to plot the read coverage and identification of discordant mapping. SNPs and INDELs were identified by using *dnadiff* and *show-snps* programs in the MUMmer4 package (Delcher *et al*., 2018).

Structural variation between the phases was then assessed by aligning phase1 unzip contigs vs. phase0 scaffolds (split at >10 Ns) and using Assemblytics as above. Haplotype divergence was calculated by aligning the FALCON-Phase “Unzip”-emit-style haplotigs to the primary, Phase0, assembly using nucmer with these settings: `--maxmatch -l 100 -c 500`. Alignments were filtered with delta-filter -g and coordinates were output using show-coords. Finally, divergence from the primary assembly was calculated using scripts from https://github.com/skingan/FC_Unzip_HaplotypeDivergence. SNPs and INDELs between the phases were identified as above. Distances of genes to structural variants were measured using bedtools *closest* command (Quinlan and Hall, 2010).

### Allele specific expression

We aligned leaf RNA-seq data from Wilson et al. (2017), to the TME7 phase0 assembly using STAR v2.7.8 (Dobin *et al*., 2013). Alignments were then deduplicated with Picard tools and SNPs were called using GATK v4.1.4.1 (Van der Auwera *et al*., 2013). After minimal quality filtering (`QD < 2.0, FS` `> 60.0, MQ < 40.0, MQRankSum < -12.5, ReadPosRankSum < -8.0`), the SNP VCF file was then imported to phASER (Castel *et al*., 2016) to accurately phase the variants within each gene model. PhASER settings were `--paired_end 1 --mapq 255 --baseq 10`. Haplotypic read counts per gene were then exported using the ‘phASER Gene AE’ tool and read into R for statistical analysis. For each gene, the REF and ALT read counts were compared using a binomial test and p-values were Bonferroni corrected. Genes with a false discovery rate of less than 0.05 were considered as showing ASE. We further categorized ASE genes as having “Complete ASE” or “Partial ASE” if allele ratios were greater or less than 0.9 towards one allele respectively. Distances to nearest INDEL were measured using bedtools *closest* command (Quinlan and Hall, 2010) and the distributions of distances of genes in different ASE categories were compared using the Kolmogorov–Smirnov test.

### SHINY app update

Reads from the RNA-seq dataset for 11 tissue types were aligned to the TME7 Phase0 assembly using HISAT2 (Kim *et al*., 2019) and abundance was quantified with Stringtie (Pertea *et al*., 2015). Read counts were transformed into robust-FPKMs using DESeq2 (Love *et al*., 2014). Finally, the annotation was matched to the transcript IDs and formatted to be read within the Shiny framework.

### Scripts and figures

All scripts described above are supplied in Supplementary File 5. All R scripts for producing figures and summary results are supplied in Supplementary File 6.

## Notes

### Competing Interest Statement

The authors have declared no competing interest.

