## Supplemental Figures for "Large structural variations in the haplotype-resolved African cassava genome"

*Author order TBD:*

^1^Ben N. Mansfeld: 0000-0001-6118-6409

^1^Adam Boyher: 0000-0001-9681-1817

^1^Jeffrey C. Berry: 0000-0002-8064-9787

^1^Mark Wilson:

^2^Shujun Ou: 0000-0001-5938-7180

^1^Seth Polydore: 0000-0002-7779-7367

^3^Todd P. Michael: 0000-0001-6272-2875

^1^Noah Fahlgren: 0000-0002-5597-4537

^1^Rebecca S. Bart: 0000-0003-1378-3481

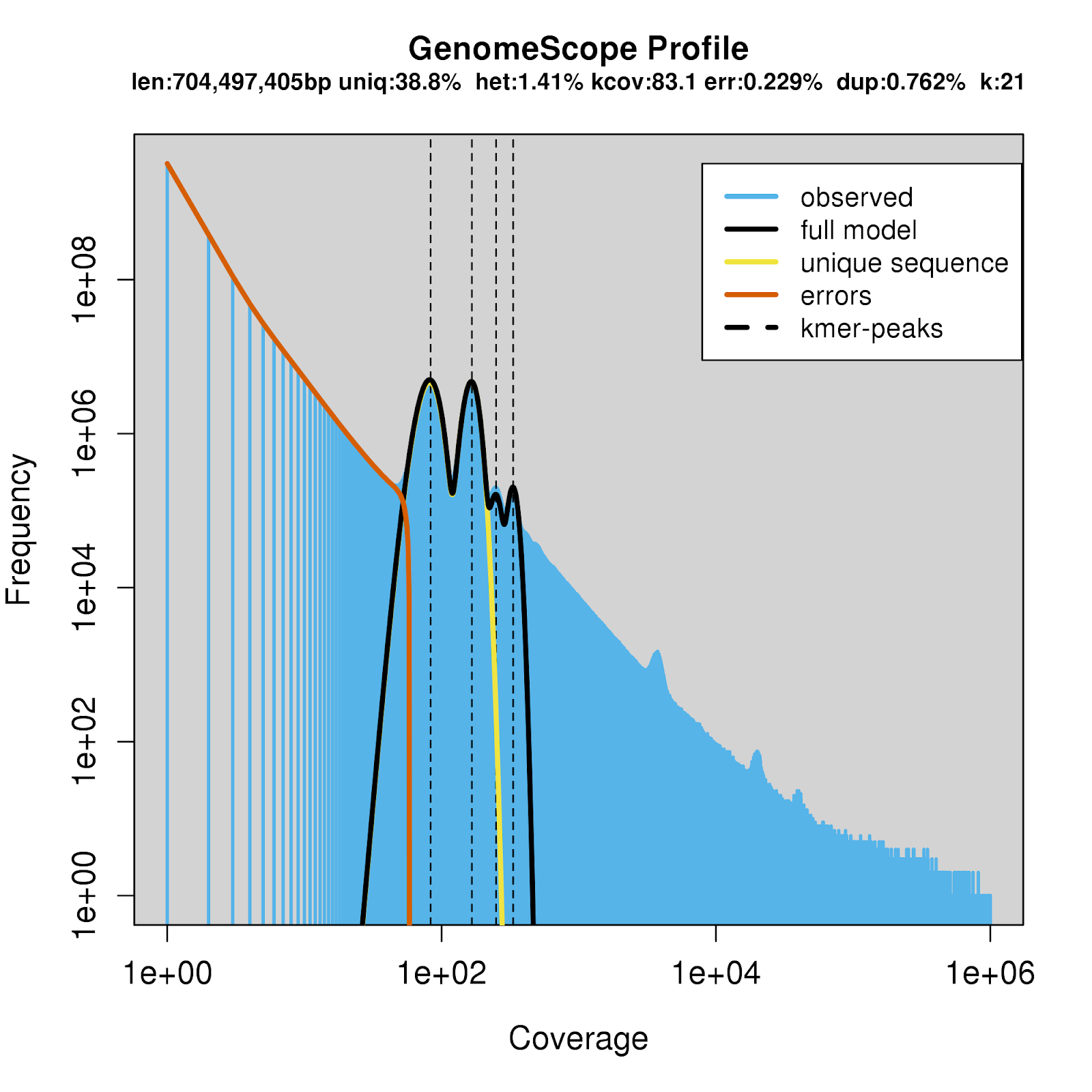
**Supplementary Figure 1. Estimation of genome size, heterozygosity, and repetitiveness using GenomeScope Profile.** The k-mer spectra from Illumina reads were used to estimate the genome parameters. The full k-mer spectra are shown here compared to main text Figure 1. K-mer size was set to 21, and k-mer coverage cutoff was set at 1e6 to include repeat regions in genome size estimates. The haploid genome size was estimated to be 704 Mb consisting of 61% repetitive sequence and a heterozygosity of 1.41%.

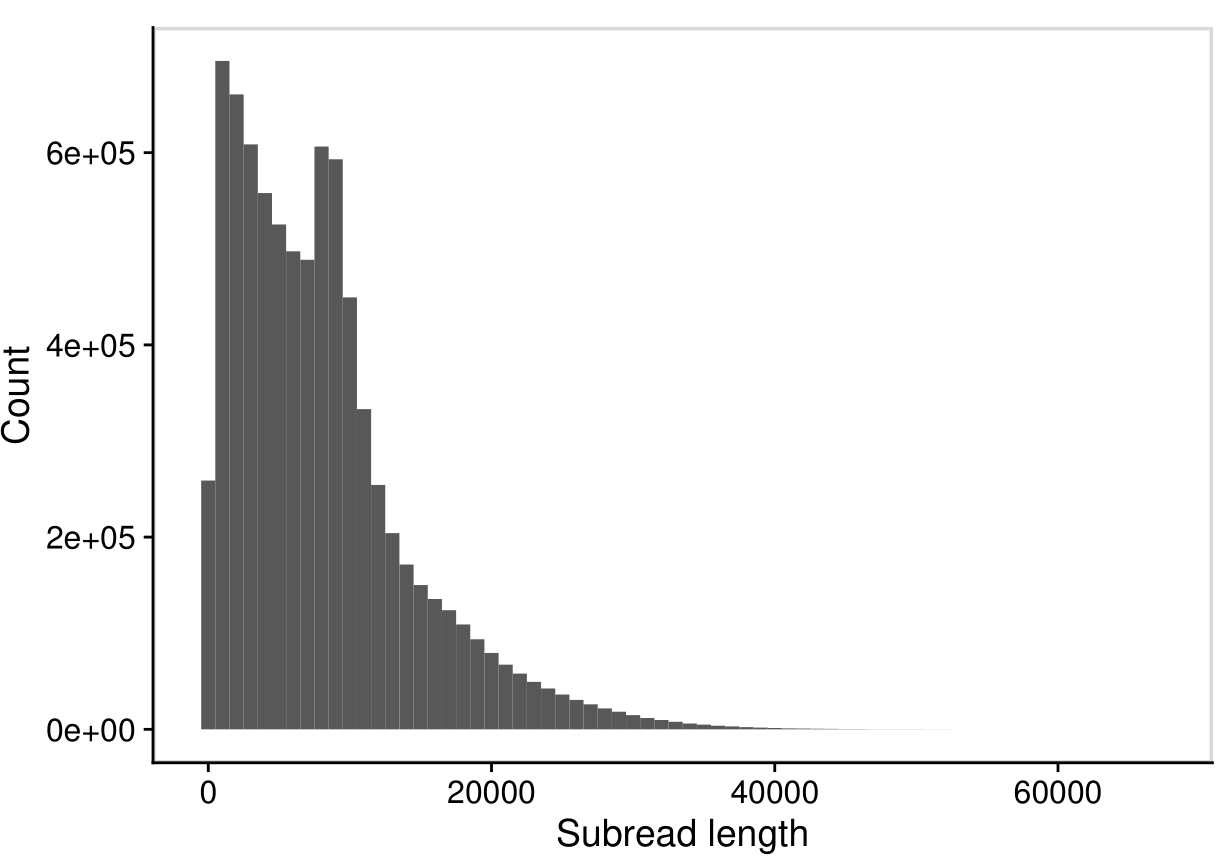

**Supplementary Figure 2. Pacific Bioscience sequencing subread length distribution.** Sequencing was done at two different times. The first round did not include a blue pippin size selection while the second round did; this can be observed as two peaks in the read length distribution.

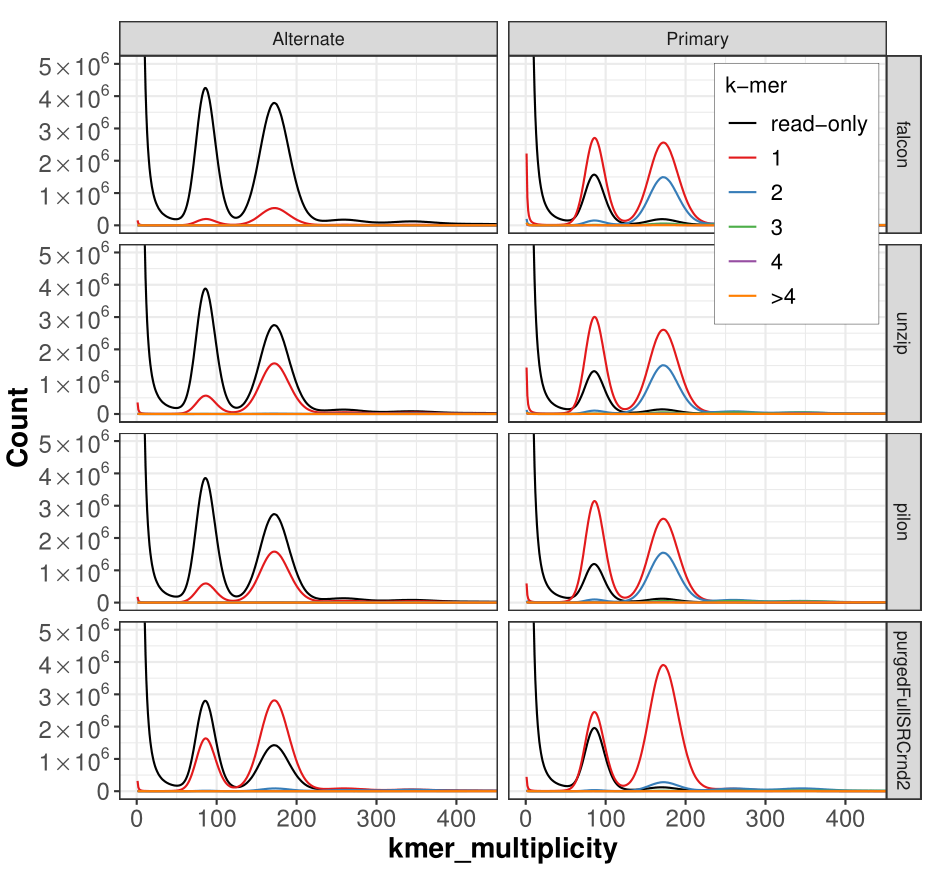
**Supplementary Figure 3. Diploid k-mer count spectra for different assembly stages.** Short read k-mer distribution plots are colored by the number of times a k-mer is present in every diploid (primary + alternate) assembly. K-mers denoted in black are missing from the assembly and represent probable short read sequencing errors (k-mer multiplicity <~ 50) or missing assembled sequence (≥~50). Purging the primary assembly (bottom panel) using purge_dups, greatly reduced the duplicated homozygous k-mer levels (blue peak ~175 k-mer multiplicity) in the primary assembly and increased the heterozygous k-mers in the alternate assembly (red peak ~75 k-mer multiplicity). This indicates that the haplotypes are well resolved.

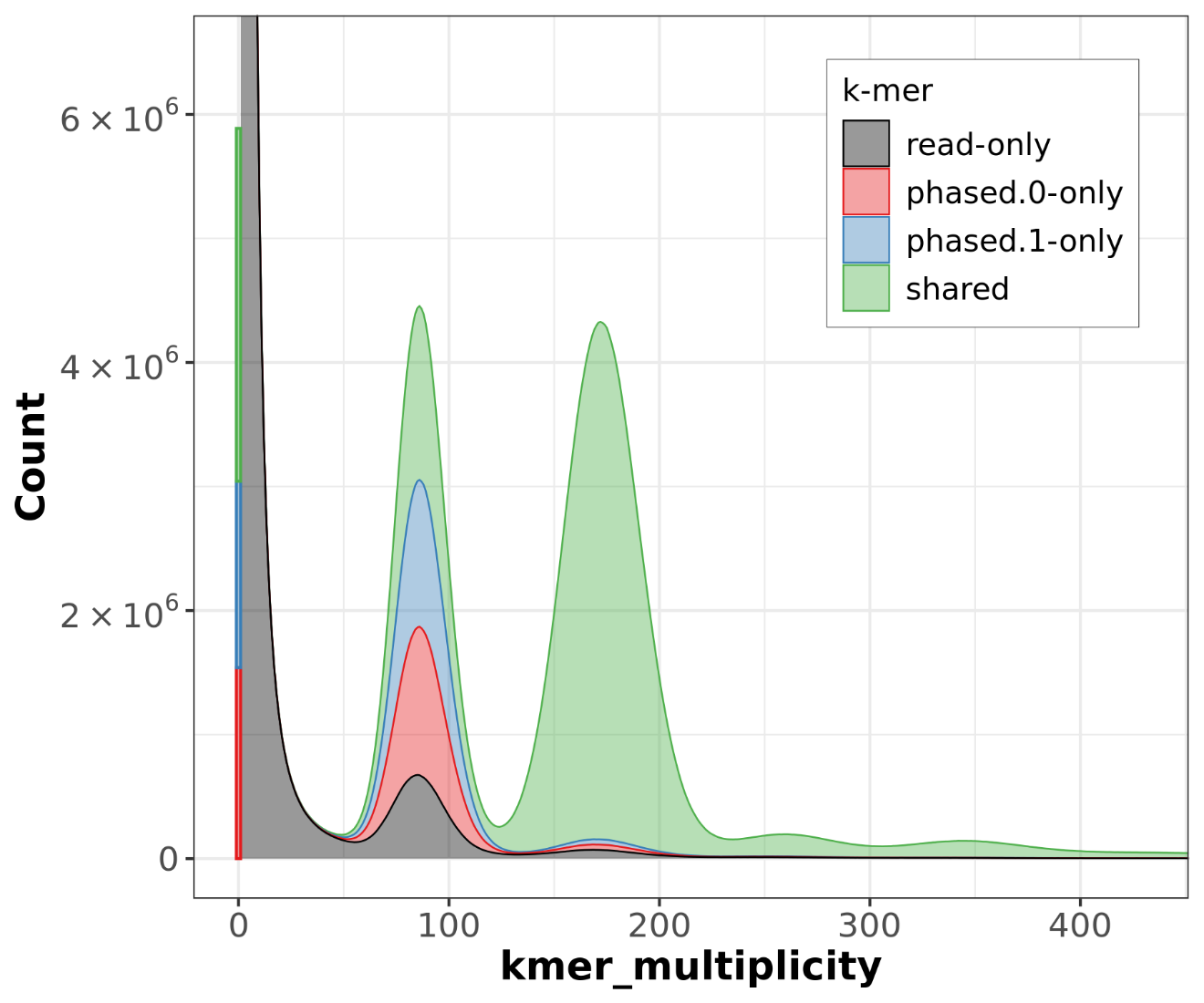
**Supplementary Figure 4. FALCON-Phase causes a reduction in heterozygous sequence with default settings.** With comparison to main text Figure 2C, a stark reduction in phase1-unique sequence (blue) is observed when using the default setting. Furthermore, an increase in “read-only” k-mers which are missing from the assembly is also observed.

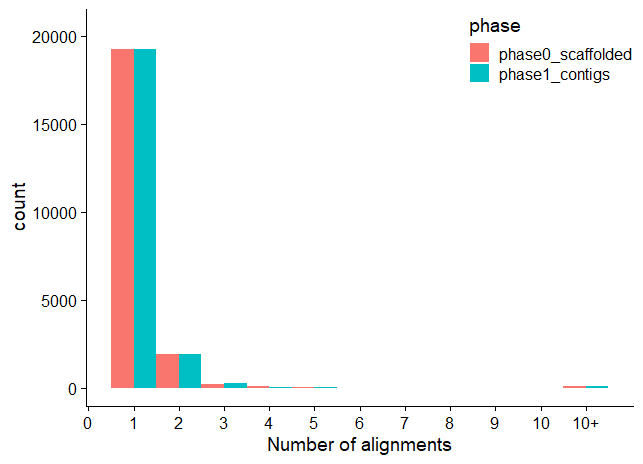

**A**

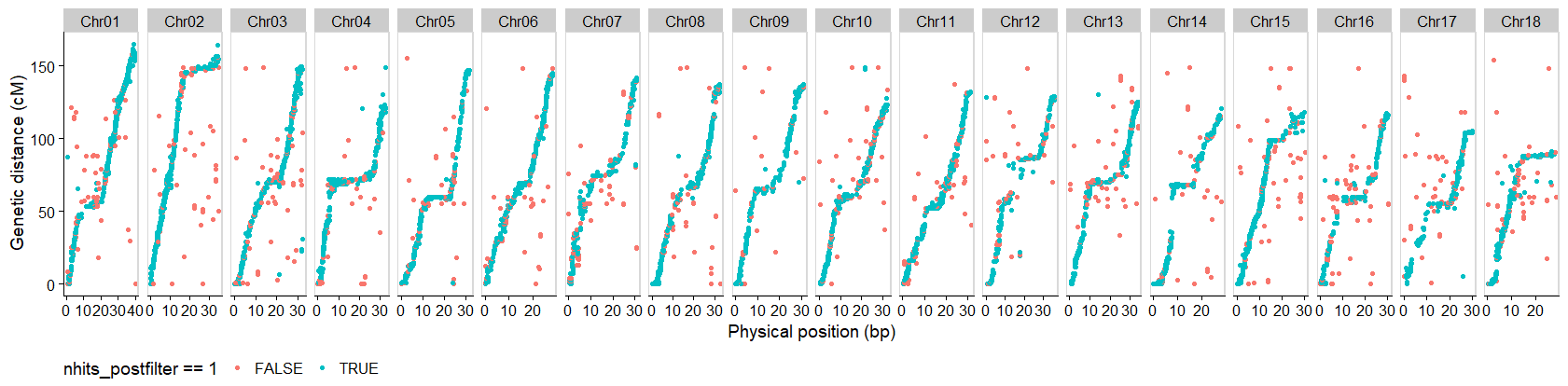

**B**

**Supplementary Figure 5. Linkage map duplication rates are minimal in the TME7 assembly. (A)** The number of filtered alignments for each of the 22K markers from the Cassava Linkage Map project. **(B)** Distribution of duplicated markers in the Phase0 assembly. Markers in blue exist exactly once in the assembly.

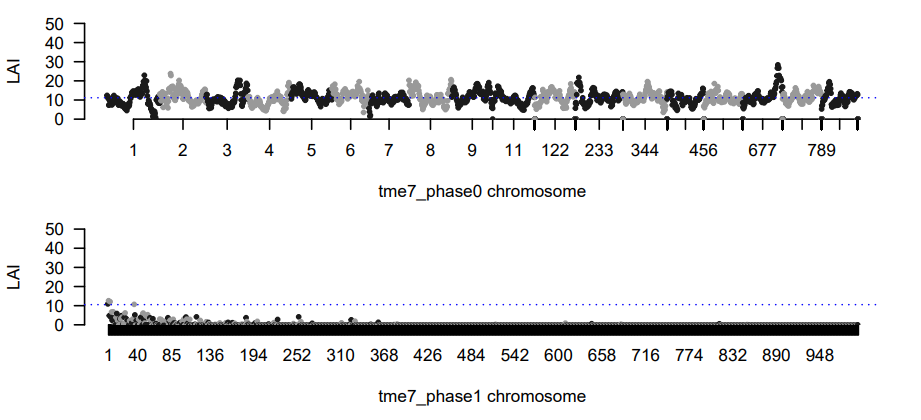

**B**

**A**

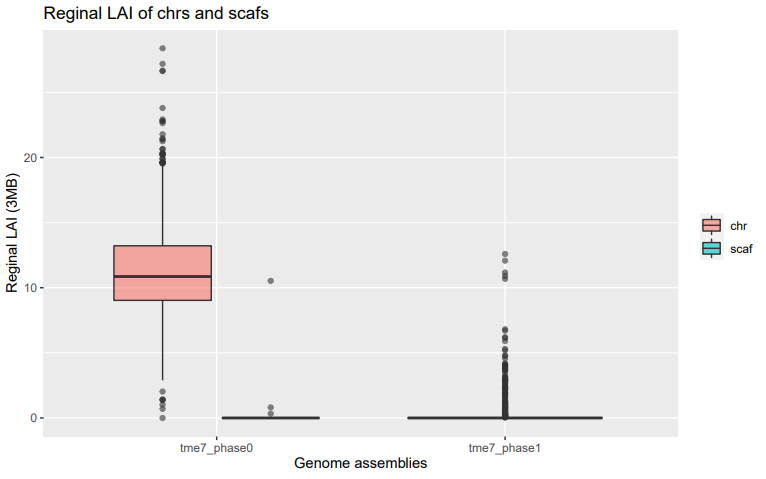

**Supplementary Figure 6. Distributions of Long Terminal Repeats (LTR) assembly index. (A)** LAI (LTR Assembly Index) as a measure of the completeness of the repetitive sequence in the two haplotype assemblies. **(B)** As expected, scaffolding increases the regional LAI.

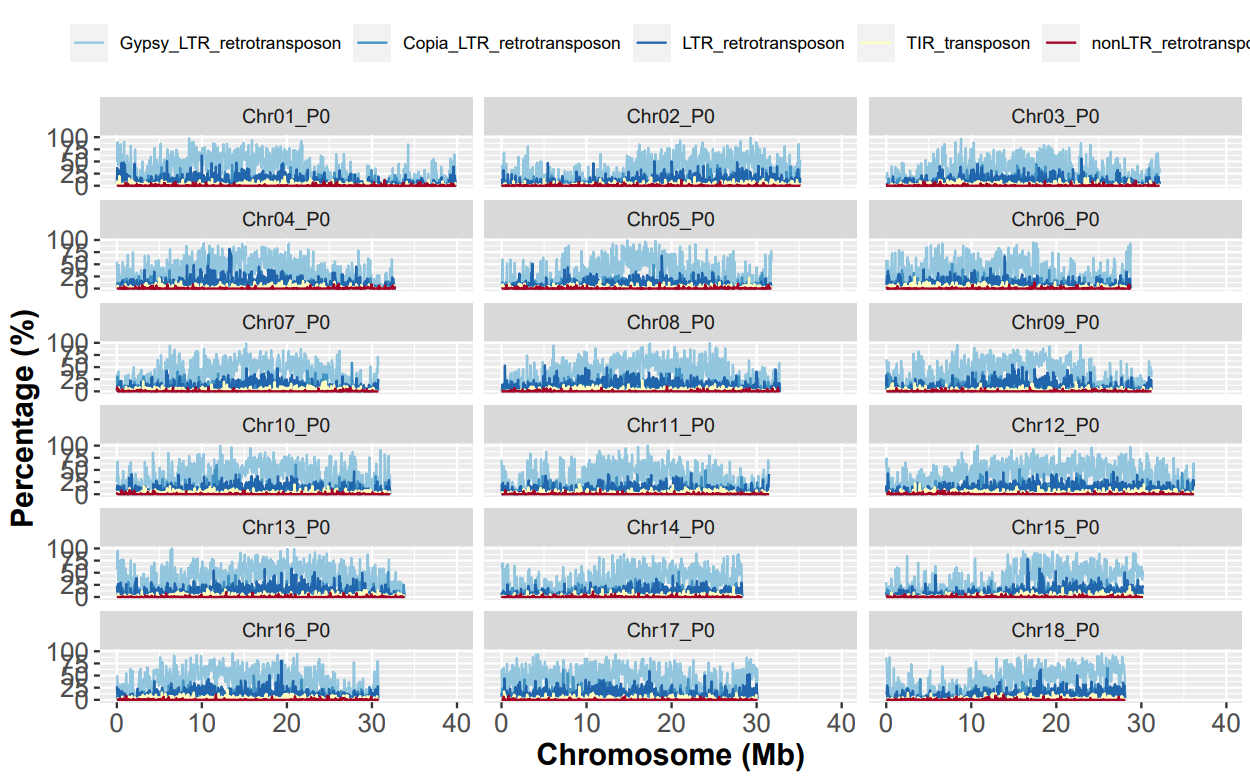
**Supplementary Figure 7. Distributions and types of transposable elements in the TME7 Phase0 assembly.** Transposable element annotation of the TME7 genome showed that ~59% of the genome is comprised of repeats and transposable elements. The repeat landscape is dominated by LTR retrotransposons that contribute about 50.5% of the genome. Terminal inverted repeat (TIR) and Helitron DNA transposons were about 2.43% of the total genome size.

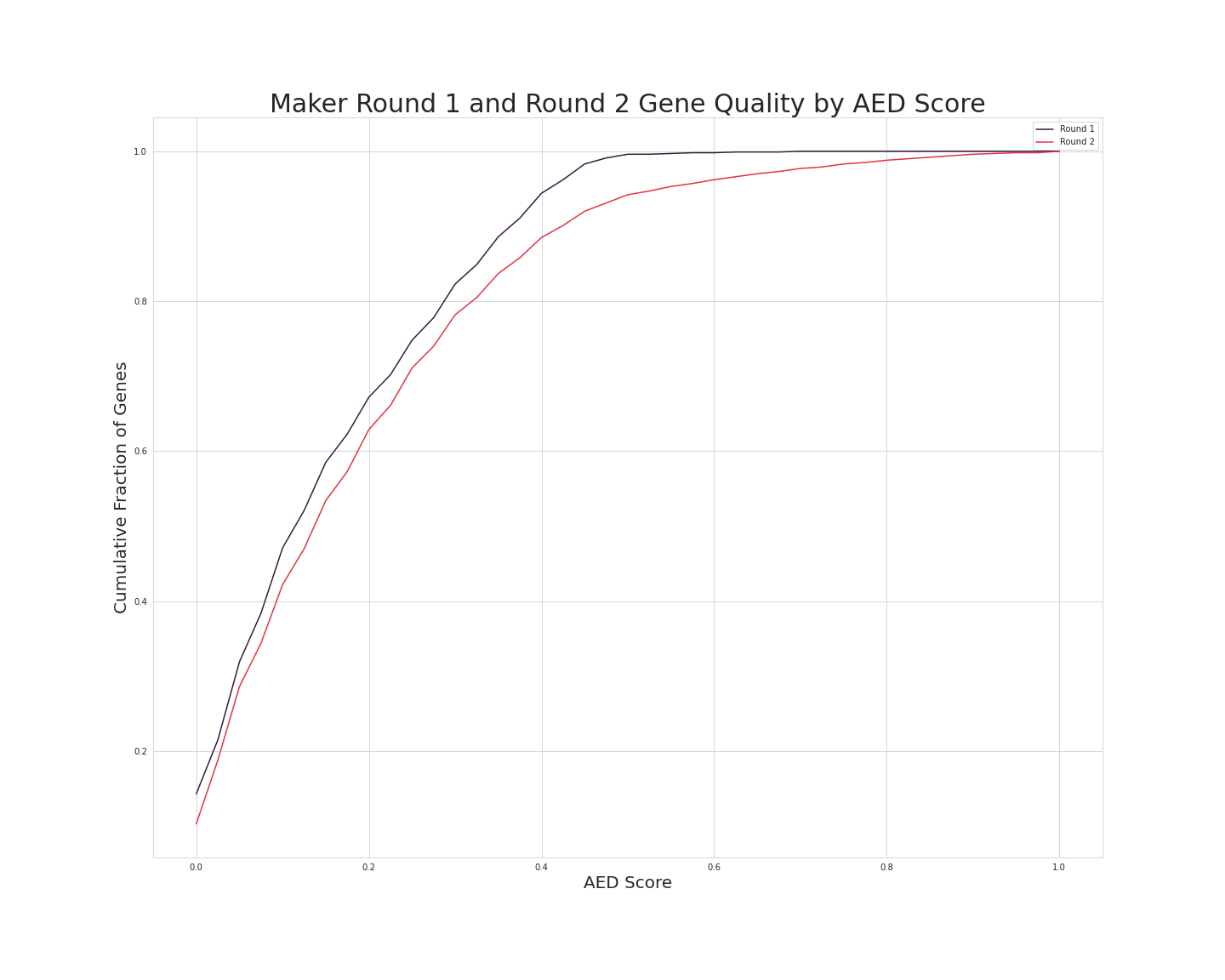
**Supplementary Figure 8. The cumulative Annotation Edit Distance (AED) distribution of two rounds of annotation.** Round 1 (black) represents an evidence-based gene annotation step with MAKER, while Round 2 (red) includes gene prediction using AUGUSTUS and SNAP. Genes with lower AED have high evidential support.

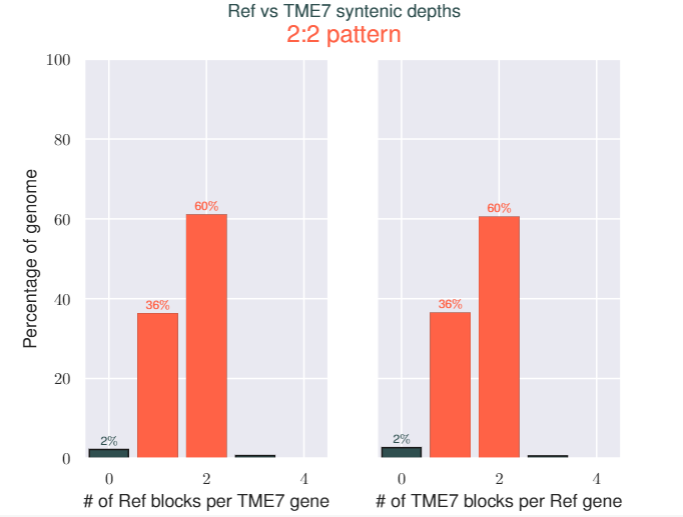
 **Supplementary Figure 9. Percent of genes and their syntenic depths in blocks between TME7 (Phase0) and AM560-2 reference genome v6.1.** Syntenic blocks of genes were compared using MCScanX and the numbers of genes existing in single, and multiple blocks is reported. Most of the genes exist in exactly two syntenic blocks in the reciprocal comparisons due to the paleo tetraploidization of the cassava genome.

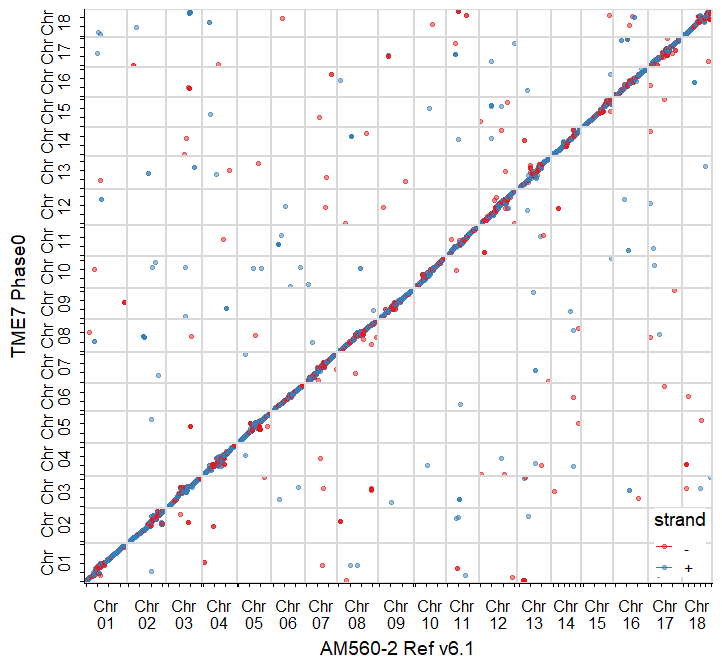

**Supplementary Figure 10. Alignment of TME7 Phase0 assembly to the homozygous AM560-2 reference.** Dotplot of the best sequence alignments of the assemblies. Color represents the alignment strand on the reference assembly.

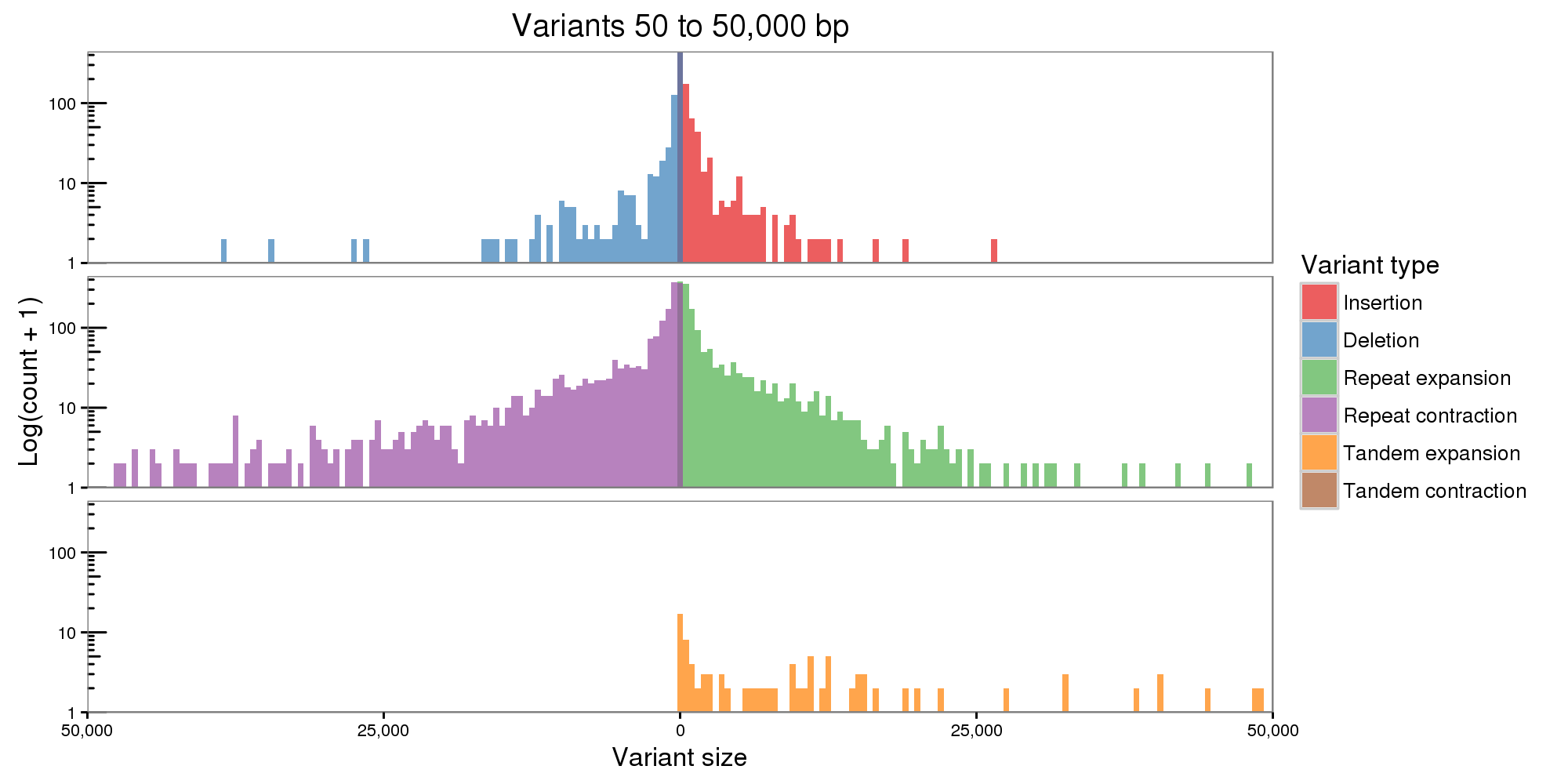

**Supplementary Figure 11. Distribution of haplotypic structural variant sizes when comparing the Phase1 to Phase0 assemblies.** The two assemblies were split at N’s and the contigs were aligned to identify structural variants (SV). Expanding the analysis from the default maximum of 10 kb to include variants up to 50 kb identified a further ~8 Mb of sequences affected by SV. Inclusion of longer SV sizes has a lower specificity.
