## Supplementary File 5 for "Large structural variations in the haplotype-resolved African cassava genome"

# This file contains additional scripts used in the assembly and analysis

### Transposable element annotation and LAI pipeline 

Sequence names of each assembly were trimmed for simplicity.

perl -i -nle 's/omosome//; s/ffold//; s/hase//; print $_' genome.fasta

Transposable elements (TEs) of each assembly were independently annotated using EDTA v1.9.7 (Ou et al. 2019) with parameters ‘--sensitive 1 --anno 1 -t 18’ and ‘--cds’ providing the coding sequences of the M. esculenta v6.1 assembly. 

# run EDTA
nohup perl ~/las/git_bin/EDTA/EDTA.pl --genome cassava_tme7_phase0_scaffolded_renamed.fasta --cds Mesculenta_305_v6.1.cds.fa --sensitive 1 --anno 1 -t 18 &
nohup perl ~/las/git_bin/EDTA/EDTA.pl --genome cassava_tme7_phase1_pseudohap_contigs.fasta --cds Mesculenta_305_v6.1.cds.fa --sensitive 1 --anno 1 -t 18 &

Then library sequences from the de novo TE library generated by EDTA were filtered and those present more than three full-length copies in the respective haploid assembly were retained. 

# get fl3cp.pass lib seq for all genomes
 for i in cassava_tme7_phase0_scaffolded_renamed.fasta cassava_tme7_phase1_pseudohap_contigs.fasta; do perl ~/las/git_bin/EDTA/util/find_flTE.pl $i.mod.EDTA.anno/$i.mod.out|awk '{print $10}'|sort|uniq -c > $i.out.flcp & done
 for i in cassava_tme7_phase0_scaffolded_renamed.fasta cassava_tme7_phase1_pseudohap_contigs.fasta; do for j in `awk '{if ($1>3) print $2}' $i.out.flcp`; do grep $j $i.mod.EDTA.TElib.fa; done > $i.mod.EDTA.TElib.fa.fl3cp.list & done
 for i in cassava_tme7_phase0_scaffolded_renamed.fasta cassava_tme7_phase1_pseudohap_contigs.fasta; do perl ~/las/git_bin/EDTA/util/output_by_list.pl 1 $i.mod.EDTA.TElib.fa 1 $i.mod.EDTA.TElib.fa.fl3cp.list -FA > $i.mod.EDTA.TElib.fa.fl3cp.fa & done

The remaining sequences from the two TE libraries were combined using the ‘make_panTElib.pl’ script in the EDTA package, generating a high-quality TE library.

# make pangenome TE lib
nohup perl ~/las/git_bin/EDTA/util/make_panTElib.pl -liblist lib.list -t 36 &
 
The final TE library was then used to annotate the two haploid genomes using RepeatMasker v4.1.1 (Smit and Hubley 2015) with parameters ‘-q -no_is -norna -nolow -div 40 -cutoff 225’ that allow for up to 40% of sequence divergence. This step helped to annotate fragmented TEs consistently. 

# annotate with pangenome lib
RepeatMasker -pa 36 -q -no_is -norna -nolow -div 40 -lib cassava_tme7_EDTA.TElib.fa -cutoff 225 cassava_tme7_phase0_scaffolded_renamed.fasta.mod &
RepeatMasker -pa 36 -q -no_is -norna -nolow -div 40 -lib cassava_tme7_EDTA.TElib.fa -cutoff 225 cassava_tme7_phase0_scaffolded_renamed.fasta.mod &

To consistently annotate intact TEs in the two haploid genomes, the final TE library and the final homology-based TE annotation were provided to EDTA with parameters ‘--evaluate 1 --anno 1 -t 18 --step final’.

# reannotate 
nohup perl ~/las/git_bin/EDTA/EDTA.pl --genome cassava_tme7_phase0_scaffolded_renamed.fasta --cds Mesculenta_305_v6.1.cds.fa --rmout cassava_tme7_phase0_scaffolded_renamed.fasta.mod.out --curatedlib cassava_tme7_EDTA.TElib.fa --evaluate 1 --anno 1 -t 18 --step final &
nohup perl ~/las/git_bin/EDTA/EDTA.pl --genome cassava_tme7_phase1_pseudohap_contigs.fasta --cds Mesculenta_305_v6.1.cds.fa --rmout cassava_tme7_phase1_pseudohap_contigs.fasta.mod.out --curatedlib cassava_tme7_EDTA.TElib.fa --evaluate 1 --anno 1 -t 18 --step final &

To evaluate the contiguity of the repetitive sequence assembly, the LTR Assembly Index (LAI) was evaluated using LAI beta3.2 (Ou et al. 2018) with input files generated by EDTA. The initial LAI estimation was done using the ‘-q’ parameter, then average LTR identity and total LTR content were obtained and further provided to the standardization of LAI, with parameters ‘-iden 95.63 -totLTR 53’. Regional LAI was calculated in 3-MB windows with 300-kb overlapping steps.

# run LAI
nohup perl ~/las/git_bin/LTR_retriever/LAI -genome cassava_tme7_phase1_pseudohap_contigs.fasta.mod -intact cassava_tme7_phase1_pseudohap_contigs.fasta.mod.EDTA.raw/LTR/cassava_tme7_phase1_pseudohap_contigs.fasta.mod.pass.list -all cassava_tme7_phase1_pseudohap_contigs.fasta.mod.out -q -t 18 &
nohup perl ~/las/git_bin/LTR_retriever/LAI -genome cassava_tme7_phase0_scaffolded_renamed.fasta.mod -intact cassava_tme7_phase0_scaffolded_renamed.fasta.mod.EDTA.raw/LTR/cassava_tme7_phase0_scaffolded_renamed.fasta.mod.pass.list -all cassava_tme7_phase0_scaffolded_renamed.fasta.mod.out -q -t 18 &

# correct LAI
nohup perl ~/las/git_bin/LTR_retriever/LAI -genome cassava_tme7_phase1_pseudohap_contigs.fasta.mod -intact cassava_tme7_phase1_pseudohap_contigs.fasta.mod.EDTA.raw/LTR/cassava_tme7_phase1_pseudohap_contigs.fasta.mod.pass.list -all cassava_tme7_phase1_pseudohap_contigs.fasta.mod.out -q -t 18 -iden 95.63 -totLTR 53 &
nohup perl ~/las/git_bin/LTR_retriever/LAI -genome cassava_tme7_phase0_scaffolded_renamed.fasta.mod -intact cassava_tme7_phase0_scaffolded_renamed.fasta.mod.EDTA.raw/LTR/cassava_tme7_phase0_scaffolded_renamed.fasta.mod.pass.list -all cassava_tme7_phase0_scaffolded_renamed.fasta.mod.out -q -t 18 -iden 95.63 -totLTR 53 &


### Contig renaming scripts to take purge_dups output and convert to FALCON-Unzip naming scheme:

# rename_contigs.R

library(dplyr)
library(ggplot2)
library(reshape2)
library(tidyr)
library(knitr)
library(readr)

# round 2 haplotig index
# strip off "hap_" and index suffix from hap and make it ito column Contig

hfai = read_tsv("haplotigs.fa.fai", col_names = c("Hap","Length")) %>%
  mutate(Contig = sapply(strsplit(Hap, "hap_"), "[", 2),
         Contig =  substr(Contig, 0, nchar(Contig)))

# round 1 primary index (just in case)

pfai = read_tsv("primary.fa.fai", col_names = c("Primary","Length")) %>%
  arrange(Primary) %>%
  mutate(NewPrimary = paste0(stringr::str_pad(row_number()-1, 6, pad = 0), "F"))

# Round 1 bed file (soft linked from the manual cutoffs dir, i promise)

bed = read_tsv("dups.bed", col_names = c("Contig", "Start","Stop","Type","Primary")) %>%
  mutate(Length = Stop - Start)

# Haplotigs and OVLPS

HB = hfai %>%
  left_join(bed, by = "Contig")
  
haplotigs = HB %>%
  filter(Type == "HAPLOTIG") %>% 
  filter(Primary %in% pfai$Primary) %>%
  select(Hap, Primary) 

align = HB %>%
  filter(Type == "HAPLOTIG") %>% 
  filter(!Primary %in% pfai$Primary) %>%
  select(Hap)

OVLP = HB %>%
  filter(Type == "OVLP") %>%
  anti_join(haplotigs, by = "Hap")

caseA = OVLP %>%
  group_by(Contig) %>%
  filter(n() == 1) %>%
  ungroup()

easy = caseA %>%
  filter(!is.na(Primary)) %>%
  select(Hap, Primary)
  
align1 = caseA %>%
  filter(is.na(Primary)) %>%
  select(Hap)

caseb = OVLP %>%
  group_by(Contig) %>%
  filter(n() > 1) %>%
  ungroup()

align2 = caseb %>%
  group_by(Contig) %>%
  filter(sum(is.na(Primary)) > 0) %>%
  filter(is.na(Primary)) %>%
  ungroup()

maxLen = caseb %>%
  group_by(Contig) %>%
  filter(sum(is.na(Primary)) == 0) %>%
  filter(Length.y == max(Length.y)) %>%
  ungroup()


# REPEATS

repeats = HB %>%
  filter(Type == "REPEAT") %>%
  filter(Primary %in% pfai$Primary) 


special_case = HB %>%
  filter(Type == "REPEAT") %>%
  filter(!Primary %in% pfai$Primary)


# JUNK

HB %>%
  filter(Type == "JUNK") %>%
  anti_join(haplotigs, by = "Hap") %>%
  anti_join(OVLP, by = "Hap") %>%
  anti_join(filter(HB, Type == "REPEAT"), by = "Hap")

HB %>%
  filter(Type == "JUNK") %>%
  select(Contig) %>%
  left_join(bed)

junk = filter(HB, Type == "JUNK")


# HIGHCOV

HB %>%
  filter(Type == "HIGHCOV") %>%
  select(Contig) %>%
  left_join(HB) %>%
  count(Type)

highcov = filter(HB, Type == "HIGHCOV")

# Make alignment list
alignlist = rbind(select(align, Hap), select(align1, Hap), select(align2, Hap), select(special_case, Hap), select(junk, Hap), select(highcov, Hap))


# start renamelist

renamelist = rbind(select(haplotigs, Hap, Primary), select(easy, Hap, Primary), select(maxLen, Hap, Primary), select(repeats, Hap, Primary))

# write alignlist
write_csv(alignlist, "hapstoalign.txt", col_names = F)


#do nucmer alignment (bash)
xargs samtools faidx haplotigs.fa < hapstoalign.txt > hapstoalign.fa
nucmer -t 20 -p primary_hap2align primary.fa hapstoalign.fa && delta-filter -q primary_hap2align.delta > primary_hap2align.deltaq && show-coords -T primary_hap2align.deltaq > primary_hap2align.showcoords

# get nucmer alignment
alignment = read_tsv("primary_hap2align.showcoords", col_names = c("Start1","End1","Start2","End2","Len1","Len2","PID","Primary","Hap"), skip = 4)

matches = alignment %>%
  group_by(Hap, Primary) %>%
  mutate(TotalAlignLen = sum(Len1)) %>%
  group_by(Hap) %>%
  filter(TotalAlignLen == max(TotalAlignLen)) %>%
  filter(n() == 1) %>%
  ungroup()

alignment %>%
  group_by(Hap, Primary) %>%
  mutate(TotalAlignLen = sum(Len1)) %>%
  group_by(Hap) %>%
  filter(TotalAlignLen == max(TotalAlignLen)) %>%
  filter(n() > 1)


a = alignment %>%
  group_by(Hap, Primary) %>%
  mutate(TotalAlignLen = sum(Len1),
         MeanPID = mean(PID)) %>%
  group_by(Hap) %>%
  filter(TotalAlignLen == max(TotalAlignLen)) %>%
  filter(n() == 1)%>%
  select(Hap, Primary) %>%
  ungroup()

renamelist = rbind(renamelist, a)

b = alignment %>%
  group_by(Hap, Primary) %>%
  mutate(TotalAlignLen = sum(Len1),
         MeanPID = mean(PID)) %>%
  group_by(Hap) %>%
  filter(TotalAlignLen == max(TotalAlignLen)) %>%
  filter(n() > 1) %>%
  mutate(primarycount = n_distinct(Primary)) %>% 
  filter(primarycount == 1) %>%
  select(Hap, Primary) %>%
  unique() %>%
  ungroup()

renamelist = rbind(renamelist, b)


c = alignment %>%
  group_by(Hap, Primary) %>%
  mutate(TotalAlignLen = sum(Len1),
         MeanPID = mean(PID)) %>%
  group_by(Hap) %>%
  filter(TotalAlignLen == max(TotalAlignLen)) %>%
  filter(n() > 1) %>%
  mutate(primarycount = n_distinct(Primary)) %>%
  filter(primarycount > 1) %>%
  filter(Start1 == max(Start1)) %>%
  ungroup() %>%
  select(Hap, Primary)

renamelist = rbind(renamelist, c)

missing = anti_join(alignlist, alignment, by = "Hap")
missing %>% write_tsv("missing_from_alignment.txt", col_names = F)
  
# Blast results (bash)
bash, eval = F}
xargs samtools faidx haplotigs.fa < missing_from_alignment.txt > missing_from_alignment.fa
blastn -query missing_from_alignment.fa -subject primary.fa -outfmt 6 > missing_from_alignment_primary.blastn

blast = read_tsv("missing_from_alignment_primary.blastn", col_names = c("Hap","Primary","PID","Length","Mismatch","Gaps","HapStart","HapEnd","PrimaryStart","PrimaryEnd","Eval","Bit"))
blast

d = blast %>%
  group_by(Hap, Primary) %>%
  mutate(TotalAlignLen = sum(Length),
         MeanPID = mean(PID)) %>%
  group_by(Hap) %>%
  filter(TotalAlignLen == max(TotalAlignLen)) %>%
  filter(MeanPID == max(MeanPID)) %>%
  filter(n_distinct(Primary) == 1) %>% 
  ungroup() %>%
  select(Hap, Primary) %>%
  unique()

renamelist = rbind(renamelist, d) 

renamelist %>%
  arrange(Primary) %>%
  left_join(select(pfai, Primary, NewPrimary), by = "Primary") %>%
  group_by(Primary) %>%
  mutate(NewHap = paste0(NewPrimary, "_", stringr::str_pad(row_number(), 3, pad = 0))) %>%
  ungroup() %>%
  select(Hap, NewHap)

hap_rename = renamelist %>%
  arrange(Primary) %>%
  left_join(select(pfai, Primary, NewPrimary), by = "Primary") %>%
  group_by(Primary) %>%
  mutate(NewHap = paste0(NewPrimary, "_", stringr::str_pad(row_number(), 3, pad = 0))) %>%
  ungroup() %>%
  select(Hap, NewHap)

primary_rename = pfai %>%
  select(Primary, NewPrimary)

hap_rename %>%
  write_tsv("haplotigs_renaming.txt", col_names = F)
primary_rename %>%
  write_tsv("primary_rename.txt", col_names = F)

# rename_fasta.py
#!/usr/bin/env python3
from Bio import SeqIO
import sys

fasta = sys.argv[1]
subs = sys.argv[2]

rmap = {}
with open(subs) as f:
    for line in f:
        los = line.rstrip("\n").split("\t")
        rmap[los[0]] = los[1]


with open(fasta, "r") as handle:
    for record in SeqIO.parse(handle, "fasta"):
        newname = rmap[record.id]
        record.id = newname
        record.name = newname
        record.description = newname
        seq = str(record.seq)
        print(f">{newname}")
        print(f"{seq}")

## run the renaming script:
python rename_fasta.py haplotigs.fa haplotigs_renaming.txt > haplotigs_renamed.fa
python rename_fasta.py primary.fa primary_renaming.txt > primary_renamed.fa


### Modified version of coords2hp.py from FALCON-Phase that forces inclusion of entire haplotigs in to the phasing process

# Modified from: Sarah B. Kingan
# 20 September 2017
#
# Pacific Biosciences
# Applications Lab
#
# Convert nucmer.coords files for all haplotigs aligned
# to primary contigs into ncbi placement file
#
###########################################################################################################

# import libraries
import numpy as np
import argparse
import sys

###########################################################################################################


def main(argv=sys.argv):
    desc = '''Make haplotig placement file from individual coords file of all haplotigs to primary'''
    parser = argparse.ArgumentParser(description=desc)
    parser.add_argument("infile", help="coords file")
    args = parser.parse_args(argv[1:])

    run(args)


def gap(hd, j, k):
    if j <= k:
        return 0
    else:
        g = int(hd[j, 0]) - int(hd[k, 1])
        if g < 0:
            return 0
        else:
            return g


def run(args):
    # input file
    coords_file = args.infile

    # read file
    d = np.genfromtxt(open(coords_file, "rb"), dtype="str")
    # coords=[pstart_1 pend hstart_1 hend pAlnL hAlnL PID pL hL pID hID]

    # unique set of haplotigs
    haplotigs = list(set(list(d[:, 10])))
    # loop through haplotigs, subset array for each
    for h in haplotigs:
        output = ['hID', 'hL', 'hStart_0', 'hEnd', 'ori', 'pID',
                  'pL', 'pStart_0', 'pEnd', 'match', 'alnL', 'mapQ']

        # get data for a haplotig
        subset = []
        for i in range(0, d.shape[0]):
            if d[i, 10] == h:
                subset.append(i)
        n = len(subset)
        hd = d[subset]

        # initialize results array
        results = [{'start_index': -1, 'end_index': -1,
                    'score': float("-inf")} for l in range(n)]

        # score chained alignments
        for i in range(0, n):
            score = 0
            for j in range(i, n):
                score += int(hd[j, 4]) * float(hd[j, 6]) * \
                    0.01 - gap(hd, j, j-1)
                result_index = j - 1
                if results[result_index]['score'] < score:
                    results[result_index]['start_index'] = i
                    results[result_index]['end_index'] = j
                    results[result_index]['score'] = score

        # get best scoring chained alignment
        max = float("-inf")
        best = {'start_index': -1, 'end_index': -1, 'score': float("-inf")}
        for i in range(0, len(results)):
            if results[i]['score'] > max:
                best = results[i]
                max = best['score']

        # load data
        output[0] = hd[0, 10]  # htig name
        output[5] = hd[0, 9]  # pcontig name
        output[6] = hd[0, 7]  # pcontig length
        output[1] = hd[0, 8]  # hcontig length
        hstart = int(hd[best['start_index'], 2])  # hstart
        hend = int(hd[best['end_index'], 3])  # hend

        # To fit entire sequence inside region instead of cutting it down on both ends to fit, set the start to 0 and end to the full length if orietnation is forward, flip if reversed
        # Adam Boyher,, 2020
        if hstart < hend:
            hstart = 1
            hend = int(hd[0, 8])
        else:
            hstart = int(hd[0, 8])
            hend = 1
        pstart = int(hd[best['start_index'], 0])  # pstart
        pend = int(hd[best['end_index'], 1])  # pend

        # start and end coords
        output[2] = hstart-1  # hstart
        output[3] = hend  # hend
        output[7] = pstart-1  # pstart
        output[8] = pend  # pend

        # orientation
        output[4] = '+'  # ori
        hori = '+'
        pori = '+'
        if hstart > hend:
            hori = '-'
            output[2] = hend-1
            output[3] = hstart  # hend
        if pstart > pend:
            pori = '-'
            output[7] = pend-1  # pstart
            output[8] = pstart  # pend

        if hori != pori:
            output[4] = '-'

        # aln details, not real data
        output[10] = output[3]-output[2]  # alnL
        output[9] = output[10]  # match
        output[11] = '60'  # mapQ
        # print to stdout
        print('\t'.join(str(o) for o in output))


if __name__ == "__main__":
    main(sys.argv)


### Cassava linkage map tag extractor

# marker_seqs.py

#!/usr/bin/env python
# for use with cassavabase markers files to create sequences around marker SNPs 
# USAGE: 
​
import os, sys
import argparse
import pandas as pd
import re
​
def get_contigs(f):
    contigs = {}
    seqid = ''
    seq = ''
                    
    with open( f, 'r' ) as fh:
        for line in fh:
            if line[0] == '>':
                if seqid != '':
                    contigs[seqid] = seq
                    seq = ''
                seqid = line.split()[0][1:]
            else:
              seq += line.rstrip()
    
    contigs[seqid] = seq
​
    print('reference contigs: {}'.format(len(contigs.keys())) )
​
    return contigs
​
​
################
##### MAIN #####
################
def main( prog_name, argv ):
    # ARG PROCESSING
    parser = argparse.ArgumentParser( prog=prog_name, description='output a fasta file containing sequences around markers.. designed to be used with cassavabase marker files',
            formatter_class=argparse.ArgumentDefaultsHelpFormatter )
    parser.add_argument('-r','--radius', dest='radius', metavar='RADIUS', type=int, default=250, help='distance on either side of markers to pull sequence')
    parser.add_argument('-o','--outfile', dest='outfile', metavar='OUTFILE', required=True, type=str, help='file to print results to')
    parser.add_argument('-m','--marker_file', dest='marker_file', metavar='MARKER_FILE', required=True, type=str, help='file to read marker SNP locations from')
    parser.add_argument('-f','--ref_file', dest='ref_file', metavar='REF_FILE', required=True, type=str, help='file to read reference sequences from')
    args = parser.parse_args(argv)
​
    # SET VARIBLES FROM COMMAND LINE ARGS
    outfile = args.outfile
    marker_file = args.marker_file
    ref_file = args.ref_file
    radius = args.radius
​
​
    #### ADD CHECKS
    files_exist = True
    if not os.path.isfile( marker_file ):
        print( 'Error: {} does not exist'.format(marker_file) )
        files_exist = False
    if not os.path.isfile( ref_file ):
        print( 'Error: {} does not exist'.format(ref_file) )
        files_exist = False
​
    if not files_exist:
        sys.exit(1)
​
​
    # READ MARKERS
    mrkr_locs = pd.read_table(marker_file, sep='\t')
    mrkr_locs = mrkr_locs.drop_duplicates('marker')
    
    # Added position column to dataframe, makes it easier to see.
    mrkr_locs["pos"] = [x.split(":")[1] for x in mrkr_locs['marker']]
​
    # calculate ranges of marker seqs using radius as distance on either side of the SNP
    # mrkr_locs['range'] = mrkr_locs['marker'].str.split(':').apply(lambda t: t[0] + ':' + str( max(0,int(t[1])-radius) ) + '-' + str(int(t[1])+radius) )
    
    # Calculate start and end range using radius as distance on either side of the SNP, ignore boundaries of sequence for now. I'll deal with that later.
    mrkr_locs['start'] = pd.to_numeric(mrkr_locs['pos']) - radius
    mrkr_locs['end'] = pd.to_numeric(mrkr_locs['pos']) + radius
​
    # READ REFERENCE
    MEv4 = get_contigs(ref_file)
​
    # CREATE MARKER SEQUENCES
    # SGN_range = mrkr_locs.set_index('SGN id')['range']
​
    #Use the whole dataframe, no need to subset
    mrkr_locs = mrkr_locs.set_index('SGN id')
    mrkr_seqs = {}
    
    # for i in SGN_range.index.values:
    for i in mrkr_locs.index.values:
        # splt_rng = re.split(':|-', SGN_range[i])
​
        # get scaffold name by slicing marker column after 'scaffold' and zero filling the numerical identifier to 5 digits
        c = 'scaffold' + mrkr_locs['marker'][i][8:].split(":")[0].zfill(5)
​
        s = int(mrkr_locs['start'][i])
        e = int(mrkr_locs['end'][i])
        pos = int(mrkr_locs['pos'][i])
​
        #Now the fun part.
        # If s is less than 0, then the marker is closer to the start of the scaffold than the radius will allow. 
        # So let's set it to zero and modify the end point to match so we have equal radius on either side of marker.
        # Simply set the end point to the position * 2
        # Example:
        # Position = 3; radius = 5; Start = 0; End = 6; Sequence indices: 0,1,2 === 3 === 4,5,6
        if s < 0:
            s = 0
            e = pos*2
​
        # So, this is a good point to think about in the future if you care about where exactly the marker is in the extracted sequence.
        # The positions we're using are physical, so 1 based, and python indexing is 0 based. 
        # I'm not in the mood to figure it out exactly, but we're probably not compensating for the correctly.
        # But it isn't necessary to right now. So, we're just moving forward.
        # Also it's only an issue with the end bp. Or is it? (not be ominous i'm just not sure).
        
​
        # If e goes beyond the length of the scaffold, we need to modify the radius, just like for start.
        # Set e to the end of the scaffold.
        # Set start to the position minus the difference between the end point and the position plus 1.
        # Example: (keep in mind, when slicing, python is exclusive of the upper bound, meaning [0:3] gives 0, 1, 2 .... [0:3+1] gives 0, 1, 2, 3 )
        # Position = 300, end is 303; radius = 5; End = 303; Start = 300 - (303 - 300 + 1) = 300 - 4 = 296; Sequence indices: 296,297,298,299 === 300 === 301, 302, 303, 304
        # Still, this might not be right, but it's not important right now. So revisit it later when I'm thinking more logically.
        
        elif e > len(MEv4[c]):
            e = len(MEv4[c])
            s = pos - (e - pos + 1)
        try:
            mrkr_seqs[i] = MEv4[c][s:e+1]
        except:
            print( 'c: {}'.format(c) )
            print( 'i: {}'.format(i) )
            print( 's: {}'.format(s) )
            print( 'e: {}'.format(e) )
            print( 'keys: {}'.format( list(MEv4.keys())[:3] ) )
            sys.exit()
​
​
​
​
        # try:
        #     # split contig name, then adjust number formatting to follow the formatting of the reference contig names
        #     c = splt_rng[0].split('d')
        #     c = '{0}d{1:05d}'.format(c[0], int(c[1]))
            
        #     s = int(splt_rng[1])
        #     e = int(splt_rng[2])
            
        #     if s == 0:
        #         print(s,e,len(MEv4[c]))
        #     elif e > len(MEv4[c]):
        #         print(s,e,len(MEv4[c]))
        #     else:
        #         mrkr_seqs[i] = MEv4[c][s:e+1]
        # except:
        #     print( 'c: {}'.format(c) )
        #     print( 'i: {}'.format(i) )
        #     print( 's: {}'.format(s) )
        #     print( 'e: {}'.format(e) )
        #     print( 'keys: {}'.format( list(MEv4.keys())[:3] ) )
        #     sys.exit()
​
​
    # WRITE SEQUENCES TO FILE
    with open(outfile, 'w') as fh:
        for i in mrkr_seqs:
            fh.write( '>{}\n'.format(i) )
            fh.write( '{}\n'.format(mrkr_seqs[i]) )
​
​
    print( 'Exited Successfully' )
​
​
if __name__=='__main__':
    main(sys.argv[0], sys.argv[1:])

### Split scaffolds at greater than 10 N's for Assemblytics analysis

# split_scaffolds.py

import sys
import re
​
# fasta as arg1
fasta = sys.argv[1]
# generate n string by number of ns for gap (arg 2), example 5 ns ("nnnnn")
ns = "n" * int(sys.argv[2])
# generate regex match string, example 5 ns ("(NNNNN+|nnnnn+)")
split_string = f"({ns.upper()}+|{ns}+)"
​
# read each line of file
with open(fasta) as fn:
    scaffold = ''
    seq = ''
    start = 1
    end = 1
    line = fn.readline().rstrip('\n')
​
    # if sequence has gap of n length, split it
    while line:
        if line.startswith(">"):
            scaffold = line[1:]
            # start = 1
            end = 1
            line = fn.readline().rstrip('\n')
            contigs = re.split(split_string, line)
            for i in range(len(contigs)):
                c = contigs[i]
                if c == '':
                    continue
                start = end
                end = start + len(c)
                if c[0].upper() == "N":
                    continue
                else:
                    print(">" + scaffold + ":" +
                          str(start) + "-" + str(end - 1))
                    print(c)
            line = fn.readline().rstrip('\n')
