## Supplementary File 6 for "Large structural variations in the haplotype-resolved African cassava genome"

Figures and analyses scripts

Ben N. Mansfeld Adam Boyher Jeffrey C. Berry Mark Wilson Shujun Ou Seth Polydore Todd P. Michael Noah Fahlgren Rebecca S. Bart

6/23/2021

Load scripts and functions:

```
knitr::opts_chunk$set(echo = TRUE, cache = TRUE, warning=FALSE, message=FALSE)
library(tidyverse)
```

```
## -- Attaching packages --------------------------------------- tidyverse 1.3.1 --
## v ggplot2 3.3.4     v purrr   0.3.4
## v tibble  3.1.2     v dplyr   1.0.7
## v tidyr   1.1.3     v stringr 1.4.0
## v readr   1.4.0     v forcats 0.5.1
## -- Conflicts ------------------------------------------ tidyverse_conflicts() --
## x dplyr::filter() masks stats::filter()
## x dplyr::lag()    masks stats::lag()
```

```
library(UpSetR)

## Merqury color settings and functions
gray = "black"
red = "#E41A1C"
blue = "#377EB8" # light blue = "#56B4E9"
green = "#4DAF4A"
purple = "#984EA3"  # purple = "#CC79A7"
orange = "#FF7F00"  # orange = "#E69F00"
yellow = "#FFFF33"

merqury_col = c(gray, red, blue, green, purple, orange)
merqury_brw <- function(dat, direction=1) {
    merqury_colors=merqury_col[1:length(unique(dat))]
    if (direction == -1) {
        merqury_colors=rev(merqury_colors)
    }
    merqury_colors
}

ALPHA=0.4
LINE_SIZE=0.3

fancy_scientific <- function(d) {
```

```
    # turn in to character string in scientific notation
    d <- format(d, scientific = TRUE)
    # quote the part before the exponent to keep all the digits and turn the 'e+' into 10^ format
    d <- gsub("^(.*)e\\+", "'\\1'%*%10^", d)
    # convert 0x10^00 to 0
    d <- gsub("\\'0[\\.0]*\\'(.*)", "'0'", d)
    # return this as an expression
    parse(text=d)
}

format_theme <- function() {
    theme(legend.text = element_text(size=11),
          # legend.position = c(0.95,0.95),  # Modify this if the legend is covering your favorite circ
          legend.background = element_rect(size=0.1, linetype="solid", colour ="grey85"),
          legend.box.just = "right",
          legend.justification = c("right", "top"),
          axis.title=element_text(size=14,face="bold"),
          axis.text=element_text(size=12))
}

format_genomic <- function(...) {
      # Format a vector of numeric values according
      # to the International System of Units.
      # http://en.wikipedia.org/wiki/SI_prefix
      #
      # Based on code by Ben Tupper
      # https://stat.ethz.ch/pipermail/r-help/2012-January/299804.html
      # Args:
      #   ...: Args passed to format()
      #
      # Returns:
      #   A function to format a vector of strings using
      #   SI prefix notation
      #

      function(x) {
            limits <- c(1e0,   1e3, 1e6)
            #prefix <- c("","Kb","Mb")

            # Vector with array indices according to position in intervals
            i <- findInterval(abs(x), limits)

            # Set prefix to " " for very small values < 1e-24
            i <- ifelse(i==0, which(limits == 1e0), i)

            paste(format(round(x/limits[i], 1),
                         trim=TRUE, scientific=FALSE, ...)
                #  ,prefix[i]
            )
      }
}
```

### Main text

#### Figure 1

```
fc <-
  read_csv(
    "Files for Figures/FlowCyto_080216.csv",
    skip = 21,
    skip_empty_rows = T,
    col_names = c("Line", "ID", "G0+G1", "Std", "DNA_Content")
  ) %>%
  fill(Line, ID) %>%
  separate(Line, into = c("Line", "Rep"), sep = " ") %>%
  mutate(Line = ifelse(Line == "Oko-iyawo", "TME7", Line))

fc_fig <- fc %>%
  filter(Line == "TME7") %>%
  ggplot(aes(
    x = Rep,
    y = DNA_Content / 2 * 1e3,
    group = Rep,
    fill = as.factor(Rep)
  )) +
  geom_boxplot() +
  geom_jitter(color = "black", width = 0.25) +
  labs(x = "Sample", y = "Weight (Mb C-Value)") +
  cowplot::theme_cowplot() +
  theme(axis.text.x = element_text(angle = 30, hjust = 1)) +
  cowplot::panel_border() +
  theme(legend.position = "none")
```

```
gs_specta <- cowplot::ggdraw() +
    cowplot::draw_image(image = "Files for Figures/genomescope1.png")
```

Make Figure 1:

```
cowplot::plot_grid(fc_fig, gs_specta, align = "h", labels = "auto", rel_widths = c(3, 7))
```

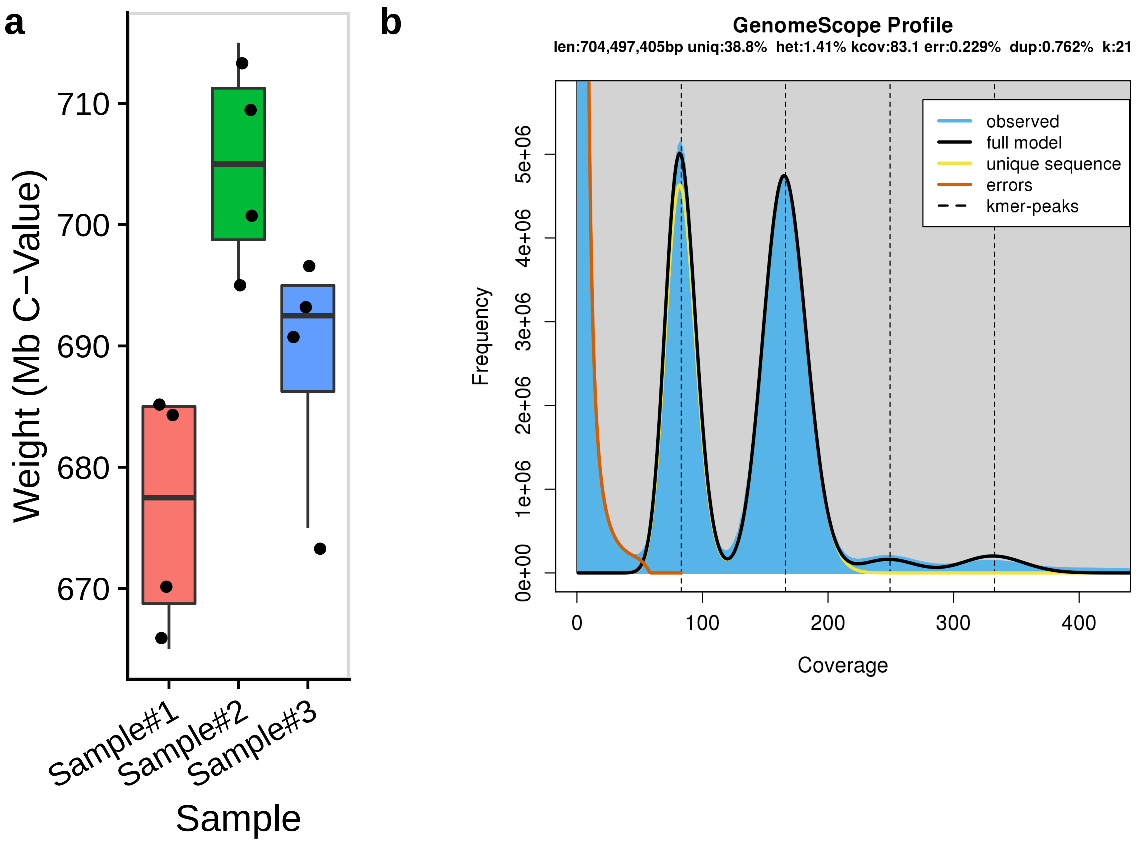


#### Figure 2

Load merqury data:

```
falcon_200703_sepcn <-
    bind_rows(
        "Primary" = read_tsv(
          file = "Files for Figures/merqury/200703_4000.p_ctg.spectra-cn.hist"),
        "Alternate" = read_tsv(
          file = "Files for Figures/merqury/200703_4000.a_ctg.spectra-cn.hist"),
        .id = "Phase"
    )

unzip_200703_sepcn <-
    bind_rows(
        "Primary" = read_tsv(
          file = "Files for Figures/merqury/tme7_200703_unzip.cns_p_ctg.spectra-cn.hist"),
        "Alternate" = read_tsv(
          file = "Files for Figures/merqury/tme7_200703_unzip.cns_h_ctg.spectra-cn.hist"),
        .id = "Phase"
    )

pilon_200703_sepcn <-
    bind_rows(
        "Primary" = read_tsv(
          file = "Files for Figures/merqury/tme7_200703_pilon.cns_p_ctg_pilon.spectra-cn.hist"),
```

```
        "Alternate" = read_tsv(
          file = "Files for Figures/merqury/tme7_200703_pilon.cns_h_ctg_pilon.spectra-cn.hist"),
        .id = "Phase"
    )


purgedFullSRA_200703_sepcn <-
    bind_rows(
        "Primary" = read_tsv(file = "Files for Figures/merqury/purgeFullSRA1/purge_full_sra_manual.purg
        "Alternate" = read_tsv(file = "Files for Figures/merqury/purgeFullSRA1/purge_full_sra_manual.pu
        .id = "Phase"
    )

purgedFullSRArnd2_200703_sepcn <-
    bind_rows(
        "Primary" = read_tsv(file = "Files for Figures/merqury/purgeFullSRA2/primary__pd_rnd2_short_ful
        "Alternate" = read_tsv(file = "Files for Figures/merqury/purgeFullSRA2/primary__pd_rnd2_short_f
        .id = "Phase"
    )

pseudo_200703_sepcn <-
    bind_rows(
        "Primary" = read_tsv(file = "Files for Figures/merqury/phase_pseudo/phased_pseudohap_minaln500.
        "Alternate" = read_tsv(file = "Files for Figures/merqury/phase_pseudo/phased_pseudohap_minaln50
        .id = "Phase"
    )

phaseUnzip_200703_sepcn <-
    bind_rows(
        "Primary" = read_tsv(file = "Files for Figures/merqury/phase_unzip/phased_unzip_minaln500.phase
        "Alternate" = read_tsv(file = "Files for Figures/merqury/phase_unzip/phased_unzip_minaln500.pha
        .id = "Phase"
    )



all_sepcn <- bind_rows(
    "falcon" = falcon_200703_sepcn,
    "unzip_200703" = unzip_200703_sepcn,
    "pilon_200703" = pilon_200703_sepcn,
    # "purgedFullSRC_200703" = purgedFullSRA_200703_sepcn,
    "AddSRC_200703" = purgedFullSRArnd2_200703_sepcn,
    "phaseUnzip_200703" = phaseUnzip_200703_sepcn,
    "phasePseudo_200703" = pseudo_200703_sepcn,
    .id = "Version") %>%
    mutate(Version = fct_inorder(Version)) %>%
    separate(Version, into = c("Step", "Run"), sep = "_", remove = FALSE) %>%
    mutate(Step = fct_relevel(Step, "falcon", "unzip", "pilon", "AddSRC", "phaseUnzip"))
```

```
falc_200703_cn <- read_tsv(
  file = "Files for Figures/merqury/200703_4000.spectra-cn.hist") %>%
    mutate(Copies = fct_relevel(Copies, "read-only"),
           Copies = fct_relevel(Copies, ">4", after = 5))
```

```
unzip_200703_cn <- read_tsv(
  file = "Files for Figures/merqury/unzip/tme7_200703_unzip.spectra-cn.hist") %>%
    mutate(Copies = fct_relevel(Copies, "read-only"),
           Copies = fct_relevel(Copies, ">4", after = 5))

pilon_200703_cn <- read_tsv(
  file = "Files for Figures/merqury/pilon/tme7_200703_pilon.spectra-cn.hist") %>%
    mutate(Copies = fct_relevel(Copies, "read-only"),
           Copies = fct_relevel(Copies, ">4", after = 5))

SRC_200703_cn <- read_tsv(
  file = "Files for Figures/merqury/pilon+SRC/cns_p_h_ctg_pilon_SRC_fullAssemb.cns_p_h_ctg_pilon_SRC.sp
    mutate(Copies = fct_relevel(Copies, "read-only"),
           Copies = fct_relevel(Copies, ">4", after = 5))

purge_200703_cn <- read_tsv(file = "Files for Figures/merqury/purgeFullSRA2/primary__pd_rnd2_short_full
    mutate(Copies = fct_relevel(Copies, "read-only"),
           Copies = fct_relevel(Copies, ">4", after = 5))

phaseUnzip_200703_cn <- read_tsv(file = "Files for Figures/merqury/phase_unzip/phased_unzip_minaln500.sp
    mutate(Copies = fct_relevel(Copies, "read-only"),
           Copies = fct_relevel(Copies, ">4", after = 5))

phasePseudo_200703_cn <- read_tsv(file = "Files for Figures/merqury/phase_pseudo/phased_pseudohap_minal
    mutate(Copies = fct_relevel(Copies, "read-only"),
           Copies = fct_relevel(Copies, ">4", after = 5))
```

```
all_spectra_cn <- bind_rows(
    "falcon" = falc_200703_cn,
    "unzip" = unzip_200703_cn,
    "pilon" = pilon_200703_cn,
    "SRC" = SRC_200703_cn,
    "purge_dups" = purge_200703_cn,
    "phaseUnzip" = phaseUnzip_200703_cn,
    "phasePseudo" = phasePseudo_200703_cn,
    # "Add_SRC" = sra_200703_cn,
    .id = "Version") %>%
    mutate(Version = fct_inorder(Version))
```

```
f200703_asm <-
    read_tsv("Files for Figures/merqury/200703_4000.spectra-asm.hist") %>%
    bind_rows(
        read_tsv(
            "Files for Figures/merqury/200703_4000.dist_only.hist",
            col_names = colnames(.)
        )
    ) %>%
    mutate(
        Assembly = case_when(
            Assembly == "a_ctg-only" ~ "Alternate-only",
            Assembly == "p_ctg-only" ~ "Primary-only",
            TRUE ~ Assembly
        )
    ) %>%
```

```
    mutate(Assembly = fct_relevel(
        Assembly,
        "shared",
        "Alternate-only",
        "Primary-only",
        "read-only"
    ))


u200703_asm <-
    read_tsv("Files for Figures/merqury/tme7_200703_unzip.spectra-asm.hist") %>%
    bind_rows(
        read_tsv(
            "Files for Figures/merqury/tme7_200703_unzip.dist_only.hist",
            col_names = colnames(.)
        )
    ) %>%
    mutate(
        Assembly = case_when(
            Assembly == "cns_h_ctg-only" ~ "Alternate-only",
            Assembly == "cns_p_ctg-only" ~ "Primary-only",
            TRUE ~ Assembly
        )
    ) %>%
    mutate(Assembly = fct_relevel(
        Assembly,
        "shared",
        "Alternate-only",
        "Primary-only",
        "read-only"
    ))


p200703_asm <-
    read_tsv("Files for Figures/merqury/tme7_200703_pilon.spectra-asm.hist") %>%
    bind_rows(
        read_tsv(
            "Files for Figures/merqury/tme7_200703_pilon.dist_only.hist",
            col_names = colnames(.)
        )
    ) %>%
    mutate(
        Assembly = case_when(
            Assembly == "cns_h_ctg_pilon-only" ~ "Alternate-only",
            Assembly == "cns_p_ctg_pilon-only" ~ "Primary-only",
            TRUE ~ Assembly
        )
    ) %>%
    mutate(Assembly = fct_relevel(
        Assembly,
        "shared",
        "Alternate-only",
        "Primary-only",
```

```
        "read-only"
    ))


purge200703_asm <-
    read_tsv("Files for Figures/merqury/purgeFullSRA1/purge_full_sra_manual.spectra-asm.hist") %>%
    bind_rows(
        read_tsv(
            "Files for Figures/merqury/purgeFullSRA1/purge_full_sra_manual.dist_only.hist",
            col_names = colnames(.)
        )
    ) %>%
    mutate(
        Assembly = case_when(
            Assembly == "purge_full_sra_manual_old.hap-only" ~ "Alternate-only",
            Assembly == "purge_full_sra_manual_old.purged-only" ~ "Primary-only",
            TRUE ~ Assembly
        )
    ) %>%
    mutate(Assembly = fct_relevel(
        Assembly,
        "shared",
        "Alternate-only",
        "Primary-only",
        "read-only"
    ))

purgedX2_200703_asm <-
    read_tsv(
        "Files for Figures/merqury/purgeFullSRA2/primary__pd_rnd2_short_full.spectra-asm.hist"
    ) %>%
    bind_rows(
        read_tsv(
            "Files for Figures/merqury/purgeFullSRA2/primary__pd_rnd2_short_full.dist_only.hist",
            col_names = colnames(.)
        )
    ) %>%
    mutate(
        Assembly = case_when(
            Assembly == "purged-only" ~ "Alternate-only",
            Assembly == "purge_full_sra_manual.purged-only" ~ "Primary-only",
            TRUE ~ Assembly
        )
    ) %>%
    mutate(Assembly = fct_relevel(
        Assembly,
        "shared",
        "Alternate-only",
        "Primary-only",
        "read-only"
    ))

phase_200703_asm <-
```

```r
    read_tsv("Files for Figures/merqury/phase_unzip/phased_unzip_minaln500.spectra-asm.hist") %>%
    bind_rows(
        read_tsv(
            "Files for Figures/merqury/phase_unzip/phased_unzip_minaln500.dist_only.hist",
            col_names = colnames(.)
        )
    ) %>%
    mutate(
        Assembly = case_when(
            Assembly == "phased.unzip.1-only" ~ "Alternate-only",
            Assembly == "phased.unzip.0-only" ~ "Primary-only",
            TRUE ~ Assembly
        )
    ) %>%
    mutate(Assembly = fct_relevel(
        Assembly,
        "shared",
        "Alternate-only",
        "Primary-only",
        "read-only"
    ))

pseudo_200703_asm <-
    read_tsv(
        "Files for Figures/merqury/phase_pseudo/phased_pseudohap_minaln500.spectra-asm.hist"
    ) %>%
    bind_rows(
        read_tsv(
            "Files for Figures/merqury/phase_pseudo/phased_pseudohap_minaln500.dist_only.hist",
            col_names = colnames(.)
        )
    ) %>%
    mutate(
        Assembly = case_when(
            Assembly == "phased.1-only" ~ "Alternate-only",
            Assembly == "phased.0-only" ~ "Primary-only",
            TRUE ~ Assembly
        )
    ) %>%
    mutate(Assembly = fct_relevel(
        Assembly,
        "shared",
        "Alternate-only",
        "Primary-only",
        "read-only"
    ))


all_spectra_asm <- bind_rows(
     "falcon_200703" = f200703_asm,
     "unzip_200703" = u200703_asm,
     "pilon_200703" = p200703_asm,
     #"purgedups_200703" = purge200703_asm,
```

```
     "purgedupsX2_200703" = purgedX2_200703_asm,
     "phaseUnzip_200703" = phase_200703_asm,
     "phasePseudo_200703" = pseudo_200703_asm,
    # "purgedups+SRA_200703" = purged_SRC_200703_asm,
    # "purgedupsfull_200703" = manualpurgefull_200703_asm,
    # "purgedupsfullX2_200703" = manualpurgeX2_200703_asm,
    .id = "Version") %>%
    mutate(Version = fct_inorder(Version)) %>%
    separate(Version, into = c("Step", "Run"), sep = "_", remove = FALSE) %>%
    mutate(Step = fct_relevel(Step, "falcon", "unzip", "pilon", "purgedupsX2", "phase"))
```

```
p1 <- all_sepcn %>%
    mutate(Copies = fct_relevel(Copies, "read-only"),
           Copies = fct_relevel(Copies, ">4", after = 5)) %>%
    filter(Step == "phaseUnzip") %>%
    ggplot(aes(x=kmer_multiplicity, y=Count, color=Copies)) +
    geom_line(size = 0.5) +
    scale_color_manual(values = merqury_brw(all_sepcn$Copies,direction = 1), name="Times in\nassembly")
    cowplot::theme_cowplot() +
    cowplot::panel_border() +
    format_theme() +
    scale_y_continuous(labels=fancy_scientific) +
    coord_cartesian(xlim=c(0, 430), ylim=c(0, 5e6)) +
    facet_grid(~ Phase) +
    theme(legend.position = c(0.95,0.95)) +
    labs(x = "k-mer multiplicity")


p2 <- all_spectra_cn %>%
    filter(Version == "phaseUnzip") %>%
    mutate(Copies = fct_rev(Copies)) %>%
    ggplot(aes(x=kmer_multiplicity, y=Count, color=Copies, fill=Copies)) +
    geom_area(alpha = 0.4) +
    scale_color_manual(values = merqury_brw(all_spectra_cn$Copies, direction=-1),
                       name="Times in\nassembly",
                       breaks=rev(levels(all_spectra_cn$Copies))) +
    scale_fill_manual(values = merqury_brw(all_spectra_cn$Copies, direction=-1),
                      name="Times in\nassembly",
                      breaks=rev(levels(all_spectra_cn$Copies))) +
    cowplot::theme_cowplot() +
    cowplot::panel_border() +
    format_theme() +
    scale_y_continuous(labels=fancy_scientific) +
    coord_cartesian(xlim=c(0, 430), ylim=c(0, 5e6)) +
    theme(legend.position = c(0.95,0.95)) +
    labs(x = "k-mer multiplicity")

p3 <- all_spectra_asm %>%
    filter(Step == "phaseUnzip") %>%
    filter(kmer_multiplicity > 0) %>%
ggplot(aes(x=kmer_multiplicity, y = Count, color=Assembly, fill=Assembly)) +
    geom_area(alpha = 0.4) +
    # geom_bar(data = all_spectra_asm %>% filter(kmer_multiplicity == 0), aes(x = 0),
    #          position="stack", stat="identity", show.legend = FALSE, width = 3, alpha = 0.4) +
```

```r
    scale_color_manual(values = merqury_brw(all_spectra_asm$Assembly, direction=-1), name="Phase specifi
    scale_fill_manual(values = merqury_brw(all_spectra_asm$Assembly, direction=-1), name="Phase specific
    cowplot::theme_cowplot() +
    cowplot::panel_border() +
    format_theme() +
    scale_y_continuous(labels=fancy_scientific) +
    coord_cartesian(xlim=c(0, 430), ylim=c(0, 5e6)) +
    theme(legend.position = c(0.95,0.95)) +
    labs(x = "k-mer multiplicity")

bottom <- cowplot::plot_grid(p2, p3, nrow = 1, labels = c("b", "c"))

pdf("fig2.pdf", width = 8, height = 8)
cowplot::plot_grid(p1, bottom,  nrow = 2, align = 'V', axis = 'l', labels = c("a", ""))
dev.off()
```

```
## pdf
##   2
```

```r
cowplot::plot_grid(p1, bottom,  nrow = 2, align = 'V', axis = 'l', labels = c("a", ""))
```

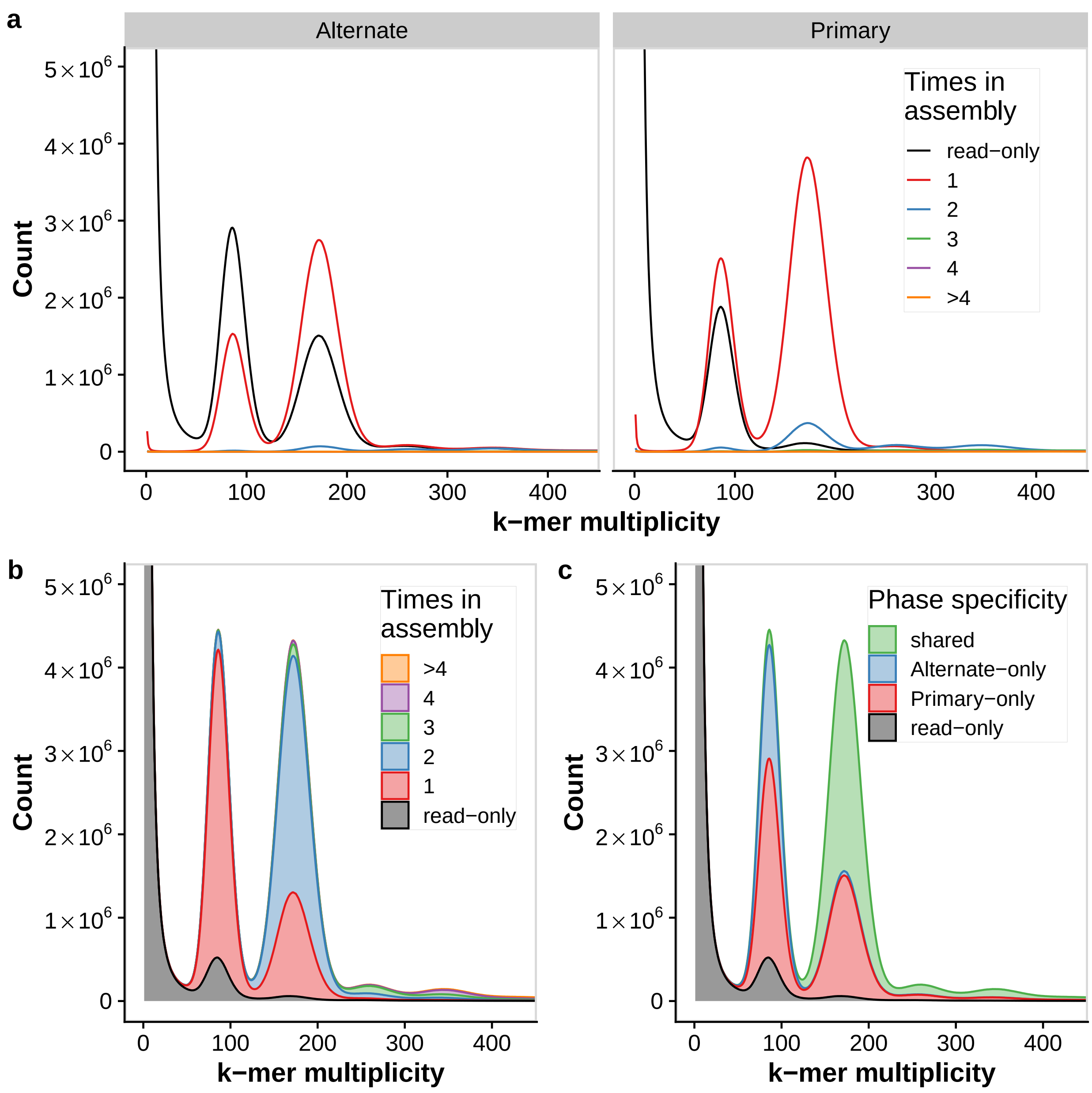


**Figure 3:**

```
hic <- read_tsv(file = "Files for Figures/hic/GInterations500.tsv",
                col_names = c("xchrom", "xstart", "xend", "ychrom", "ystart", "yend", "interactions"))

hic <- hic %>% filter(grepl("Chr", xchrom),
                      grepl("Chr", ychrom))


hic_chrm_lengths_x <-
    hic %>% group_by(xchrom) %>% summarise(length = max(xend)) %>%
    ungroup() %>%
    mutate(xpad = lag(length, default = 0),
```

```r
           xcumsumpad = cumsum(xpad),
           labelpos_x = xcumsumpad + length/2,
           label_x = paste0("Chr\n", str_extract(xchrom, "[0-9][0-9]")))

hic_chrm_lengths_y <-
    hic %>% group_by(ychrom) %>% summarise(length = max(yend)) %>%
    ungroup() %>%
    mutate(ypad = lag(length, default = 0),
           ycumsumpad = cumsum(ypad),
           labelpos_y = ycumsumpad + length/2,
           label_y = paste0("Chr\n", str_extract(ychrom, "[0-9][0-9]")))

hic_mat <- hic %>%
    left_join(hic_chrm_lengths_x, by = "xchrom") %>%
    left_join(hic_chrm_lengths_y, by = c("ychrom"))


 hic_plot <- hic_mat %>%
    ggplot() +
    geom_tile(
        aes(
            x = xstart + xcumsumpad,
            y = ystart + ycumsumpad,
            fill = log10(interactions)
        ),
        width = 500000,
        height = 500000
    ) +
    geom_tile(
        aes(
            y = xstart + xcumsumpad,
            x = ystart + ycumsumpad,
            fill = log10(interactions)
        ),
        width = 500000,
        height = 500000
    ) +
    geom_vline(data = hic_chrm_lengths_x,
               aes(xintercept = xcumsumpad),
               linetype = 2) +
    geom_hline(data = hic_chrm_lengths_x,
               aes(yintercept = xcumsumpad),
               linetype = 2) +
    geom_text(data = hic_chrm_lengths_x, aes(x = labelpos_x, y = -20e6, label = label_x)) +
    labs(x = "Phase0", y = "Phase0") +
    scale_fill_viridis_c() +
    coord_fixed() +
    cowplot::theme_cowplot() +
    theme(legend.position = c(0.75, 0.01))
```

```r
map <- read_tsv(file = "Files for Figures/marker_alignment/mesculenta_map_2014.txt") %>%
    mutate(CHROM = paste0(
        "Chromosome",
        stringr::str_pad(
```

```r
            gtools::roman2int(chromosome),
            width = 2,
            pad = "0"
        )
    )) %>%
    rename(LG = chromosome)


header <- c(
    "query_id",
    "ref_id",
    "perc_ident",
    "alignment_length",
    "mismatch",
    "num_gaps",
    "query_start",
    "query_end",
    "ref_start",
    "ref_end",
    "evalue",
    "bitscore"
)

blast_pri <-
    read_tsv(
      file = "Files for Figures/marker_alignment/cassava_tme7_phase0_scaffolded_renamed_2.linkage_map.bl[illegible]
             col_names = header) %>%
    # distinct(query_id, .keep_all = TRUE) %>%
    right_join(map, by = c("query_id" = "SGN id"))


blast_alt <- read_tsv(
  file = "Files for Figures/marker_alignment/p1_pseudohap_contigs.linkage_map_blast.txt",
                      col_names = header) %>%
    # distinct(query_id, .keep_all = TRUE) %>%
    right_join(map, by = c("query_id" = "SGN id"))


markers <- bind_rows("phase0_scaffolded" = blast_pri,
                     "phase1_contigs" = blast_alt,
                     .id = "phase")

markers_count <- markers %>%
    add_count(phase, query_id, .drop = FALSE, name = "nhits") %>%
    mutate(nhits = ifelse(nhits >= 10, 11, nhits),
           nhits = ifelse(is.na(ref_id), 0, nhits)) %>%
    mutate(qual =  (perc_ident >= 95 &
               alignment_length >= 150))

# filter qual
markers_count_filt <- markers_count %>%
    filter(qual) %>%
    add_count(phase, query_id, .drop = FALSE, name = "nhits_postfilter") %>%
```

```
    mutate(nhits_postfilter = ifelse(nhits_postfilter >= 10, 11, nhits_postfilter),
           nhits_postfilter = ifelse(is.na(ref_id), 0, nhits_postfilter))
    # mutate(ref_id = fct_relevel(ref_id, as.character(chrm_order) ))

# summarize post filter
markers_count_filt %>%
    group_by(phase, query_id) %>%
    summarise(count = unique(nhits_postfilter)) %>%
    group_by(phase, count) %>%
    summarize(nMarkers = n_distinct(query_id)) %>%
    group_by(phase) %>%
    mutate(percent = (nMarkers / sum(nMarkers)) * 100,
           count = ifelse(count == 11, "10+", count)) %>%
    pivot_wider(names_from = phase, values_from = c(nMarkers, percent)) %>%
    knitr::kable()
```

| count | nMarkers_phase0_scaffolded | nMarkers_phase1_contigs | percent_phase0_scaffolded | percent_phase1_contigs |
|---|---|---|---|---|
| 1 | 19252 | 19228 | 88.6290397 | 88.6409736 |
| 2 | 1941 | 1928 | 8.9356413 | 8.8880693 |
| 3 | 242 | 267 | 1.1140779 | 1.2308685 |
| 4 | 98 | 76 | 0.4511555 | 0.3503596 |
| 5 | 29 | 27 | 0.1335052 | 0.1244699 |
| 6 | 17 | 18 | 0.0782617 | 0.0829799 |
| 7 | 10 | 18 | 0.0460363 | 0.0829799 |
| 8 | 11 | 12 | 0.0506399 | 0.0553199 |
| 9 | 13 | 8 | 0.0598472 | 0.0368800 |
| 10+ | 109 | 110 | 0.5017954 | 0.5070994 |

```
# corr for each chrom
markers_count_filt %>%
    filter(phase == "phase0_scaffolded") %>%
    filter(nhits_postfilter < 10) %>%
    group_by(phase, ref_id) %>%
    summarise(cor(ref_start, position, method = "spearman"))
```

```
## # A tibble: 560 x 3
## # Groups:   phase [1]
##    phase            ref_id            `cor(ref_start, position, method = "spear~
##    <chr>            <chr>                                                  <dbl>
##  1 phase0_scaffold~ Chromosome01_Pha~                                      0.986
##  2 phase0_scaffold~ Chromosome02_Pha~                                      0.946
##  3 phase0_scaffold~ Chromosome03_Pha~                                      0.959
##  4 phase0_scaffold~ Chromosome04_Pha~                                      0.979
##  5 phase0_scaffold~ Chromosome05_Pha~                                      0.984
##  6 phase0_scaffold~ Chromosome06_Pha~                                      0.983
##  7 phase0_scaffold~ Chromosome07_Pha~                                      0.941
##  8 phase0_scaffold~ Chromosome08_Pha~                                      0.966
##  9 phase0_scaffold~ Chromosome09_Pha~                                      0.983
## 10 phase0_scaffold~ Chromosome10_Pha~                                      0.970
## # ... with 550 more rows
```

```
### all correlation
```

```
markers_count_filt %>%
    filter(phase == "phase0_scaffolded") %>%
    filter(nhits_postfilter < 10) %>%
    group_by(phase, ref_id) %>%
    summarise(correl = cor(ref_start, position, method = "spearman")) %>% filter(grepl(x = ref_id,patte
```

```
## # A tibble: 1 x 2
##   phase              `mean(correl)`
##   <chr>                       <dbl>
## 1 phase0_scaffolded           0.966
```

```
# plot
linkage_plot <- markers_count_filt %>%
    mutate(LG = fct_relevel(LG, as.character(as.roman(1:18)))) %>%
    filter(phase == "phase0_scaffolded") %>%
    filter(nhits_postfilter < 10) %>%
    filter(grepl("Chrom", ref_id)) %>%
    ggplot() +
    geom_point(aes(x = ref_start, y = position, color = LG)) +
    facet_grid(paste0("Chr", str_extract(ref_id, pattern = "[0-9][0-9]")) ~ ., scales = "free") +
    labs(x = "Physical position (Mb)", y = "Genetic distance (cM)") +
    cowplot::theme_cowplot() +
    cowplot::panel_border() +
    #theme(legend.position = "bottom", axis.text.x = element_text(angle = 45)) +
    guides(colour = guide_legend("Linkage group\nin map", ncol = 1)) +
    scale_x_continuous(labels = format_genomic()) +
    scale_color_viridis_d()
```

```
pdf("fig3.pdf", width = 14, height = 10)
cowplot::plot_grid(hic_plot + theme(legend.background = element_rect(fill = "white"), legend.position =
dev.off()
```

```
## pdf
##   2
```

#### Figure 4

```
all_buscos <- read_tsv("Files for Figures/busco/all_busco.tsv.txt", col_names = c("dir", "results")) %>
    separate(dir, into = "Version", sep = "/") %>%
    separate(results, into = c("Complete", "Duplicated", "Fragmented", "Missing", "n"), sep = ",") %>%
    separate(Complete, into = c("Complete", "Single"), sep = "\\[") %>%
    mutate(Complete =      as.numeric(gsub("[^0-9.-]", "", Complete)),
           Single =        as.numeric(gsub("[^0-9.-]", "", Single)),
           Duplicated =    as.numeric(gsub("[^0-9.-]", "", Duplicated)),
           Fragmented = as.numeric(gsub("[^0-9.-]", "", Fragmented)),
           Missing =       as.numeric(gsub("[^0-9.-]", "", Missing)),
           n =             as.numeric(gsub("[^0-9.-]", "", n))
           ) %>%
    mutate(Step = c(rep("Falcon", 3),
                    rep("Unzip", 3),
                    rep("Pilon", 3),
                    "Add SRC",
                    rep("Purge", 3),
                    rep("Phase_Unzip", 3),
```

```
                    rep("Phase_Pseudohap", 2),
                    rep("Phase0_scaffolded", 3)
                    ),
           Assembly = case_when(
               Version %in% c("BUSCO_2__a_ctg", "BUSCO_4__cns_h_ctg", "BUSCO_5__cns_h_ctg_pilon", "BUSCO
               Version %in% c("BUSCO_2__p_ctg", "BUSCO_4__cns_p_ctg", "BUSCO_5__cns_p_ctg_pilon", "BUSCO
               Version == "BUSCO_5__cns_h_p_ctg_pilon_SRA" ~ "Full",
               Version == "BUSCO_9__TME7_p0_p1_Unzip" ~ "Full (Unzip)",
               Version == "BUSCO_9__TME7_p0_p1_Pseudohap" ~ "Full (Pseudohap)",
               TRUE ~ "Full"
           )) %>%
    mutate(Step = fct_inorder(Step),
           Assembly = fct_relevel(Assembly, "Alternate", "Primary", "Full", "Full_Unzip", "Full_Pseudoha
    gather(-Version, -Step, -Assembly, key = "Category", value = "Value") %>%
    filter(Category != "n") %>%
    mutate(Number = ceiling(Value / 100 * 1614)) %>%
    mutate(label = paste0(round(Number / 1614 * 100, digits = 2), "%")) %>%
    mutate(Category = fct_rev(fct_relevel(Category, "Single", "Duplicated", "Fragmented", "Missing")))


#line just AP
AP <- all_buscos %>%
    filter(Category != "Complete") %>%
    filter(Assembly %in% c( "Alternate", "Primary")) %>%
    mutate(Assembly = case_when(grepl("Full", Assembly) ~ "Full",
                                TRUE ~ as.character(Assembly))) %>%
    ggplot(aes(x = Step, y = Number/1614*100, color = Assembly, group = Assembly)) +
    #geom_bar(stat = "identity", position = "stack") +
    geom_line() +
    geom_point(size = 2) +
    facet_grid(~ Category) +
    cowplot::theme_cowplot() +
    theme(axis.text.x = element_text(angle = 40, hjust = 1)) +
    labs(y = "Percent") +
    cowplot::panel_border()

# Just full assems
full <- all_buscos %>%
    filter(Category != "Complete") %>%
    filter(grepl("Full", Assembly)) %>%
    # mutate(Step = case_when(Assembly == "Full_SRC" ~ "Add SRC",
    #                          TRUE ~ as.character(Step))) %>%
    # filter(Assembly %in% c( "Alternate", "Primary")) %>%
    # mutate(Assembly = case_when(grepl("Full", Assembly) ~ "Full",
    #                              TRUE ~ as.character(Assembly))) %>%
    mutate(Step = case_when(
        Step == "Phase_Unzip" ~ "Falcon-Phase contigs + Unzip haplotigs",
        Assembly == "Full (Unzip)" ~ "Scaffolded + Unzip haplotigs",
        Assembly == "Full (Pseudohap)" ~ "Scaffolded + Pseuodhap haplotigs",
        TRUE ~ as.character(Step))
    ) %>%
    mutate(Step = fct_inorder(Step)) %>%
    ggplot(aes(x = Step, y = Number, color = Category, fill = Category, group = Category)) +
```

```r
    geom_bar(stat = "identity", position = "stack") +
    geom_text(aes(label = label), size = 3, color = "black", stat = "identity", position = position_stac
    # geom_line() +
    # geom_point() +
    # facet_grid(~ Category) +
    cowplot::theme_cowplot() +
    theme(axis.text.x = element_text(angle = 30, hjust = 1)) +
    # labs(y = "Percent") +
    cowplot::panel_border()
```

```r
phase0_busco <-
    read_tsv(
        file = "Files for Figures/busco/BUSCO_full_table_phase0_unzip.tsv",
        skip = 3,
        col_names = c(
            "Busco id",
            "Status",
            "Sequence",
            "Gene Start",
            "Gene End",
            "Score",
            "Length",
            "OrthoDB url",
            "Description"
        )
    ) %>%
    distinct(`Busco id`, .keep_all = TRUE)

phase1_busco <-
    read_tsv(
        file = "Files for Figures/busco/BUSCO_full_table_phase1_unzip.tsv",
        skip = 3,
        col_names = c(
            "Busco id",
            "Status",
            "Sequence",
            "Gene Start",
            "Gene End",
            "Score",
            "Length",
            "OrthoDB url",
            "Description"
        )
    ) %>%
    distinct(`Busco id`, .keep_all = TRUE)

full_buscos <- phase0_busco %>%
    bind_rows("phase0" = ., "phase1" = phase1_busco, .id = "phase") %>%
    mutate(set = paste(phase, Status, sep = "_")) %>%
    select(set, `Busco id`) %>%
    arrange(set) %>%
    mutate(i = 1) %>%
    spread(set, value = i, fill = 0) %>%
    select(contains("Comp"), contains("Dup"), contains("Frag"), contains("Miss"))
```

```
# svg(filename = "busco_ovlp.svg", width = 18, height = 6)
# upset(as.data.frame(full_buscos), sets = colnames(full_buscos), keep.order = T, mb.ratio = c(0.60, 0.
# dev.off()
```

```
busco_top <- cowplot::plot_grid(AP, full, nrow = 1, rel_widths = c(1, 0.6), align = "hv", axis = "b", la
pdf(file = "fig4.pdf", width = 16, height = 12)
cowplot::plot_grid(busco_top, cowplot::ggdraw() + cowplot::draw_image(image = "busco_ovlp.svg"), nrow =
dev.off()
```

```
## pdf
##   2
```

#### Figure 5

```
SV_p1 <-
    read_tsv(file = "Files for Figures/variation/cassava_tme7_phase1_unzip_contigs_vs_p0_scaffolds_spli
    separate(reference, into = c("Chrm", "coords"), sep = ":") %>%
    separate(
        coords,
        into = c("ctg_start", "ctg_end"),
        sep = "-",
        convert = TRUE
    ) %>%
    mutate(
        SV_start = ctg_start + ref_start,
        SV_end = ctg_start + ref_stop,
        SV_cmd = paste0(
            "-o ",
            Chrm,
            "_",
            SV_start,
            "_",
            SV_end,
            " -c ",
            Chrm,
            " -s ",
            SV_start,
            " -e ",
            SV_end
        )
    ) %>%
    mutate(query_coordinates = ifelse(
        str_count(query_coordinates, ":") == 2,
        sub(":", ":0-0:",
            query_coordinates),
        query_coordinates
    )) %>%
    separate(
        query_coordinates,
        into = c(
            "query_Chrm",
            "query_ctg_pos",
            "query_int_pos",
```

```
            "query_strand"
        ),
        sep = ":"
    ) %>%
    separate(
        col = query_ctg_pos,
        into = c("query_ctg_start", "query_ctg_end"),
        sep = "-",
        convert = T
    ) %>%
    separate(
        col = query_int_pos,
        into = c("query_int_start", "query_int_end"),
        sep = "-",
        convert = T
    ) %>%
    mutate(
        query_SV_start = query_ctg_start + query_int_start,
        query_SV_end = query_ctg_start + query_int_end
    )

write_delim(
    x = SV_p1 %>%  select(
        Chrm,
        SV_start,
        SV_end,
        everything(),
        -SV_cmd,
        -ctg_start,
        -ctg_end,
        -ref_start,
        -ref_stop,
        -contains("query_ctg"),
        -contains("query_int")
    ),
    "Supplementary_file_2_TME7_Phase1_vs_Phase0_SVs.tsv",
    delim = "\t"
)

SV_p1 %>% select(Chrm, SV_start, SV_end, type, size) %>%
    filter(type %in% c("Insertion", "Deletion"),!is.na(SV_start)) %>%
    arrange(Chrm, SV_start) %>% write_delim("SV_p1_indels.bed", delim = "\t", col_names = F)

SV_p1 %>% dplyr::select(Chrm, SV_start, SV_end, type, size) %>%
    filter(!is.na(SV_start)) %>% arrange(Chrm, SV_start) %>%
    write_delim("SV_p1_allSVs.bed", delim = "\t", col_names = F)
```

```
readDelta <- function(deltafile){
    lines = scan(deltafile, 'a', sep='\n', quiet=TRUE)
    lines = lines[-1]
    lines.l = strsplit(lines, ' ')
    lines.len = lapply(lines.l, length) %>% as.numeric
    lines.l = lines.l[lines.len != 1]
    lines.len = lines.len[lines.len != 1]
```

```r
    head.pos = which(lines.len == 4)
    head.id = rep(head.pos, c(head.pos[-1], length(lines.l)+1)-head.pos)
    mat = matrix(as.numeric(unlist(lines.l[lines.len==7])), 7)
    res = as.data.frame(t(mat[1:5,]))
    colnames(res) = c('rs','re','qs','qe','error')
    res$qid = unlist(lapply(lines.l[head.id[lines.len==7]], '[', 2))
    res$rid = unlist(lapply(lines.l[head.id[lines.len==7]], '[', 1)) %>% gsub('^>', '', .)
    res$strand = ifelse(res$qe-res$qs > 0, '+', '-')
    res
}

filterMum <- function(df, minl=1000, flanks=1e4){
    coord = df %>% filter(abs(re-rs)>minl) %>% group_by(qid, rid) %>%
        summarize(qsL=min(qs)-flanks, qeL=max(qe)+flanks, rs=median(rs)) %>%
        ungroup %>% arrange(desc(rs)) %>%
        mutate(qid=factor(qid, levels=unique(qid))) %>% select(-rs)
    merge(df, coord) %>% filter(qs>qsL, qe<qeL) %>%
        mutate(qid=factor(qid, levels=levels(coord$qid))) %>% select(-qsL, -qeL)
}

delta <- readDelta("Files for Figures/variation/cassava_tme7_phase1_unzips_p0_scaffolds.delta.filter")
delta_chr <- filter(delta, str_detect(rid, "Chr"), re-rs > 1e4)  #%>% arrange(rid, rs, qs) %>%
#rename(A = rid, B = qid, AStart = rs, AEnd =  re, BStart = qs, BEnd =  qe)

diagMum <- function(df){
    ## Find best qid order
    rid.o = df %>% group_by(qid, rid) %>% summarize(base=sum(abs(qe-qs)),
                                                    rs=weighted.mean(rs, abs(qe-qs))) %>%
        ungroup %>% arrange(desc(base)) %>% group_by(qid) %>% do(head(., 1)) %>%
        ungroup %>% arrange(desc(rid), desc(rs)) %>%
        mutate(qid=factor(qid, levels=unique(qid)))
    ## Find best qid strand
    major.strand = df %>% group_by(qid) %>%
        summarize(major.strand=ifelse(sum(sign(qe-qs)*abs(qe-qs))>0, '+', '-'),
                  maxQ=max(c(qe, qs)))
    merge(df, major.strand) %>% mutate(qs=ifelse(major.strand=='-', maxQ-qs, qs),
                                       qe=ifelse(major.strand=='-', maxQ-qe, qe),
                                       qid=factor(qid, levels=levels(rid.o$qid)))
}

delta_chr.diag <- diagMum(delta_chr) %>%
  mutate(rlab = paste0("Chr", str_extract(rid, "[0-9][0-9]"))) %>%
  mutate(similarity = 1 - error / abs(qe - qs)) %>%
  arrange(desc(as.numeric(qid)))

ctg_lengths <-
    delta_chr.diag %>% group_by(qid) %>% summarise(ctg_length = sum(abs(qe-qs))) %>%
    ungroup() %>%
    arrange(desc(as.numeric(qid))) %>%
    mutate(ypad = lag(ctg_length, default = 0),
           ycumsumpad = cumsum(ypad)
           )
```

```r
chrm_lengths <-
    delta_chr.diag %>% group_by(rid) %>% summarise(length = max(re)) %>%
    ungroup() %>%
    mutate(xpad = lag(length, default = 0),
           xcumsumpad = cumsum(xpad),
           labelpos = xcumsumpad + length/2,
           label = paste0("Chr", str_extract(rid, "[0-9][0-9]")))

delta_chr.diag <- delta_chr.diag %>%
    left_join(chrm_lengths, by = "rid") %>%
    left_join(ctg_lengths, by = "qid")

hapDotPlot <- delta_chr.diag %>%
  ggplot() +
  geom_point(
    data = filter(delta_chr.diag, similarity >= 0.98),
    aes(
      x = xcumsumpad + rs,
      y = ycumsumpad + qs,
      color = similarity,
      size = ctg_length
    ),
    alpha = 0.5
  ) +
  geom_point(
    data = filter(delta_chr.diag, between(similarity, 0.94, 0.98)),
    aes(
      x = xcumsumpad + rs,
      y = ycumsumpad + qs,
      color = similarity,
      size = ctg_length
    ),
    alpha = 0.5
  ) +
  geom_point(
    data = filter(delta_chr.diag, similarity <= 0.94),
    aes(
      x = xcumsumpad + rs,
      y = ycumsumpad + qs,
      color = similarity,
      size = ctg_length
    ),
    alpha = 0.5
  ) +
  geom_vline(data = chrm_lengths,
             aes(xintercept = xcumsumpad),
             linetype = 2) +
  geom_text(data = chrm_lengths, aes(x = labelpos, y = 1000, label = label)) +
  scale_size_continuous(name = "Haplotig\nlength") +
  labs(x = "Phase0 scaffolds", y = "Phase1 contigs") +
  scale_color_viridis_c() +
  cowplot::theme_cowplot()
```

```r
tme7_gff <-
    read_tsv(
        "Files for Figures/gffs/tme7_200703_falcon_phase0.gff",
        skip = 3,
        col_names = c(
            "seqid",
            "source",
            "type",
            "start",
            "end",
            "score",
            "strand",
            "phase",
            "attributes"
        ),
        comment = "#"
    )

tme7_TEs <-
    read_tsv(
        "Files for Figures/gffs/cassava_tme7_phase0_scaffolded_renamed.fasta.mod.EDTA.TEanno.gff3",
        skip = 3,
        col_names = c(
            "seqid",
            "source",
            "type",
            "start",
            "end",
            "score",
            "strand",
            "phase",
            "attributes"
        ),
        comment = "#"
    )

tme7_SVs <- SV_p1 %>% select(Chrm, SV_start, type) %>%
    rename(seqid = Chrm, start = SV_start) %>%
    mutate(seqid = str_remove(seqid, "omosome"),
           seqid = str_replace(seqid, "Phase", "P"))

genesTEs <-
    bind_rows(
        "Genes" = tme7_gff %>% filter(type == "gene"),
        "TE" = tme7_TEs,
        "SVs" = tme7_SVs,
        .id = "anno"
    )


annoDist <- genesTEs %>%
    filter(grepl("Chr", seqid)) %>%
    ggplot() +
```

```r
    geom_density(aes(x = start, y = after_stat(ndensity), fill = anno, color = anno), alpha = 0.6) +
    facet_wrap(~ seqid, ncol = 6, scales = "free_x") +
    cowplot::theme_cowplot() +
    theme(axis.text.x = element_text(angle = 30, hjust = 1)) +
    cowplot::panel_border() +
    labs(x = "Genomic position (Mb)", y = "Normalized density") +
    shades::lightness(scale_color_manual(values = c(viridisLite::viridis(4)[-4]),
                                         name="Feature"), shades::scalefac(0.6)) +
    scale_fill_manual(values = c(viridisLite::viridis(4)[-4]),
                      name="Feature") +
    scale_x_continuous(labels=format_genomic())

pdf("fig5.pdf", width = 12, 12)
cowplot::plot_grid(hapDotPlot, annoDist , ncol = 1, labels = "auto")
dev.off()
```

```
## pdf
##   2
```

#### Figure 6

In python MCScanX

#### Figure 7

```r
SV_Ref <-
    read_tsv(file = "Files for Figures/variation/Cassava_Phase0_renamed_split10_Ns_vs_esculenta_305_v6_
    separate(reference, into = c("Chrm", "coords"), sep = ":") %>%
    separate(
        coords,
        into = c("ctg_start", "ctg_end"),
        sep = "-",
        convert = TRUE
    ) %>%
    mutate(
        SV_start = ctg_start + ref_start,
        SV_end = ctg_start + ref_stop,
        SV_cmd = paste0(
            "-o ",
            Chrm,
            "_",
            SV_start,
            "_",
            SV_end,
            " -c ",
            Chrm,
            " -s ",
            SV_start,
            " -e ",
            SV_end
        )
    ) %>%
    mutate(query_coordinates = ifelse(
        str_count(query_coordinates, ":") == 2,
```

```r
        sub("Phase0:", "Phase0:0-0:",
            query_coordinates),
        query_coordinates
    )) %>%
    separate(
        query_coordinates,
        into = c(
            "query_Chrm",
            "query_ctg_pos",
            "query_int_pos",
            "query_strand"
        ),
        sep = ":"
    ) %>%
    separate(
        col = query_ctg_pos,
        into = c("query_ctg_start", "query_ctg_end"),
        sep = "-",
        convert = T
    ) %>%
    separate(
        col = query_int_pos,
        into = c("query_int_start", "query_int_end"),
        sep = "-",
        convert = T
    ) %>%
    mutate(
        query_SV_start = query_ctg_start + query_int_start,
        query_SV_end = query_ctg_start + query_int_end
    )


SV_Ref %>%
    filter(type == "Deletion") %>%
    arrange(desc(size)) %>% head
```

```
## # A tibble: 6 x 23
##   Chrm     ctg_start  ctg_end ref_start ref_stop ID           size strand type
##   <chr>        <int>    <int>     <dbl>    <dbl> <chr>       <dbl> <chr>  <chr>
## 1 Chromos~  11718858 11753535     23242    33116 Assemblyti~  9874 +      Delet~
## 2 Chromos~  19391747 19444245     18902    27886 Assemblyti~  8984 +      Delet~
## 3 Chromos~  26031509 26121103     50400    59066 Assemblyti~  8666 +      Delet~
## 4 Chromos~  22419269 22465670     32419    40418 Assemblyti~  7982 +      Delet~
## 5 Chromos~  14164795 14206982     23279    30937 Assemblyti~  7658 +      Delet~
## 6 Chromos~    543394   619976     54989    62492 Assemblyti~  7499 +      Delet~
## # ... with 14 more variables: ref_gap_size <dbl>, query_gap_size <dbl>,
## #   query_Chrm <chr>, query_ctg_start <int>, query_ctg_end <int>,
## #   query_int_start <int>, query_int_end <int>, query_strand <chr>,
## #   method <chr>, SV_start <dbl>, SV_end <dbl>, SV_cmd <chr>,
## #   query_SV_start <int>, query_SV_end <int>
```

```r
# SV_Ref %>% #filter(type != "Tandem_contraction") %>%
#     ggplot() +
#     geom_histogram(aes(x = size, fill = type), binwidth = 100) +
```

```
#     facet_wrap(~ str_replace(type, "_", " "),
#                ncol = 1, scales = "free_y") +
#     cowplot::theme_cowplot() +
#     theme(axis.text.x = element_text(angle = 30, hjust = 1)) +
#     cowplot::panel_border() +
#     labs(x = "Variant size (bp)", y = "Count") +
#     guides(fill = guide_none()) +
#     scale_x_log10()

write_delim(x = SV_Ref %>%  select(Chrm, SV_start, SV_end, everything(), -SV_cmd, -ctg_start, -ctg_end,

SV_Ref_dist_plot <- SV_Ref %>%
    mutate(bin = cut(size, breaks = c(0, 100, 500, 1000, 2500, 5000, 10000))) %>%
    group_by(type) %>%
    add_count(name = "total") %>%
    mutate(facet_label = paste0(str_replace(type, "_", " "), " (n=", total, ")")) %>%
    group_by(bin, facet_label) %>%
    count() %>%
    ggplot() +
    geom_bar(aes(x = bin, y = n, fill = facet_label), stat = "identity") +
    facet_wrap(~ facet_label,
               ncol = 1,
               scales = "free_y") +
    cowplot::theme_cowplot() +
    theme(axis.text.x = element_text(angle = 30, hjust = 1)) +
    cowplot::panel_border() +
    labs(x = "Variant size (bp)", y = "Count") +
    scale_x_discrete(labels = c("0-100", "100-500", "500-1000", "1000-2500", "2500-5000", "5000-10000")
    guides(fill = guide_none()) + scale_fill_viridis_d()


DELs <- SV_Ref %>% filter(type == "Deletion") %>%
    select(Chrm, SV_start, SV_end) %>%
    arrange(Chrm, SV_start) %>%
    rename()

#write_tsv(DELs, path = "dels.bed", col_names = FALSE)


gff <- read_tsv("Files for Figures/variation/Mesculenta_305_v6.1.gene.gff3", skip = 3, col_names = c("se

gff %>%
    filter(type == "gene") %>%
    arrange(seqid, start) %>%
    write_tsv(path = "genes.gff3", col_names = FALSE)

gene_dist <- gff %>%
    filter(type == "gene") %>%
    group_by(seqid, strand) %>% arrange(seqid, start) %>%
    mutate(distanceUp = case_when(
      strand == "+" ~ start - lag(end),
      strand == "-" ~ lead(start) - end
    ))
```

```
# From bedtools closest -D b
closest <- read_tsv(file = "Files for Figures/variation/closest.bed", col_names = FALSE)
gene_closest <- read_tsv("Files for Figures/variation/AM560genes_vs_TME7dels.bed", col_names = FALSE)


closest <- closest %>%
    mutate(WhereDel = ifelse(X13 < 0, "Upstream", "Downstream"), #this is correct because i used closes
        # WhereDel = ifelse((X10 == "+" & X13 < 0) | (X10 == "-" & X13 > 0), "Upstream", "Downstream"),
           WhereDel = fct_relevel(WhereDel, "Upstream"))

SV_distance_plot <- closest %>% filter(X13 != 0) %>% filter(abs(X13) <= 1e4) %>%
    ggplot() +
    geom_density(aes(x = X13)) + facet_grid(~ WhereDel, scales = "free") +
    labs(x = "Distance to nearest gene") +
    cowplot::theme_cowplot() +
    theme(axis.text.x = element_text(angle = 30, hjust = 1)) +
    cowplot::panel_border()  +
    scale_x_continuous(breaks = c(seq(0, 25000, by = 2000), seq(0, -25000, by = -2000)))
```

Chr03 Het SV dotplot

```
readDelta <- function(deltafile){
  lines = scan(deltafile, 'a', sep='\n', quiet=TRUE)
  lines = lines[-1]
  lines.l = strsplit(lines, ' ')
  lines.len = lapply(lines.l, length) %>% as.numeric
  lines.l = lines.l[lines.len != 1]
  lines.len = lines.len[lines.len != 1]
  head.pos = which(lines.len == 4)
  head.id = rep(head.pos, c(head.pos[-1], length(lines.l)+1)-head.pos)
  mat = matrix(as.numeric(unlist(lines.l[lines.len==7])), 7)
  res = as.data.frame(t(mat[1:5,]))
  colnames(res) = c('rs','re','qs','qe','error')
  res$qid = unlist(lapply(lines.l[head.id[lines.len==7]], '[', 2))
  res$rid = unlist(lapply(lines.l[head.id[lines.len==7]], '[', 1)) %>% gsub('^>', '', .)
  res$strand = ifelse(res$qe-res$qs > 0, '+', '-')
  res
}

mumgp <- readDelta("Files for Figures/variation/cassava_tme7_phase1_unzips_p0_scaffolds.delta")

filterMum <- function(df, minl=1000, flanks=1e4){
    coord = df %>% filter(abs(re-rs)>minl) %>% group_by(qid, rid) %>%
        summarize(qsL=min(qs)-flanks, qeL=max(qe)+flanks, rs=median(rs)) %>%
        ungroup %>% arrange(desc(rs)) %>%
        mutate(qid=factor(qid, levels=unique(qid))) %>% select(-rs)
    merge(df, coord) %>% filter(qs>qsL, qe<qeL) %>%
        mutate(qid=factor(qid, levels=levels(coord$qid))) %>% select(-qsL, -qeL)
}

mumgp.filt = filterMum(mumgp, minl=1e5)

Chr03_mumgp <- mumgp %>% filter(rid == "Chromosome03_Phase0", qid == "001856F_006")
```

```
Chr03_mumgp_plot <- Chr03_mumgp %>%
    ggplot(aes(
        x = rs,
        xend = re,
        y = qs,
        yend = qe
    )) +
    geom_segment() +  cowplot::theme_cowplot() + geom_hline(yintercept = 64969, linetype = 3) +
    geom_hline(yintercept = 72186, linetype = 3) +
    xlab('Chromosome03_Phase0') +
    ylab('Haplotig 001856F_006') +
    xlim(17200000, 17237181) +
    scale_y_continuous(limits = c(62000, 90000),
                       breaks = c(60000, 64969, 70000, 72186, 80000, 90000))
```

```
SV1 <- cowplot::ggdraw() +
  cowplot::draw_image(image = "Files for Figures/variation/Chromosome14_20000228_20004341.png")

SV2 <- cowplot::ggdraw() +
  cowplot::draw_image(image = "Files for Figures/variation/Chromosome03_14207739_14214954.png")

SV_right <- cowplot::plot_grid(SV1, SV2, ncol = 1, labels = c("c", "d"))

SV_top <- cowplot::plot_grid(SV_Ref_dist_plot, SV_right, ncol = 2, rel_widths = c(0.3, 1), labels = c("

SV_bot <- cowplot::plot_grid(SV_distance_plot, Chr03_mumgp_plot, rel_widths = c(6, 4), labels = c("b",

pdf("fig7.pdf", 14, 14)
cowplot::plot_grid(SV_top, SV_bot, ncol = 1, rel_heights = c(2.5, 1))
dev.off()
```

```
## pdf
##   2
```

#### Figure 8

```
ase <- read_tsv("Files for Figures/variation/phaser_leaf_basq10_phase0_gene_ase.txt") %>%
     filter(totalCount > 10) %>%
     mutate(geneid = str_remove(str_extract(name, "(?<=ID=)(.*)(?=Name)"), ";"))  %>%
     rowwise() %>%
      mutate(pval = binom.test(aCount, totalCount, p = 0.5)$p.value
             ) %>%
      ungroup() %>%
  mutate(qval = p.adjust(pval, method = "fdr"),
         a_ratio = aCount / totalCount) %>%
  select(contig, start, stop, geneid, everything())

dist_ase <- read_tsv(file = "Files for Figures/variation/phaser_genes_dist_to_INDELs_io_iu.bed", col_na

ase_analysis <- ase %>%
  left_join(dist_ase, by = c(
    "contig" = "Chrm",
    "start" = "start",
```

```
    "stop" = "end"
  )) %>%
  mutate(ASE_type = case_when(
    # qval < 0.05 &
    #   abs(log2_aFC) >= 4 ~ "Complete",
    (a_ratio >= 0.90 | a_ratio <= 0.10) & qval < 0.05 ~ "Complete",
    qval < 0.05 ~ "Partial",
    TRUE ~ "No ASE"
  )) %>%
  mutate(ASE_type = fct_relevel(ASE_type, "No ASE", "Partial", "Complete")) %>%
  filter(type != ".")

write_delim("Supplementary_file_3_ASE_and_SVs.tsv", x = ase_analysis, delim = "\t")

ase_analysis %>%
  group_by(ASE_type) %>%
  summarise(n(), mean(dist), median(dist), max(a_ratio), min(a_ratio))
```

```
## # A tibble: 3 x 6
##   ASE_type `n()` `mean(dist)` `median(dist)` `max(a_ratio)` `min(a_ratio)`
##   <fct>    <int>        <dbl>          <dbl>          <dbl>          <dbl>
## 1 No ASE    8798      593311.         281417          0.846          0.154
## 2 Partial   3451      579986.         293471          0.897          0.101
## 3 Complete   494      651028.         344818.         1              0
```

```
ase_summ <- ase_analysis %>%
  filter(dist <= 10000) %>%
  group_by(ASE_type) %>%
  summarise(n(), mean(dist), median(dist), max(a_ratio), min(a_ratio))

ase_summ
```

```
## # A tibble: 3 x 6
##   ASE_type `n()` `mean(dist)` `median(dist)` `max(a_ratio)` `min(a_ratio)`
##   <fct>    <int>        <dbl>          <dbl>          <dbl>          <dbl>
## 1 No ASE     640        3990.          3442.          0.818          0.167
## 2 Partial    264        4169.          4012.          0.894          0.104
## 3 Complete    26        3248.          3174           1              0
```

```
ase1 <- ase_analysis %>%
    ggplot() +
    geom_point(aes(x = aCount, y = bCount, color = ASE_type), alpha = 0.6) +
    scale_x_log10() +
    scale_y_log10() +
    cowplot::theme_cowplot() +
    scale_color_manual(values = c(viridisLite::viridis(4, direction = -1))[-1],
                       name="ASE tpe") +
    labs(x = "Reference read count", y = "Alternate read count")

# INDELS
ase2 <- ase_analysis %>%
  filter(dist <= 10000) %>%
  ggplot() +
  geom_density(aes(x = dist, fill = ASE_type), alpha = 0.6) +
    cowplot::theme_cowplot() +
```

```r
  labs(x = "Distance to nearest large InDel") +
  scale_fill_manual(values = c(viridisLite::viridis(4, direction = -1))[-1],
                    name="ASE type")

legend <- cowplot::get_legend(
  # create some space to the left of the legend
  ase2 + theme(legend.box.margin = margin(0, 0, 0, 12))
)
```

```r
pdf("fig8.pdf", width = 8, height = 4)
cowplot::plot_grid(ase1 + theme(legend.position="none"), ase2 , rel_widths = c(3, 4), ncol = 2, labels =
dev.off()
```

```
## pdf
##   2
```
